# Mural norrin/β-catenin signaling regulates Lama2 expression to promote neurovascular unit assembly

**DOI:** 10.1101/2022.02.18.481046

**Authors:** Saptarshi Biswas, Sanjid Shahriar, Nicholas P. Giangreco, Panos Arvanitis, Markus Winkler, Nicholas P. Tatonetti, William J. Brunken, Tyler Cutforth, Dritan Agalliu

**Affiliations:** Departments of Neurology, Columbia University Irving Medical Center, New York, NY 10032, USA; Pathology & Cell Biology, Columbia University Irving Medical Center, New York, NY 10032, USA; Systems Biology, Columbia University Irving Medical Center, New York, NY 10032, USA; Biomedical Informatics and Columbia University Irving Medical Center, New York, NY 10032, USA; Biomedical Engineering, Columbia University Irving Medical Center, New York, NY 10032, USA; Faculty of Medicine, Institute of Anatomy, Ludwig-Maximilians Universität, Munich 80336, Germany; Departments of Ophthalmology & Visual Sciences, SUNY Upstate Medical University, Syracuse, NY 13210, USA

## Abstract

Neurovascular unit (NVU) assembly and barrier maturation rely on vascular basement membrane (vBM) composition. Laminins, a major vBM component, are critical for these processes, yet which signaling pathway(s) regulate their expression remains unknown. Here we show that mural cells have active Norrin/β-catenin signaling during central nervous system development. Bulk RNA sequencing and validation using P10 and P14 wild-type versus *Apcdd1^-/-^* retinas reveal that Lama2 (Laminin-α2 chain) mRNA and protein levels are increased in mutant vasculature undergoing higher Norrin/β-catenin signaling. Mural cells are the main source of Lama2, and β-catenin activation induces Lama2 expression in mural cells *in vitro*. Markers of mature astrocytes including Aquaporin-4 (a water channel in astrocyte endfeet) and Integrin-α6 (a laminin receptor) are upregulated in *Apcdd1*^-/-^ retinas following higher Lama2 vBM deposition. Thus, the Norrin/β-catenin pathway regulates Lama2 expression in mural cells to promote NVU assembly and neurovascular barrier maturation.

**SUMMARY:** Biswas *et al*., demonstrate that Norrin/β-catenin signaling is active in CNS mural cells and regulates Lama2 deposition in the vascular basement membrane, promoting neurovascular unit assembly and blood-CNS barrier maturation.

## INTRODUCTION

In the developing central nervous system (CNS), reciprocal interactions among vascular components such as endothelial cells (ECs), mural cells [pericytes (PCs) and vascular smooth muscle cells (vSMCs)] and glial components (astrocytes in the brain; astrocytes and Müller cells in the retina) are critical to establish the neurovascular unit (NVU) (Biswas et al., 2020; Diaz-Coranguez et al., 2017; Iadecola, 2017). The NVU is essential for neurovascular coupling, regulation of blood flow and the establishment of CNS vascular barrier properties termed the blood-brain barrier (BBB) and the blood-retinal barrier (BRB), in the brain and retina respectively. The neurovascular barrier, consisting of tight junction proteins that restrict paracellular permeability between ECs, active or passive transporters regulating the transport of nutrients and the absence of caveolae to reduce transcytosis within ECs (Biswas et al., 2020; Diaz-Coranguez et al., 2017; Liebner et al., 2018), ensures a protective milieu for neuronal function (Iadecola, 2017; Liebner et al., 2018). Because NVU integrity and BBB/BRB properties are perturbed in many cerebrovascular and ocular diseases (Biswas et al., 2020), it is critical to elucidate the cellular and molecular mechanisms regulating their integrity.

Although the cell biological components of the neurovascular barrier reside within ECs, interactions with mural cells and astrocytes via the vascular basement membrane (vBM) are critical for BBB/BRB integrity (**Fig. 1A**) (Baeten and Akassoglou, 2011; Biswas et al., 2020). Laminins, heterotrimeric glycoproteins made of α-, β- and γ-chains (Durbeej, 2010; Patarroyo et al., 2002), are extracellular matrix (ECM) components of the vBM that play a critical role in NVU assembly and BBB/BRB properties. Global deletion of the gene encoding for the Laminin-α2 chain (*Lama2*) (Menezes et al., 2014), or astrocyte-specific deletion of the Laminin γ1 chain (Chen et al., 2013), cause BBB leakage, hemorrhagic stroke, deficits in pericyte differentiation and loss of astrocyte endfeet polarization around blood vessels – phenotypes that are associated with disorganization of endothelial tight junctions (Yao et al., 2014). Loss of Lama2 also reduces expression of α-smooth muscle actin (αSMA; a vSMC marker) and Pdgfrβ (a mural cell marker) in cerebral vessels (Menezes et al., 2014). Consistent with these phenotypes, neuronal- or glial-specific deletion of Dystroglycan, a receptor for laminins, also increases BBB permeability (Menezes et al., 2014). Elimination of astrocyte-derived Laminin-γ1 or global deletion of γ3 chains also decreases vSMC coverage of cerebral (Chen et al., 2013) and retinal (Biswas et al., 2018) arteries, respectively. The expression of several receptors such as Integrin-α2, -α6, -β2 and -β4 or Dystroglycan (Paulus et al., 1993), required for attachment of astrocyte endfeet to the vBM, is also regulated by laminins. Loss of either Laminin-α2 (Menezes et al., 2014) or -β2 chain (Gnanaguru et al., 2013) reduces expression of Integrin-β1 in astrocyte endfeet in the brain and delays astrocyte migration in the retina causing vascular defects and neurovascular barrier leakage. Although all NVU cell types produce laminins, the precise cellular sources of laminin chains remain controversial. Astrocytes have been traditionally thought to be the main source of Laminin-α2 in the vBM of cerebral blood vessels (Sixt et al., 2001); however recent studies have shown that mural cells are the major source of Laminin-α2 in brain blood vessels (Armulik et al., 2010; He et al., 2018; Menezes et al., 2014; Vanlandewijck et al., 2018). Although *Lama2*^-/-^ mice have a disrupted BBB integrity (Menezes et al., 2014), phenocopying mice with loss of Wnt/β-catenin function in ECs, the upstream signaling pathways that regulate Lama2 expression and secretion into the vBM are unknown.

**Figure 1.**
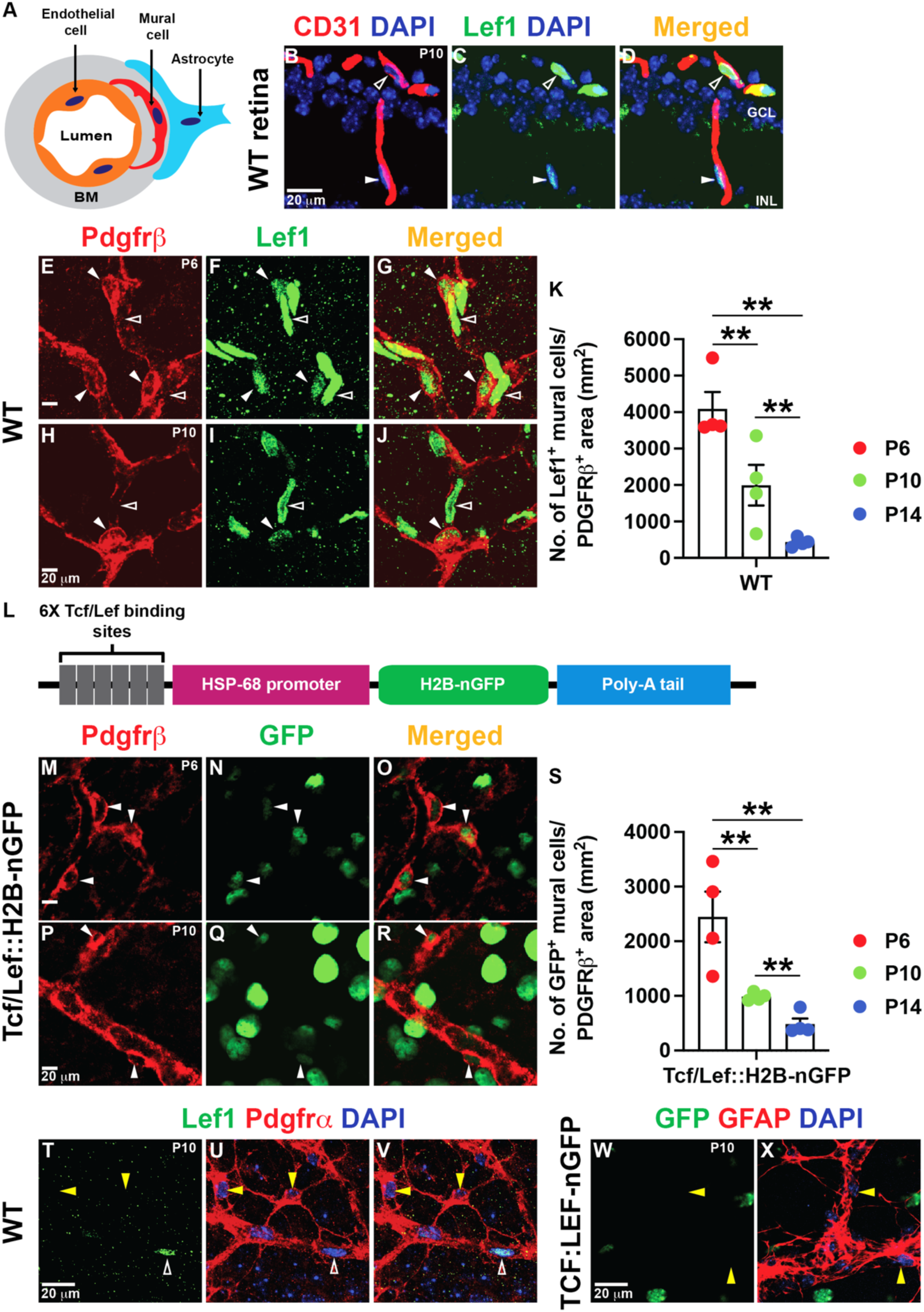
Retinal vascular mural cells have active Norrin/β-catenin signaling. **A)** Schematic diagram of the CNS cell types [endothelial cells (EC; orange), mural cells (red) and astrocytes (blue)] that form the neurovascular unit (NVU). The vascular basement membrane (BM) is depicted in gray. **B-D)** Postnatal (P) 10 wild-type (WT) retinal sections were labelled for Lef1 (Wnt/β-catenin component; green) and CD31 (EC marker; red). Empty arrowheads point to Lef1^+^ ECs and solid arrowhead point to Lef1^+^ cells next to the ECs in the ganglion cell layer (GCL) and inner nuclear layer (INL) of the retina. **E-J)** P6 and P10 WT retinal flat-mounts were labelled for Lef1 (green) and Pdgfrβ (red; mural cell marker). Empty arrowheads point to Lef1^+^ ECs and solid arrowheads point to Lef1^+^ mural cells. **K)** Dotter bar graph of Lef1^+^ mural cell numbers in the retina at three time-points (n=4/mice per time point). **L)** Schematic diagram of the *Tcf/Lef*::H2B-nGFP transgene. **M-R)** P6 and P10 *Tcf/Lef*::H2B-nGFP retinal flat-mounts were labelled for GFP (green) and Pdgfrβ (red). Solid arrowheads indicate GFP^+^ mural cells. **S)** Dotted bar graph of GFP^+^ mural cell numbers in the retina at indicated time-points (n=4/mice per time point). **T-V)** P10 WT retinal flat-mounts were labelled for Lef1 (green), Pdgfrα (astrocyte marker; red) and DAPI (blue). Yellow arrowheads point to Lef1^-^ astrocytes. **W, X)** P10 *Tcf/Lef*::H2B-nGFP retinal flat-mounts were labelled for GFP (green), GFAP (red, astrocyte marker) and DAPI (blue). Yellow arrowheads point to GFP^-^ astrocytes and empty arrowhead points to Lef1^+^ EC. **K, S**) For dotted bar graphs, the significance was determined by one-way ANOVA with Bonferroni corrections, **p<0.02. Graphs show mean +/- SEM. Scale bars: 20 μm.

Wnt/β-catenin signaling has emerged as a critical pathway for CNS angiogenesis and neurovascular barrier maturation. Genetic ablation of Wnt-7a, -7b ligands or downstream pathway components in ECs perturbs both brain angiogenesis and BBB integrity (Daneman et al., 2009; Liebner et al., 2008; Stenman et al., 2008; Zhou and Nathans, 2014; Zhou et al., 2014). The Wnt/β-catenin pathway also regulates retinal angiogenesis and blood-retina barrier (BRB) development through the Norrin-Fzd4-Lrp5/6 signaling module (Wang et al., 2012; Ye et al., 2009; Zhou et al., 2014). Genetic inactivation of either *Ndp* (encoding the Norrin ligand), *Fzd4*, *Lrp5/6* or *Tspan* receptors in ECs cause delayed growth of the superficial vascular plexus and loss of the deep vascular plexus in the retina, cerebellar hypovascularization and loss of the neurovascular barrier in both the retina and the cerebellum (Chen et al., 2012; Junge et al., 2009; Luhmann et al., 2005; Xia et al., 2010; Xu et al., 2004; Ye et al., 2009; Zuercher et al., 2012). We have shown that the Wnt inhibitor Apccd1 (Shimomura et al., 2010) is essential to coordinate vascular development, pruning and barrier maturation in the developing retina and cerebellum by modulating Wnt/Norrin signaling activity (Mazzoni et al., 2017). *Apcdd1*-deficient mice exhibit a transient increase in vessel density due to delayed vessel pruning, but a precocious maturation in the BBB/BRB, suggesting a critical role for this protein in neurovascular barrier formation (Mazzoni et al., 2017). Endothelial Wnt/β-catenin signaling promotes BBB/BRB maturation primarily by regulating expression of tight junction proteins, suppression of transcytosis and expression of a subset of transporters (e.g. Slc2a1) (Biswas et al., 2020). Recent studies have also implicated endothelial Wnt/β-catenin signaling in regulation of expression and deposition of Collagen IV, a critical vBM protein for neurovascular function, due to the activity of Fgfbp1 (Cottarelli et al., 2020). Although the roles of Wnt/β-catenin function in ECs for both CNS angiogenesis and BBB/BRB maturation have been well-characterized, it is unknown whether: a) this pathway is active in CNS mural cells, and b) how its activity in mural cells affects vBM composition, interactions with ECs and astrocytes and BBB/BRB development.

In this study, we have identified that mural cells have active Wnt/β-catenin signaling during CNS development, albeit at levels lower than ECs. Bulk RNA sequencing comparison and validation in P10 and P14 wild-type versus *Apcdd1^-/-^* retinas, which have higher Wnt/β-catenin signaling in the neurovasculature (Mazzoni et al., 2017), show that Lama2 mRNA and protein are upregulated in the vBM of *Apcdd1^-/-^* retina. Consistent with mural cells being the main source of Lama2 *in vivo*, we demonstrate that Wnt/β-catenin signaling activation induces *Lama2* expression in retinal mural cells *in vivo* and brain-derived mural cells *in vitro*. Finally, physiologic effectors of astrocytic homeostasis, including Integrin-α6 (a laminin receptor) and Aquaporin-4 (a water channel localized in polarized astrocyte endfeet), are upregulated in *Apcdd1*^-/-^ retina blood vessels. Our findings reveal that Wnt/β-catenin activation in mural cells regulates Lama2 expression to promote NVU assembly as well as astrocyte and neurovascular barrier maturation.

## RESULTS

### Retinal and cerebellar mural cells have active Norrin/β-catenin signaling

While Norrin (Wnt)/β-catenin signaling is well-characterized in retinal ECs, its role in other NVU cells is currently unknown. To determine whether Norrin/β-catenin signaling is active in NVU cells other than ECs during retina development, we examined the expression of Lef1, a downstream target and readout of pathway activation (Fancy et al., 2014). Lef1 was expressed at high levels in wild-type (WT) retinal ECs (**Fig. 1B-J; empty arrowheads**) and at lower levels in Pdgfrβ^+^ mural cells (Pdgfrβ^+^ is a mural cell marker; **Fig. 1B-J; solid arrowheads**) in the developing retina. The number of Lef1^+^ cells within the Pdgfrβ^+^ area covered by mural cells in flat mount retinas decreased significantly from P6 to P14 as the BRB matures (see Materials and Methods for quantification details; **Fig. 1K**) (Mazzoni et al., 2017). We confirmed these findings in the *Tcf/Lef*::H2B-nGFP transgenic mice, where nuclear GFP expression is under the control of the Wnt/β-catenin activity (**Fig. 1L**) (Ferrer-Vaquer et al., 2010; Lengfeld et al., 2017). Consistent with Lef1 expression, nuclear GFP was present in Pdgfrβ^+^ retinal mural cells (**Fig. 1M-R; solid arrowheads**), and their number decreased sharply with the maturation of the retinal vasculature (**Fig. 1S**). In contrast, Lef1 (WT) and GFP (*Tcf/Lef*::H2B-nGFP) were not expressed by astrocytes in the retina (**Fig. 1T-X; yellow arrowheads**), suggesting no active Norrin/β-catenin signaling. Nuclear GFP was also present in neurons of the *Tcf/Lef*::H2B-nGFP retinas (**Fig. 1M-R**), consistent with published single-cell RNA sequencing data from P14 retina (https://singlecell.broadinstitute.org/single_cell/study/SCP301/c57b6-wild-type-p14-retina-by-drop-seq#study-summary) (Macosko et al., 2015). Finally, we compared expression of Apcdd1, a downstream target and negative regulator of the Norrin/β-catenin signaling (Mazzoni et al., 2017; Shimomura et al., 2010) in endothelial versus mural cells in the retina. Double fluorescent RNA *in situ* hybridization with antisense probes against *Pdgfrb* and *Apcdd1* along with immunostaining for Caveolin-1 (EC marker) demonstrated that almost all (>95%) Caveolin-1^+^ ECs (**Fig. 2A-C, G; open arrowhead**) and a subset (∼20%) of *Pdgfrb*^+^ mural cells (**Fig.2D-G; solid arrowhead**) express *Apcdd1* mRNA in P12 retinal sections, confirming active Norrin/β-catenin signaling (Mazzoni et al., 2017). Similarly, analysis of P14 retinal single-cell RNA seq database (Macosko et al., 2015) confirmed that three Norrin/β-catenin targets (*Lef1*, *Apcdd1*, *Axin2*) are expressed at high levels in ECs and lower levels in mural cells, but are absent in astrocytes (**Sup. Fig. 1A**).

**Figure 2.**
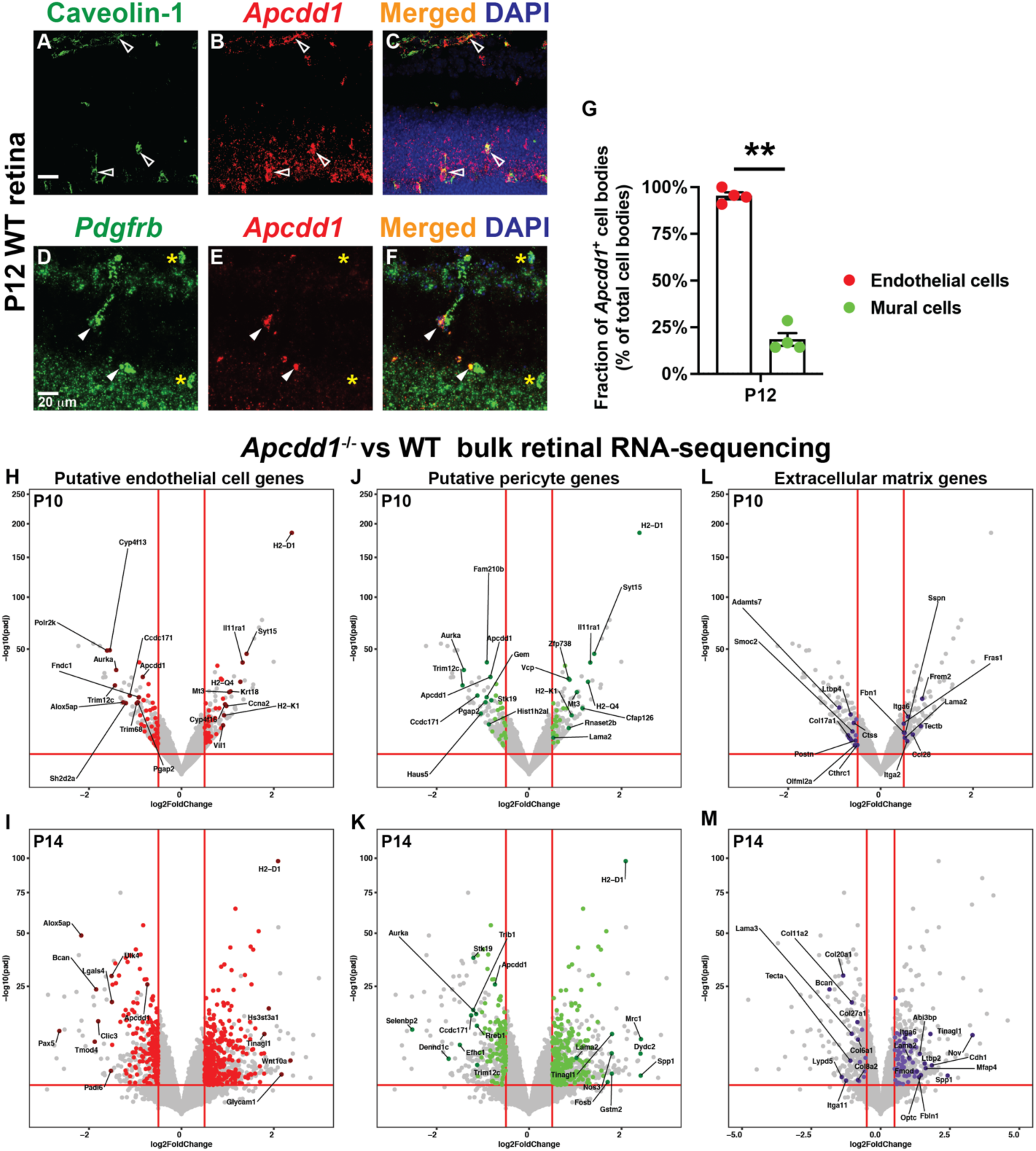
*Apcdd1* is expressed in both retinal ECs and PCs and *Lama2* expression is increased in *Apcdd1^-/-^* retinas. **A-C)** Fluorescence *in situ* hybridization of P12 wild-type (WT) retinal section with antisense probe against *Apcdd1* (red) followed by immunostaining for Caveolin-1 (EC marker; green). Empty arrowheads point to *Apcdd1*^+^ ECs. **D-F)** Double fluorescent *in situ* hybridization of P12 WT retinal section with antisense probes against *Apcdd1* (red) and *Pdgfr*β (green). Solid arrowheads point to *Apcdd1*^+^ mural cells and yellow asterisks point to *Apcdd1*^-^ mural cells. **G)** Dotted bar graph of the fraction of *Apcdd1*^+^ ECs and mural cells in the retina (n=4 mice; **p<0.02; Student’s t-test; Bars show mean +/- SEM.). **H-M)** Volcano plots showing genes assigned to P10 and P14 ECs (**H, I: red dots**), pericytes (**J, K: green dots**) and extracellular matrix (**L, M; purple dots**) that are differentially expressed between *Apcdd1*^-/-^ and WT retinas. Genes above horizontal red lines (Y axis) are significantly changed in their expression. Genes to the right of right vertical red lines (X-axis) are significantly upregulated and genes to the left of left vertical red lines (X-axis) are significantly downregulated in *Apcdd1*^-/-^ retinas. The top up and downregulated genes are named in the plots. Scale bars: 20 μm.

Norrin is critical for angiogenesis and barrier maturation, not only in the retina, but also cerebellum, during postnatal development (Wang et al., 2018; Wang et al., 2012; Zhou et al., 2014). Similar to the retina, developing P3 cerebellar mural cells, labeled with Foxc1 (Siegenthaler et al., 2013), had low Lef1 levels (Foxc1^+^ Lef1^low^; **Sup. Fig. 1B-C’, solid arrowhead**), in contrast to Foxc1^-^ Lef1^high^ ECs (**Sup. Fig. 1B-C’: empty arrowhead**). P10 cerebellar Pdgfrβ^+^ mural cells had also Lef1^Low^ expression (**Sup. Fig. 1D-E’; solid arrowhead**) next to Pdgfrβ^-^ Lef1^high^ ECs (**Sup. Fig. 1D-E’: empty arrowhead**). We found similar results in P10 and P14 *Tcf/Lef*::H2B-nGFP cerebella where Pdgfrβ^+^ mural cells expressed low nGFP (**Sup. Fig. 1F-I’; solid arrowheads**) adjacent to ECs with high nGFP (**Sup. Fig. 1F-I’; empty arrowheads**). In summary, mural cells, in addition to ECs, have active Norrin/β-catenin signaling in the retina and cerebellum, albeit at lower levels than ECs. Similar to ECs, Norrin/β-catenin activity decreases gradually in mural cells as the BRB matures during the retinal development.

### Lama2, expressed by mural cells, is upregulated in the *Apcdd1^-/-^* retinal vBM

We have previously shown that global deletion of the Wnt inhibitor Apcdd1 (*Apcdd1*^-/-^) leads to precocious maturation of the paracellular neurovascular barrier in the retina and cerebellum between P10 - P14 due to upregulation of Wnt/β-catenin activity (Mazzoni et al., 2017). Since Apcdd1 is expressed by both ECs and PCs, to understand how Apcdd1 regulates neurovascular barrier maturation, we performed a bulk mRNA sequencing and computation analysis between WT and *Apcdd1*^-/-^ retinas at two developmental stages (P10 and P14) to identify differentially expressed transcripts and assign them to specific NVU cell types. Briefly, total cellular mRNAs were extracted from WT and *Apcdd1*^-/-^ retinas, cDNA libraries were prepared and sequenced to generate read-counts for individual RNA species. Differential expression analyses were performed for these mRNAs using the Bioconductor package DESeq2 in R. Cutoffs of the Benjamini-Hochberg adjusted p values (padj) ≤ 0.05 and |log_2_ fold change| ≥ 0.5 were used to identify statistically significant, differentially expressed genes. Gene sets from the Molecular Signatures Database v6.1 (*MSigDB: software.broadinstitute.org /gsea/msigdb/genesets.jsp*) were used to curate comprehensive gene catalogs for various cellular and molecular pathways and functions. Putative retinal EC or mural cell sources for Apcdd1-regulated transcripts were assigned using the P14 retinal single cell RNAseq database (Macosko et al., 2015). Finally, differentially-expressed gene lists and volcano plots were constructed for each cell type and pathway category (**Fig. 2H-M**; **Sup. data sheet 1-5**). This computational analysis revealed that several EC-specific genes were differentially expressed between WT and *Apcdd1*^-/-^ retinas at both P10 and P14 (**Fig. 2H, I; Sup. data sheet 1, 2**) since *Apcdd1* mRNA is expressed at high levels in retinal ECs (Mazzoni et al., 2017). Interestingly, many pericyte-specific genes were also differentially expressed between WT and *Apcdd1*^-/-^ retinas at both developmental stages (**Fig. 2J, K; Sup. data sheet 1, 3**), consistent with *Apcdd1* mRNA expression in a subset of mural cells (**Fig. 2D-G**).

Among extracellular matrix (ECM) genes, *Lama2* (Laminin-α2) and *Itga6* (Integrin α6) transcripts were upregulated consistently at both P10 and P14 *Apcdd1*^-/-^ retinas (**Figs. 2L-M, Sup. data sheet 1, 3, 4**). In contrast, other ECM mRNAs such as *Lama1*, *Lama4* or *Lama5* (coding for Laminin-α1, Laminin-α4 and -α5 chains), or *Col4a1, Col4a2* (coding for Collagen 4 chains), respectively, were not significantly different between the two genotypes (**Sup. data sheet 1, 4**). Since *Apcdd1* is expressed by both ECs and mural cells, we examined which cell type produces *Lama2* mRNA. We immunolabelled P14 *Lama2^Lacz^*^/+^ reporter retinas, where *LacZ* encoding for β- Galactosidase is inserted into the *Lama2* locus (Kuang et al., 1998), with markers for ECs (Lectin), mural cells (NG2; αSMA^+^) and astrocytes (GFAP). Vascular β-Galactosidase colocalized with NG2^+^ mural cells and αSMA^+^ vSMCs (**Fig. 3A-B’; solid arrowheads**), but not Lectin^+^ ECs (**Fig. 3C, C’; empty arrowheads**). The astrocyte marker GFAP also partially colocalized with vascular β-Galactosidase (**Fig. 3D, D’; solid arrowheads**). Using the P14 retinal single-cell RNA sequencing database (Macosko et al., 2015), we confirmed that mural cells, specifically pericytes, are the predominant source of *Lama2* in the P14 retina (**Sup. Fig. 2A**). Consistent with these data, fluorescent *in situ* hybridization in P14 retina sections showed that *Lama2* mRNA is localized to mural cells (**Sup. Fig. 2B: solid arrowheads**), but not ECs (**Sup. Fig. 2C: empty arrowheads**). Similar to the retina, β-Galactosidase was also localized with NG2^+^ mural cells (**Sup. Fig. 3D-F; solid arrowheads**), but not Lectin^+^ ECs (**Sup. Fig. 3A-C; empty arrowheads**), in cerebellar blood vessels of the *Lama2^Lacz^*^/+^ reporter mice, a finding consistent with brain single cell RNA sequencing studies showing that mural, but not endothelial, cells express *Lama2* (Armulik et al., 2010; He et al., 2018; Menezes et al., 2014; Vanlandewijck et al., 2018).

**Figure 3.**
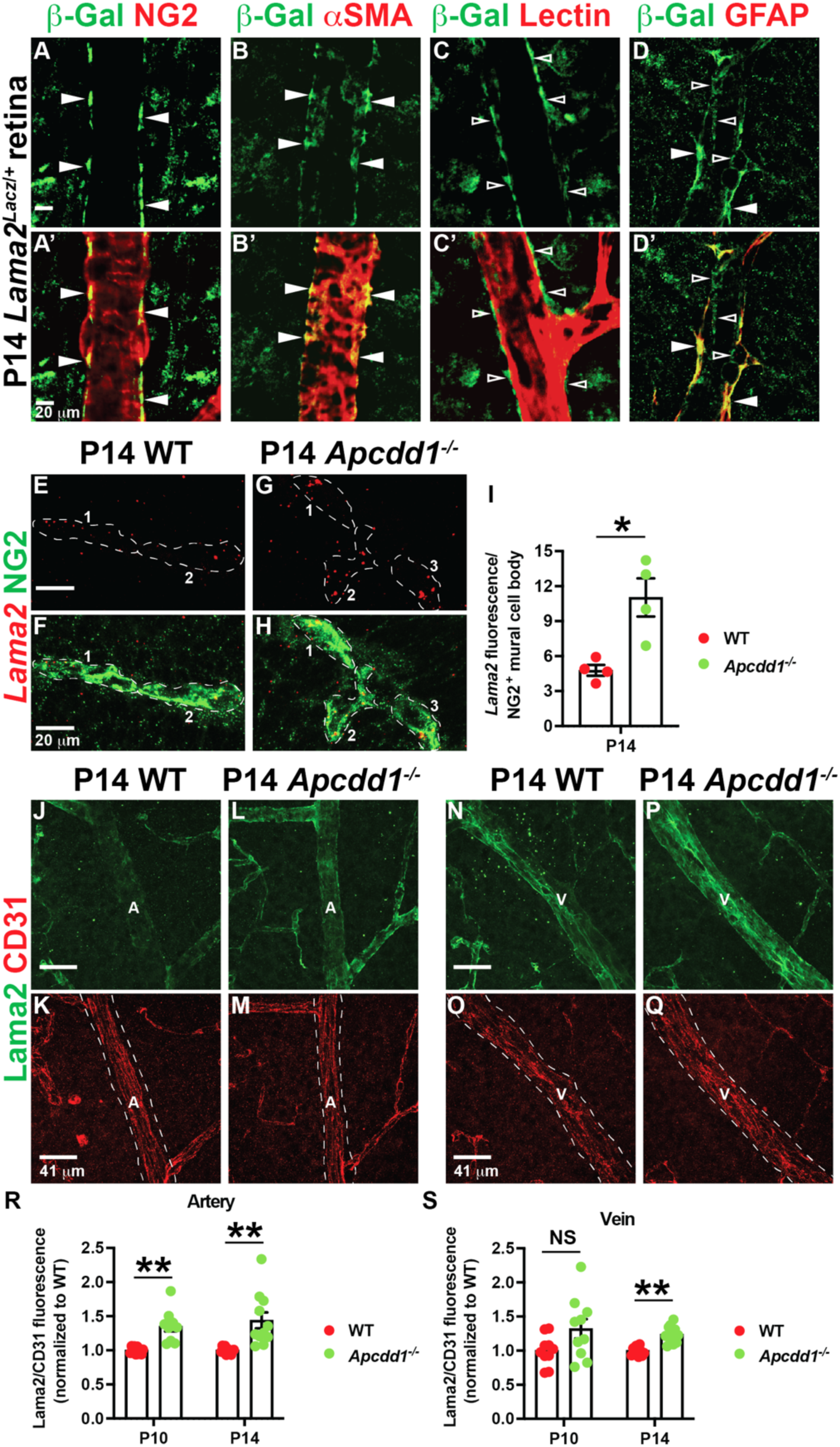
*Lama2* expression in mural cells and Lama2 deposition to the vBM is increased in *Apcdd1^-/-^* retinas. **A-D’)** P14 *Lama2^Lacz^*^/+^ retinal flat-mounts were labelled for β-galactosidase (β- gal; green) and either NG2 (mural cell marker; red), αSMA (vSMC marker; red), Lectin (EC marker; red) or GFAP (astrocyte marker; red). Empty arrowheads point to vascular β-gal not colocalized with ECs and astrocytes. Solid arrowheads point to vascular β-gal colocalized with mural cells, vSMCs and astrocytes. **E-H)** Fluorescent *in situ* hybridization of P14 WT and *Apcdd1*^-/-^ retinal sections with antisense probe against *Lama2* (red) along with immunolabeling for NG2 (green). Individual mural cell bodies are outlined with dashed lines. **I)** Dotted bar graph of *Lama2* mRNA mean fluorescence intensity (M.F.I) quantification of all RNA puncta in individual mural cell bodies (n=4 mice/genotype). **J-Q)** P14 WT and *Apcdd1*^-/-^ retinal flat-mounts were labelled for Lama2 (green) and CD31 (red). Arteries (A) and veins (V) are outlined with dashed white lines. **R, S)** Dotted bar graphs of the ratio of Lama2/CD31 mean fluorescence intensities in arteries and veins normalized to the WT average values at P10 and P14 (10 arteries or veins analyzed from n=5 mice/genotype). Student’s t-test *p<0.05, **p<0.02, NS: not significant. Dotted bar graphs show mean +/- SEM. Scale bars: A-D, E-H= 20 μm; J-Q = 41 μm.

To validate the bulk RNAseq data with regard to increased *Lama2* mRNA expression in *Apcdd1*^-/-^ mural cells, we performed fluorescent *in situ* hybridization with an antisense probe for *Lama2* together with immunolabeling for NG2 (mural cells) in P14 WT and *Apcdd1^-/-^* retinas. The number of *Lama2* mRNA puncta was higher in *Apcdd1*^-/-^ compared to WT retinal mural cells (**Fig. 3E-I**), supporting the bulk RNA seq data (**Fig. 2K, M; Sup. data sheet 4**). Next, we examined Laminin-α2 (Lama2) protein deposition into the vBM of WT and *Apcdd1*^-/-^ retinas. We used CD31 fluorescence intensity to normalize Lama2 fluorescence signal in the retinal vBM since CD31 mean fluorescent intensity (M.F.I) did not change between the two genotypes (**Fig. 3K, M, O, Q; data not shown**). Quantification of Lama2 relative to CD31 M.F.I in retinal vessels revealed that Lama2 deposition was upregulated by ∼25% in P10 arterial, but not venous, vBM (**Fig. 3R, S**), and by ∼30% in P14 arterial and venous vBMs (**Fig. 3J, L, N, P, R, S**) of *Apcdd1*^-/-^ compared to WT retinas. The effects were more pronounced in arteries since Lama2 protein deposition is lower in arterial compared to venous vBM at P14 retina (**Sup. Fig. 2d-f**). Western blot analyses for Lama1 and Lama2 proteins also showed higher Lama2 levels in P14 *Apcdd1*^-/-^ compared to WT retina, whereas Lama1 levels were similar (**Sup. Fig. 2g-i**). Quantification of Lama2 intensity relative to CD31 in cerebellar blood vessels also showed a 2-fold increase in vBM deposition in P14 *Apcdd1*^-/-^ compared to WT mice (**Sup. Fig. 3G-K**). In summary, while both ECs and mural cells have active Norrin/β-catenin signaling, only mural cells express *Lama2* mRNA. Retinal astrocytes also express *Lama2* mRNA; however, they lack active Norrin/β-catenin signaling. Loss of Apcdd1 leads to increased *Lama2* mRNA expression by mural cells and Lama 2 protein deposition in the vBM of both the retina and cerebellum. Thus, Apcdd1 controls Laminin-α2 expression in mural cells and deposition in the retinal vBM.

To confirm the relationship between Lama2 deposition in the vBM and neurovascular barrier maturation, we analyzed paracellular BRB permeability in *Lama2*^-/-^ mice (Kuang et al., 1998). We quantified accumulation of biocytin-tetramethylrhodamine (biocytin-TMR), an 870 Da molecular weight tracer that crosses the leaky or immature BBB/BRB (Lengfeld et al., 2017; Mazzoni et al., 2017) (**Fig. 4A**), into the *Lama2*^-/-^ and WT CNS parenchyma, 30 minutes after intravenous injections. There was a 5.5-fold increase in biocytin-TMR M.F.I. in both central and peripheral *Lama2*^-/-^ compared to WT retinas (**Fig. 4B-G**) indicative of a leaky BRB. In addition, expression of the tight junction protein Occludin was nearly absent in *Lama2*^-/-^ retinas (**Fig. 4H-K).** This is consistent with a previous study showing increased BBB leakage and loss of tight junction-associated proteins in *Lama2*^-/-^ brain (Menezes et al., 2014). The *Lama2* ^-/-^ vascular phenotype is opposite to that of the *Apcdd1^-/-^*retina where Occludin levels are increased in ECs (Mazzoni et al., 2017) and Lama2 deposition is upregulated in the vBM (**Fig.3, Sup. Fig. 3**).

**Figure 4.**
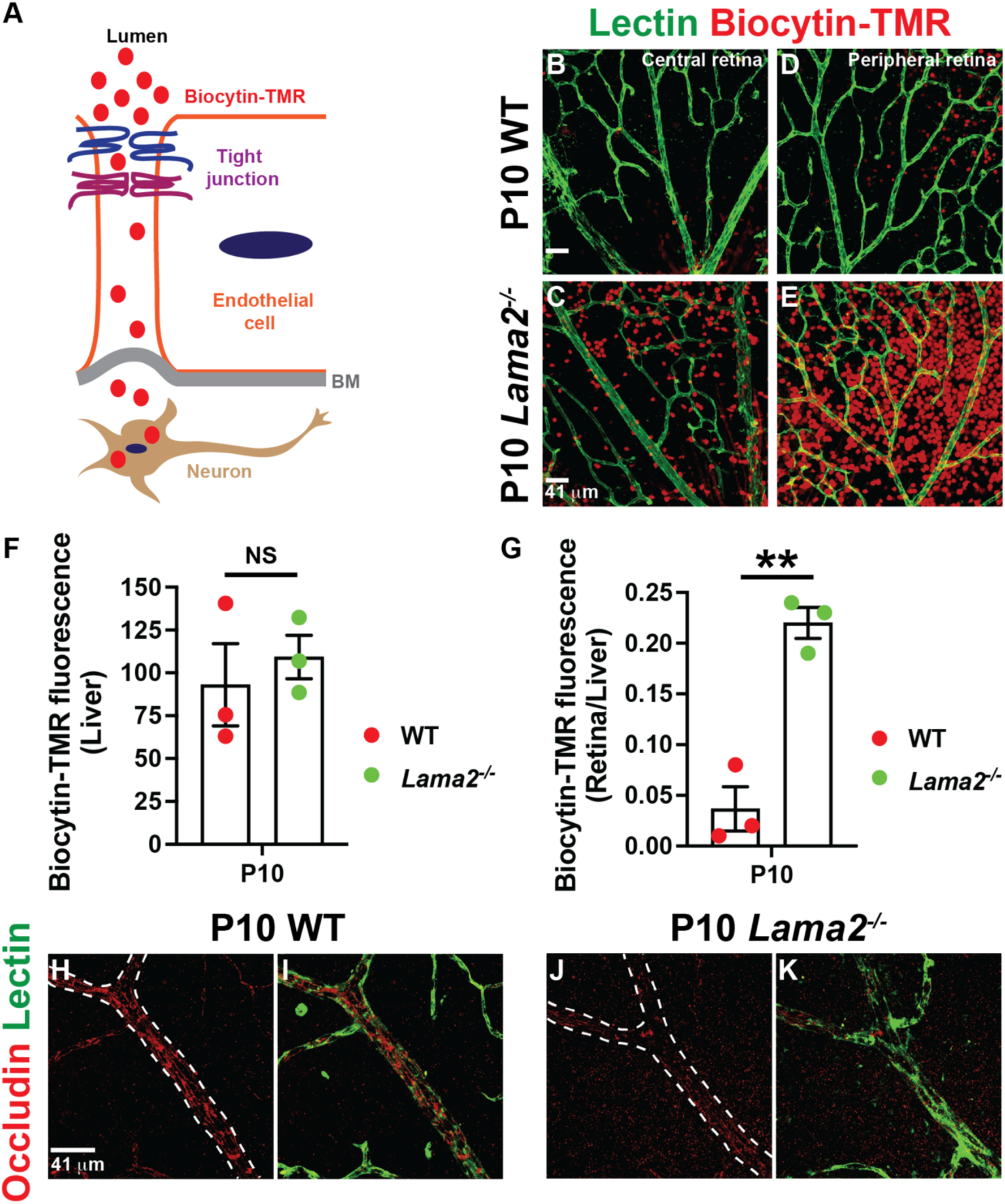
Paracellular BRB permeability is increased in *Lama2^-/-^*retinas. **A)** Schematic diagram of biocytin-tetramethylrhodamine (biocytin-TMR) extravasation from the blood into the CNS through the paracellular route. **B-E)** WT (**B, D**) and *Lama2*^-/-^ (**C, E**) retinal flat-mounts from P10 pups, injected with biocytin-TMR 30 minutes prior to sacrifice, were labelled for Lectin (green). *Lama2*^-/-^ retinas have a very high degree of biocytin-TMR extravasation into the parenchyma compared to WT controls. **F, G)** Dotted bar graphs of biocytin-TMR fluorescence intensities in the liver parenchyma (**F)** and the ratio of biocytin-TMR fluorescence intensities in the retina over liver parenchyma (**G**) (n=3 mice/groups; Student’s t-test; **p<0.02, NS: not significant). **H-K)** P10 WT (**H, I**) and *Lama2*^-/-^ (**J, K**) retinal flat-mounts were labelled for Occludin (tight junction protein, red) and Lectin (green). Occludin is almost absent in *Lama2*^-/-^ blood vessels. Student’s t-test **p<0.02, NS: not significant. Dotted bar graphs show mean +/- SEM. Scale bars: 41 μm.

### Norrin/β-catenin signaling positively regulates Lama2 expression and deposition in the neurovascular BM

The above data suggest that the precocious BBB/BRB maturation in *Apcdd1*^-/-^ cerebella and retinas (Mazzoni et al., 2017), is caused, in part, by higher levels of Lama2 in the vBM from increased Norrin/β-catenin activity in mural cells. To test this hypothesis, we cultured primary purified brain PCs and ECs plated into poly-D-lysine-coated dishes, in the presence of either DMSO (control) or the Wnt agonist CHIR99021 (1.0 μM)(Tran and Zheng, 2017) for 48 hours and analyzed expression of Lef1 and Lama2 proteins. CHIR99021 induced expression and nuclear localization of Lef1 by ∼3-fold (**Fig. 5A-C**) and increased Lama2 expression by ∼2-fold (**Fig. 5D-F**) in primary brain PCs (Pdgfrβ^+^ or NG2^+^ cells) *in vitro* within 48 hours. In contrast, there was no Lama2 expression (**Fig. 5G-I**) in either DMSO- or CHIR-treated primary brain ECs (Lectin^+^ cells) after 48 hours. These data suggest that canonical Wnt signaling functions as a positive regulator of Lama2 expression and secretion by CNS mural cells.

**Figure 5.**
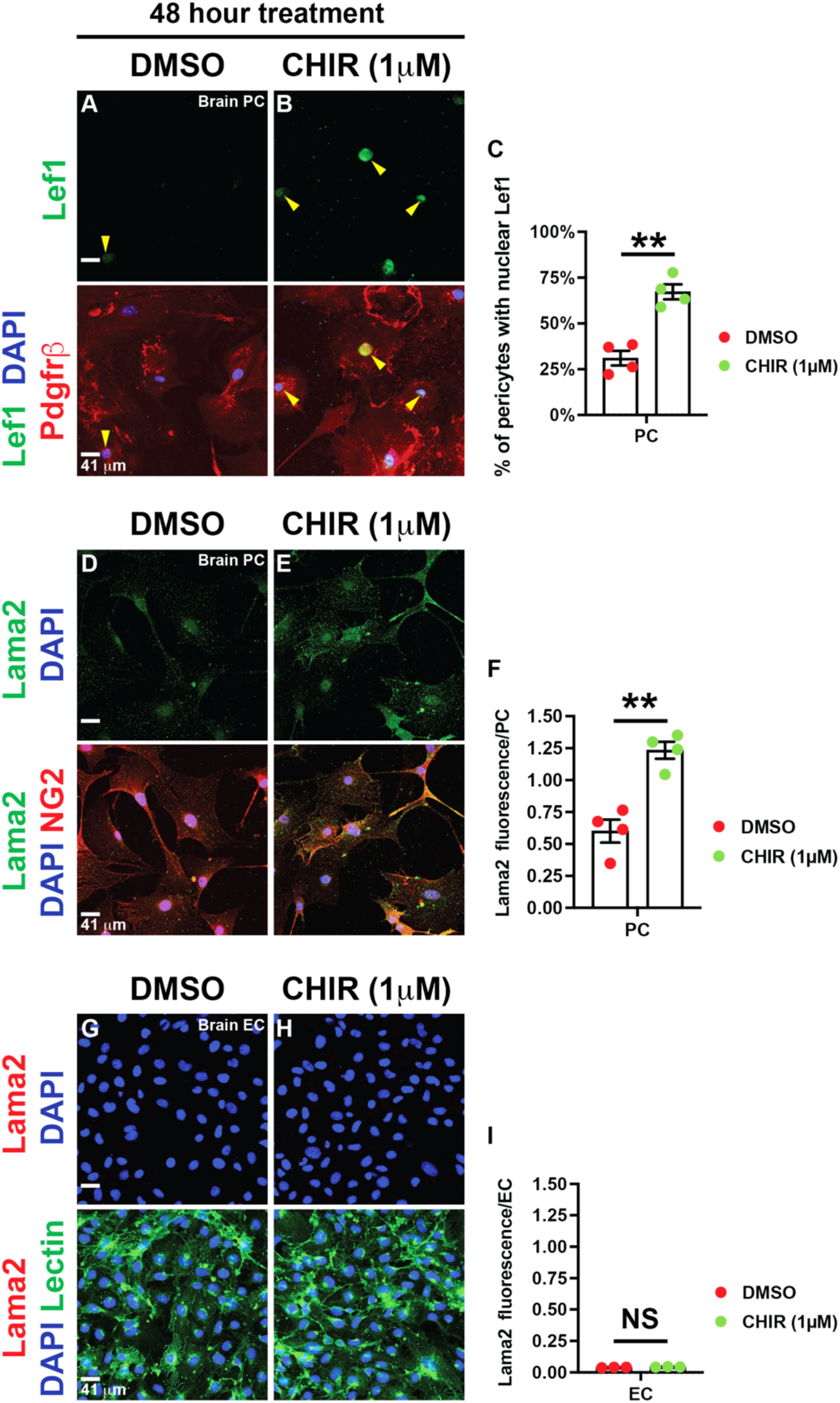
Lama2 expression is upregulated in primary brain PCs, but not in ECs, upon Norrin/β-catenin activation. **A, B)** Primary mouse brain pericytes (PCs) were treated with either DMSO (A; control) or the Wnt activator CHIR99021 (B; 1μM) for 48 hours, followed by staining for Pdgfrβ (red), Lef1 (green) and DAPI (blue). Yellow arrows point to Lef1^+^ PC nuclei. **C)** Quantification of the percentage of PCs with nuclear Lef1 expression (n=4 independent experiments). **D, E)** Primary mouse brain PCs were treated with either DMSO (D) or CHIR (E; 1μM) for 48 hours and stained for NG2 (red), Lama2 (green) and DAPI (red). **F)** Quantification of the Lama2 fluorescence intensities in PCs normalized to the cell number (n=4 independent experiments). **G, H)** Primary mouse brain ECs were treated with either DMSO (G) or CHIR (H; 1μM) for 48 hours and stained for Lectin (green), Lama2 (red) and DAPI (red). **I)** Quantification of the Lama2 fluorescence intensities in ECs normalized to the cell number (n=4 independent experiments). Student’s t-test; **p<0.02, NS: not significant. Dotted bar graphs show +/- SEM. Scale bars = 41 μm.

We next sought to identify which α2 chain-containing Laminin trimer is affected in the *Apcdd1*^-/-^ retina. Three laminin trimers contain the α2 chain: Laminin-211 (α2, β1, γ1), -221 (α2, β2, γ1) and -213 (α2, β1, γ3) (Patarroyo et al., 2002) (**Sup. Fig. 4A, J**). Previous studies have reported that loss of Laminin-γ1 chain in pericytes leads to BBB leakage (Gautam et al., 2016), whereas loss of Laminin-γ3 chain does not affect BRB integrity (Gnanaguru et al., 2013). Moreover, Laminin-γ3 chain is almost completely absent in the arterial vBM of P12 retinas (Biswas et al., 2018); thus Laminin-213 is unlikely to contribute to the increase in α2 chain- containing Laminin trimer deposition in the *Apcdd1*^-/-^ retinal vBM. To distinguish between the remaining laminin trimers (Laminin-211 and 221), we examined vBM deposition of Laminin-β1 and -β2 chains in both WT and *Apcdd1*^-/-^ retinas. The deposition of Laminin-β1 chain followed a similar pattern to Laminin α2, i.e., it was upregulated primarily in the arterial (∼30%) and to a smaller extent in the venous (∼10%) vBM in *Apcdd1*^-/-^ compared to WT retinas (**Sup. Fig. 4B-I, S, T**). In contrast, the vBM deposition of Laminin-β2 chain in the retina was not different between the two genotypes (**Sup. Fig. 4K-R, U, V**). The P14 retinal single-cell RNA-sequencing database (Macosko et al., 2015) confirmed that mural cells are the main source of *Lamb1* (Laminin-β1 chain) and *Lamc1* (Laminin-γ1 chain), whereas *Lamb2* (Laminin-β2 chain) is predominantly expressed by astrocytes (**Sup. Fig. 4W)**. Thus, modulation of Norrin/β-catenin signaling in mural cells by Apcdd1 regulates Lama2 deposition in the retinal vBM, with Laminin-211 likely being the most affected laminin trimer.

### Astrocytes mature precociously in the *Apcdd1*^-/-^ retina

Next, we asked whether the NVU assembly is affected in response to increased Lama2 deposition in *Apcdd1*^-/-^ retinal vBM. We have previously shown that loss of Apcdd1 does not affect mural cell coverage of retinal vessels (Mazzoni et al., 2017). Since astrocytes adhere to the vBM through their endfeet and changes in the vBM composition affect astrocyte homeostasis (Armulik et al., 2010; Menezes et al., 2014), we asked whether astrocyte polarization and attachment are affected in response to increased Lama2 deposition in *Apcdd1*^-/-^ retinal vBM. Our bulk RNAseq data showed that several transcripts expressed by mature astrocytes (e.g., *Mlc1*, *Apoe*, *Gja1* and *Gfap*) (Li et al., 2019; Tao and Zhang, 2014) are upregulated in *Apcdd1*^-/-^ retinas only at P14, with the exception of two transcripts, *Plcd4* and *Aqp4* (Li et al., 2019), that are also upregulated at P10 *Apcdd1*^-/-^ retina (**Fig. 6A, B; Sup. data sheet 5**). In contrast, the transcript for an immature astrocyte marker, *Sh3pxd2b* (Li et al., 2019), is downregulated at P14 *Apcdd1*^-/-^ retina (**Fig. 6B**). Thus, transcriptional changes related to astrocyte maturation become significant at a later developmental stage compared to those in either ECs or mural cells in the *Apcdd1*^-/-^ retinas.

**Figure 6.**
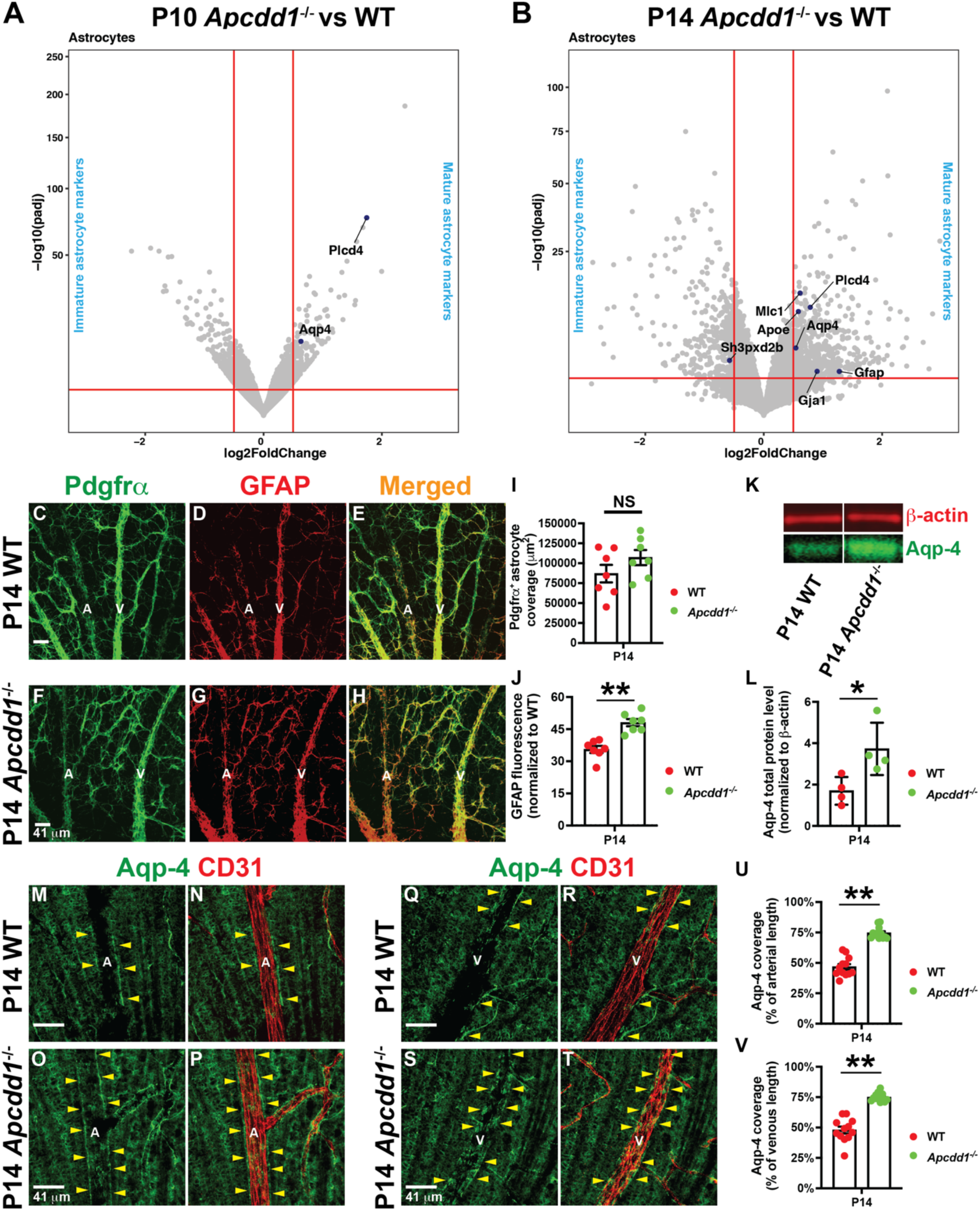
*Apcdd1^-/-^* retinas show precocious astrocyte maturation and endfeet polarization. **A, B)** Volcano plots of statistically significant differentially expressed astrocyte maturation genes (blue dots) between *Apcdd1*^-/-^ and WT retinas at P10 (**A**) and P14 (**B**). Conventions as outlined in Figure 2H-M. **C-H)** P14 WT and *Apcdd1*^-/-^ retinal flat-mounts were labelled for Pdgfrα (green) to label all astrocytes and GFAP (red) to label mature astrocytes. A: arteries and V: veins. **I, J)** Dotted bar graphs of Pdgfrα^+^ astrocyte coverage (**I**; n=7 mice/group) and GFAP mean fluorescence intensity (**J**; n=7 mice/genotype). **K)** P14 WT and *Apcdd1*^-/-^ whole retinal lysates were probed for Aquaporin-4 (Aqp-4: green bands) and β-actin (red bands) by western blot. **L)** Quantification of total Aqp-4 protein level (normalized to β-actin) by western blot (n=4 samples/ group). **M-T)** P14 WT and *Apcdd1*^-/-^ retinal flat-mounts were labelled for Aqp-4 (green; polarized astrocyte endfeet) and CD31 (red). Yellow arrowheads point at Aqp-4^+^ astrocyte endfeet around blood vessels. A: arteries and V: veins. **U-V)** Quantification of the percentage of vascular length surrounded by Aqp-4^+^ astrocyte endfeet at P14 (12 arteries or veins analyzed from n=6 mice/group). Student’s t-test *p<0.05, **p<0.02, NS: not significant. Dotted bar graphs show mean +/- SEM (I, J, U, V) or SD (L). Scale bars: 41 μm.

To validate these bulk RNA seq findings, we labeled P14 WT and *Apcdd1*^-/-^ retinas for Pdgfrα and GFAP, marking all or mature astrocytes, respectively (Tao and Zhang, 2014). While Pdgfrα^+^ astrocyte coverage was unchanged, GFAP M.F.I. immunoreactivity was increased by ∼30% in *Apcdd1*^-/-^ compared to WT retinas (see Materials and Methods for quantification details**; Fig. 6C-J**). These data suggest that retinal astrocytes mature precociously in *Apcdd1*^-/-^ retinas between P10 and P14, in parallel with a precocious BRB maturation (Mazzoni et al., 2017). Müller glia, which span the thickness of the retina, also upregulate GFAP expression during gliosis (Iandiev et al., 2006) prompting us to examine whether Müller gliosis contributes to increased GFAP expression in *Apcdd1*^-/-^ retinas. GFAP was detected only in astrocytes over the ganglion cell layer (GCL) in both P14 WT and *Apcdd1*^-/-^ retinal sections, with no expression in deeper layers where Müller cell bodies and processes are located (data not shown). Thus, GFAP upregulation in the *Apcdd1*^-/-^ retina occurs exclusively in astrocytes.

Next, we examined astrocyte endfeet polarization around retinal blood vessels. Aquaporin-4 (Aqp-4), a water channel protein expressed by mature astrocytes (Li et al., 2019), exhibits polarized expression in astrocyte endfeet surrounding blood vessels. We measured total Aqp-4 protein levels by western blot and found that it was significantly upregulated in *Apcdd1*^-/-^ compared to WT retinas (**Fig. 6K, L**). Next, we labeled WT and *Apcdd1*^-/-^ retinas with Aqp-4 and CD31, and quantified the percentage of vessel length covered by Aqp-4^+^ astrocyte endfeet. Aqp-4^+^ astrocyte endfeet coverage of retinal vessels was increased by ∼45% in P14 *Apcdd1*^-/-^ compared to WT mice (**Fig. 6M-V**). Interestingly, there was no difference in Aqp-4 coverage of deeper vascular plexus by Müller glia endfeet between *Apcdd1*^-/-^ and WT retinas (data not shown), suggesting that these phenotypes are confined to the astrocytes in the superficial vascular plexus.

These data suggest a causal relationship between increased Lama2 expression, and precocious astrocyte maturation and endfeet polarization in *Apcdd1*^-/-^ retinas. To confirm this relationship, we examined astrocyte maturation and polarization in *Lama2*^-/-^ retinas. Polarized expression of Aqp-4 in astrocyte end-feet around superficial plexus blood vessels was drastically reduced in the *Lama2*^-/-^ retina (**Sup. Fig. 5A-D**), consistent with previous observations in the brain (Menezes et al., 2014). Given the critical role of polarized Aqp-4 expression in maintaining BRB integrity (Nicchia et al., 2004), these data explain, in part, the precocious BRB maturation in the *Apcdd1*^-/-^ retina, and the opposite phenotype in the *Lama2*^-/-^ retina. Finally, we asked whether astrocyte endfeet polarization is affected in the *Apcdd1*^-/-^ cerebellum. Similar to the retina, Aqp-4 immunoreactivity was also increased in astrocyte endfeet around *Apcdd1*^-/-^ cerebellar vessels (**Sup. Fig. 6A-D)**. Taken together, these data indicate that the amount of Lama2 deposition in the vBM affects the timing of astrocyte maturation and polarization in the developing CNS.

### Astrocytic expression of Integrin-α6 is increased in *Apcdd1*^-/-^ retina to compensate for Lama2 levels

Next, we asked how retinal astrocytes are affected in the *Apcdd1*^-/-^ retina. We examined expression of Lef1 and found that it was absent in P10 *Apcdd1*^-/-^ retinal astrocytes (**Fig. 7A-D: yellow arrowheads**), indicative of no aberrant Norrin/β-catenin signaling activation by *Apcdd1*^-/-^ retinal astrocytes. Although astrocytes do not show aberrant Norrin/β-catenin signaling activation at P10 *Apcdd1*^-/-^ retina, we hypothesized that their maturation is accelerated by P14, due to increased adhesion to the vBMs from higher Lama2 levels. Previous studies have shown that several ECM receptors including Integrin-α2, -α6, -β1 and -β4 chains expressed by astrocytes (Gnanaguru et al., 2013; Nirwane and Yao, 2018), are affected by changes in vBM protein composition. For example, Integrin-β1 expression at the gliovascular interphase is significantly decreased in the *Lama2*^-/-^ brain, and blocking Integrin-β1-mediated adhesion in astrocytes significantly decreases Aqp-4 clustering at the cell-ECM interphase (Menezes et al., 2014); however the α-partner of Integrin-β1 responsible for this process was not identified in this study. Since *Apcdd1*^-/-^ retina exhibits an opposite phenotype to that of *Lama2*^-/-^ retina, we hypothesized that astrocytic Integrin expression must increase in response to higher levels of Lama2 in the vBM. Consistent with this hypothesis, our bulk RNAseq revealed that *Itga6* (Integrin-α6) levels were significantly increased in the *Apcdd1*^-/-^ retina at both P10 and P14 (**Fig. 2L, M; Sup. data sheet 4**). We also found that while Integrin-α6 protein is localized at higher levels at retinal astrocyte processes as well as endfeet characterized by higher GFAP expression (**Fig. 7H-J: solid arrowheads**), Integrin-α2 (also upregulated in the *Apcdd1*^-/-^ retina at P10; (**Fig. 2L, sup. data sheet 4**) did not colocalize to astrocyte endfeet (**Fig. 7E-G: empty arrowheads**). These data are consistent with *in vitro* finding that Integrin-α6β1 binds several Laminin trimers, including Laminin-211 (Nishiuchi et al., 2006). Quantification of the ratio of Integrin-α6 over GFAP fluorescence at P10 and P14 WT and *Apcdd1^-/-^* retinas showed a significant upregulation of Integrin-α6 intensity in *Apcdd1*^-/-^ retinal astrocytes at both P10 (∼40%; **Fig. 7O, T**) and P14 (∼50%; **Fig. 7K-T**) compared to WT astrocytes. These observations are consistent with a previous report that Integrin expression is regulated by the presence of the cognate Laminin ligands in the vBM (Gnanaguru et al., 2013). Integrin-α6 expression was also increased in *Apcdd1*^-/-^ compared to WT cerebellar astrocytes (**Sup. Fig. 6E-H**), similar to the retina. Thus, astrocytic Integrin-α6β1 is likely responsible for mediating interactions of astrocytes with Lama2 in the vBM and this interaction affects astrocyte maturation and polarization around blood vessels in the developing CNS.

**Figure 7.**
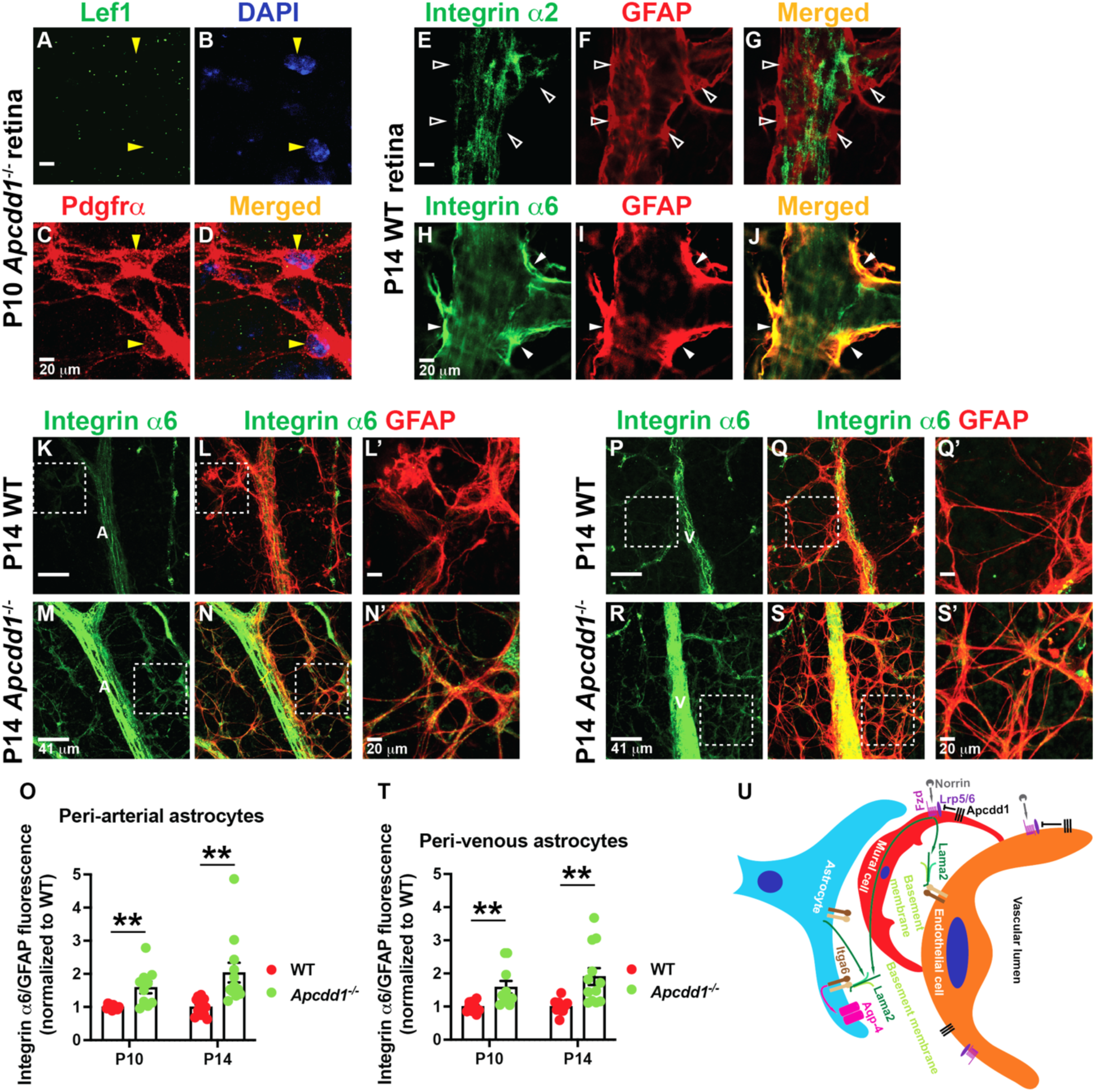
Astrocytic Integrin. α**6 expression is upregulated in the *Apcdd1^-/-^* retinas. A-D)** P10 *Apcdd1*^-/-^ retinal flat-mounts were labelled for Lef1 (green), Pdgfrα (astrocyte marker; red) and DAPI (blue). Yellow arrowheads point to Lef1^-^ astrocytes. **E-G)** P14 WT retinal flat-mounts were labelled for Integrin α2 (green) and Gfap (red). Empty arrowheads point at the lack of Integrin α2 expression in Gfap^+^ astrocyte end-feet. **H-J)** P14 WT retinal flat-mounts were labelled for Integrin α6 (green) and Gfap (red). Solid arrowheads point at the Integrin α6 expression in Gfap^+^ astrocyte end-feet. **K-N’)** P14 WT and *Apcdd1*^-/-^ retinal flat-mounts were labelled for Integrin α6 (green) and GFAP (red). Boxed areas in l and n are magnified in l’ and n’, respectively. A = artery. **O)** Dotted bar graph of the ratio of Integrin α6/GFAP fluorescence intensities in peri-arterial astrocytes, and normalized to the WT average values at P10 and P14 (10 arteries analyzed from n=5 mice/group at P10 and 12 arteries analyzed from n=6 mice/group at P14). **P-S’)** P14 WT and *Apcdd1*^-/-^ retinal flat-mounts were labelled for Integrin α6 (green) and GFAP (red). Boxed areas in q and s are magnified in q’ and s’, respectively. V = vein. **T)** Dotted bar graph of the ratio of Integrin α6/GFAP fluorescence intensities in peri-venous astrocytes, and normalized to the WT average values at P10 and P14 (10 veins analyzed from n=5 mice/group at P10 and 12 arteries analyzed from n=6 mice/group at P14). U) The schematic diagram illustrates the proposed model by which Norrin/β-catenin signaling in mural cells regulates Lama2 deposition in the vBM and NVU maturation (see Discussion for explanation). Student’s t -test **p<0.02. Dotted bar graphs show mean +/- SEM. Scale bars: **A-Jj, L’, N’, Q’, S’**: 20 μm; **K, L, M, N, P, Q, R, S**: 41 μm.

## DISCUSSION

The establishment and maintenance of BBB/BRB integrity is critical to support neuronal function and restrict the entry of immune cells and pathogens into the healthy CNS. Vascular BM proteins, especially laminins and Collagen IV, play a critical role in maintaining BBB/BRB integrity (Armulik et al., 2010; Chen et al., 2013; Cottarelli et al., 2020; Gautam et al., 2016; Gnanaguru et al., 2013; Menezes et al., 2014; Sixt et al., 2001; Yao et al., 2014). However, the molecular mechanisms that regulate expression and/or deposition of various vascular BM components in CNS blood vessels remain elusive. Elucidation of these mechanisms may also provide potential therapeutic strategies to restore neurovascular barrier integrity in either neurological or ocular disorders associated with impaired BBB or BRB, respectively. Our study provides an important step forward to understand which signaling pathway induce expression of one vBM protein, Laminin, in the developing CNS vasculature and elucidate a missing link between the well-established role of Norrin (Wnt)/β-catenin signaling in BBB/BRB maturation and the role of vBM in this process. Prior studies that address the role of Norrin/β-catenin pathway in the CNS vasculature and BBB/BRB formation have been focused on ECs, since they are the main NVU cell type that show high Norrin/β-catenin pathway activation (Cottarelli et al., 2020; Daneman et al., 2009; Lengfeld et al., 2017; Liebner et al., 2008; Mazzoni et al., 2017; Wang et al., 2018; Wang et al., 2012; Ye et al., 2009; Zhou et al., 2014). To our knowledge, this is the first study to demonstrate that: a) Norrin/β-catenin signaling is activated in CNS mural cells, and b) Norrin/β-catenin signaling regulates Lama2 expression and deposition into the vBM and as a consequence the maturation of astrocytes and neurovascular barrier properties. Below we discuss: a) how Norrin/β-catenin signaling in mural cells regulates the composition of the vBM proteins and the BBB/BRB maturation and b) the effects of vBM protein composition for NVU assembly, astrocyte and neurovascular barrier maturation.

Although it is well-established for more than a decade that Norrin/β-catenin signaling is activated in ECs during brain and retina development [reviewed in (Biswas et al., 2020)], we have found that retinal and cerebellar mural cells also have active Norrin/β-catenin signaling, albeit at a lower level than ECs, by examining expression of downstream Norrin/β-catenin signaling targets Lef1(Fancy et al., 2014) and *Apcdd1* (Mazzoni et al., 2017; Shimomura et al., 2010), as well as nGFP in *Tcf/Lef*::H2B-nGFP Wnt reporter mice (Ferrer-Vaquer et al., 2010; Lengfeld et al., 2017) (**Fig. 1, 2** and **Sup. Fig 1**). Moreover, we have found that Lef1 is translocated into the nucleus in primary pericytes upon pharmacological activation of Wnt/β-catenin signaling (**Fig. 5**). Our findings are consistent with published single cell RNA sequencing studies that transcripts for several Norrin/β-catenin signaling components including receptors (*e.g., Fzd4*, *Lrp5*), downstream effectors (*e.g., Lef1)* and negative regulators (*e.g., Apcdd1, Axin2*) are expressed by P14 retinal and adult brain mural cells (He et al., 2018; Macosko et al., 2015; Vanlandewijck et al., 2018). In contrast, retinal or cerebellar astrocytes, that are in close contact with either a Wnt or Norrin source (Mazzoni et al., 2017; Wang et al., 2018; Wang et al., 2012; Ye et al., 2009; Zhou et al., 2014), do not show active Norrin/β-catenin signaling during BRB/BBB maturation, respectively. During CNS development, Wnt or Norrin is produced by either neural progenitors (brain), Müller glia (retina) or Bergman glia (cerebellum), and as a consequence these ligands are available to interact with their cognate receptors in either ECs or mural cells. Why is the activation of canonical Wnt signaling lower in CNS mural cells than ECs? One possibility is that negative regulators of Norrin/β-catenin signaling may be expressed at higher levels in mural cells. Consistent with this hypothesis, expression levels of *Axin2* mRNA are higher in adult brain or P14 retinal PCs than in ECs (He et al., 2018; Macosko et al., 2015; Vanlandewijck et al., 2018). *Nkd2*, another negative regulator of canonical Wnt signaling, is expressed exclusively by PCs in the adult brain, and shows higher expression in retinal PCs than ECs (He et al., 2018; Macosko et al., 2015; Vanlandewijck et al., 2018). Alternatively, several receptors that activate non-canonical Wnt signaling (e.g., Ryk) (Green et al., 2014) are almost exclusive expressed by brain mural cells (He et al., 2018; Vanlandewijck et al., 2018), and shows higher expression in P14 retinal PCs than ECs (Macosko et al., 2015). It is possible that expression of these negative regulators of Norrin/β-catenin signaling reduces pathway activation more rapidly in CNS mural cells compared to ECs. Future studies are needed to address the dynamic nature of Norrin/β-catenin activation in mural cells during CNS development.

Although vBM components such as Collagen IV and Laminins are important for maintaining BBB/BRB integrity (Baeten and Akassoglou, 2011; Chen et al., 2013; Cottarelli et al., 2020; Gautam et al., 2016; Gnanaguru et al., 2013; Gould et al., 2005; Menezes et al., 2014; Sixt et al., 2001; Yao et al., 2014), the signaling pathways that regulate their expression by NVU-forming cells and deposition to vBM is still unknown. We have recently shown that expression of Fgfbp1, a downstream effector of the Wnt/β-catenin signaling in ECs, regulates Collagen IV deposition in the vBM through the activity of Fgfbp1 (Cottarelli et al., 2020). However, it is unknown whether Wnt/β-catenin signaling may regulate deposition of other key vBM proteins such as Laminins. While traditionally astrocytes were thought to be the main NVU cell source for Lama2 expression and deposition into the vBM (Sixt et al., 2001), recent single-cell RNAseq studies have shown the mural cells are a major source of *Lama2* mRNA transcript (encoding for Laminin α2 chain) in either adult brain or P14 retina, whereas astrocytes contribute to a lesser extent (Armulik et al., 2010; He et al., 2018; Macosko et al., 2015; Menezes et al., 2014; Vanlandewijck et al., 2018). We confirmed these RNAseq findings by using *Lama2^Lacz^*^/+^ reporter mice where we found that vascular β-Gal was colocalized with mural cells in the retina and cerebellum of *Lama2^Lacz^*^/+^ mice. We also observed partial vascular β-Gal colocalization with retinal astrocytes, reflecting some Lama2 expression by astrocytes (**Fig. 3A-D’**). Although Lama2 has been shown to be critical for maintaining BBB integrity (Menezes et al., 2014), our findings demonstrate that it is also essential for maintaining BRB integrity (**Fig. 4**). Loss of Laminin α2 chain dramatically disrupts the expression and organization of EC tight junction proteins such as Claudin5 and Occludin in brain ECs (Menezes et al., 2014). We also found a drastic reduction of Occludin expression in the *Lama2^-/-^* retinal ECs consistent with impaired paracellular BRB permeability (**Fig. 4**). This is the opposite phenotype of the *Apcdd1^-/-^* retina (Mazzoni et al., 2017) where Norrin/β-catenin signaling as well as Lama2 expression and deposition into the vBM are upregulated (**Fig. 3**). Previous studies have shown that genetic ablation of the Norrin/β-catenin signaling in retinal ECs reduces expression of tight junction proteins (Wang et al., 2018; Wang et al., 2012). Thus, expression of tight junction proteins and organization of endothelial cell junctions that control paracellular barrier integrity in the CNS ECs seem to be regulated by at least two mechanisms: a) activation of Norrin/β-catenin signaling in ECs, and b) laminin composition of the CNS vBM.

Our findings indicate that Norrin/β-catenin activity in retinal and cerebellar mural cells is an upstream and positive regulator of Lama2 expression and deposition to the vBM. In *Apcdd1*^-/-^ mice, where Norrin/β-catenin signaling is upregulated leading to a precocious maturation of the BRB between P10-P14 (Mazzoni et al., 2017), we find that *Lama2* mRNA transcript levels are increased specifically in mural cells leading to higher deposition of Lama2 in the vBM by four independent approaches: 1) bulk RNAseq comparison between wild-type and *Apcdd1*^-/-^ retinas, 2) RNA *in situ* hybridization with an antisense probe against *Lama2* mRNA, 3) Western blotting of whole retinal lysates, and 4) immunolabelling of retinas and cerebella for Lama2 (**Fig. 2, 3 and Sup. Fig. 2**). Given that both ECs and mural cells express Apcdd1 and ECs do not produce Lama2, our findings suggest that Norrin/β-catenin signaling in mural cells facilitates Lama2 expression and deposition in the vascular BM in the developing CNS. Consistent with this hypothesis, pharmacological activation of Norrin/β-catenin signaling in primary brain pericytes increases Lama2 production *in vitro* (**Fig. 5**). In contrast, since retinal astrocytes do not have active Norrin/β- catenin signaling in either WT or *Apcdd1*^-/-^ mice; therefore, Lama2 production by astrocytes does not seem to be under the control of canonical Wnt signaling. Since mural cells are the predominant source of *Lama2* mRNA in the retina (**Sup. Fig. 2**), our findings indicate that the majority of Lama2 production and deposition in the retina vBM is thus regulated by Norrin/β-catenin activity in mural cells. It should be noted that only a subset (∼20%) of retinal pericytes express *Apcdd1* mRNA *in vivo* compared to ECs (**Fig. 2A-G**) which could be due to a more rapid loss of Norrin/β- catenin activation in mural cells compared to ECs during development (**Fig. 1K, S;** data not shown). This finding may account for the modest effects on *Lama2* mRNA expression and Lama2 deposition in the vBM observed in *Apcdd1^-/-^* compared to WT retinas.

It is possible that interactions between endothelial and mural cells are affected in the *Apcdd1*^-/-^ retina, which in turn can increase Lama2 production by mural cells as a secondary effect. However, our data from the bulk RNA seq comparison at P10 and P14 retinas and *in vitro* cultures with primary brain PCs argue that Norrin/β-catenin activation plays a direct effect on Lama2 production by brain PCs. Additionally, none of the transcripts for several signaling pathways responsible for endothelial-mural cell interactions (e.g., *Tgfb*, *Tgfbr1*, *Smad2/3*, *Smad2/3*, *Ang1*, *Ang2*, *Tie2*, *Jag1*, *Notch3*) were differentially expressed between *Apcdd1*^-/-^ and WT retinas in our bulk RNAseq analyses (**Sup. data sheet 1-3**). Thus, the most plausible explanation is that Norrin/β-catenin activity in mural cells directly facilitates Lama2 expression and deposition in the vascular BM.

What are the effects of increased Lama2 deposition on NVU cell interactions in the *Apcdd1*^-/-^ retina? Our bulk RNA seq studies, confirmed by immunofluorescence and Western blotting, demonstrate that *Apcdd1*^-/-^ retinal astrocytes have increased expression of several maturity markers including *Gfap* (Tao and Zhang, 2014) and *Aqp4* (Li et al., 2019) by P14. Aqp-4 expression is lost from polarized astrocyte endfeet in several retinovascular disorders associated with BRB dysfunction such as experimental autoimmune uveitis (Motulsky et al., 2010) and genetic ablation of *Aqp4* also leads to disrupted BRB integrity (Nicchia et al., 2004). Our findings are consistent with previous studies demonstrating that Aqp-4 localization in glial endfeet is regulated by the BM composition, and that loss of either Lama2 (Menezes et al., 2014) or Lamb2 and Lamc3 together (Hirrlinger et al., 2011) drastically reduce Aqp-4 localization in glial endfeet in the brain and retina, respectively. In addition, our bulk retinal RNAseq and immunohistochemical analyses at P14 demonstrate that Integrin α6 (*Itga6*) expression was upregulated in the *Apcdd1*^-/-^ retinal astrocytes. Integrin-α6 forms a dimer with β1 (Baeten and Akassoglou, 2011), an ECM receptor expressed in astrocytes, to bind various Laminins including Laminin-211 (Menezes et al., 2014; Nishiuchi et al., 2006). We postulate that Integrin α6β1 binding to α2-containing Laminins mediates the interaction of retinal astrocytes with the vascular BM. However, we cannot rule out the involvement of other Integrins, such as α6β4, in this process.

Our findings lead us to propose a new model of the role of Norrin/β-catenin signaling in retinal and cerebellar NVU assembly and BRB/BBB maturation (**Fig. 7U**). The activation of Norrin/β-catenin signaling in retinal mural cells, rather than ECs, leads to production and deposition of Lama2 (likely Laminin-211) in the vBM. Astrocytes also produce Lama2 chain, but they do not have active Norrin/β-catenin signaling. Thus, the upregulation of Norrin/β-catenin signaling due to the loss of Apcdd1 increases Lama2 production by mural cells, but has no effect on the astrocytic production of Lama2. Astrocytes express Integrin α6β1, a Laminin receptor whose expression is upregulated in response to increased deposition of Lama2 in the in the *Apcdd1*^-/-^ vBM. We postulate that the increased binding of Integrin-α6β1 to Lama2 in the vBM of *Apcdd1*^-/-^ retinas and cerebella induces a precocious maturation of astrocytes leading to precocious polarization of astrocyte endfeet around blood vessels reflected in upregulation of Aqp-4 transcript and protein levels around CNS blood vessels in *Apcdd1*^-/-^ retinas. Consequently, these changes lead to a precocious BBB/BRB maturation in the *Apcdd1*^-/-^ retinas and cerebella. Thus, Norrin/β- catenin signaling in retinal and cerebellar mural cells functions upstream of Laminin production and is a positive modulator of both astrocyte and neurovascular barrier maturation.

In conclusion, our study shows a novel mechanism by which Norrin/β-catenin signaling in retinal and cerebellar mural cells regulates Lama2 production and deposition in the vascular BM, and thereby modulates NVU and BBB/BRB integrity. Since Norrin/β-catenin signaling in both ECs and mural cells regulates BBB/BRB integrity, this may be a potential target for pharmacological intervention in neurovascular diseases where BBB/BRB integrity is compromised.

## MATERIALS AND METHODS

### Mice

All animal usage and protocols were approved by the Institutional Animal Care and Use Committee (IACUC) at Columbia University Irving Medical Center. Experimental procedures were optimized to minimize animal stress and the number of animals used for experiments. Both sexes (males and females) were used for experiments and analyses.

#### *Apcdd1* mutant mice genotyping

Generation and validation of *Apcdd1*^-/-^ mice (CD1 background) was described before (Mazzoni et al., 2017). These mice were bred as *Apcdd1*^+/-^ and gave birth in Mendelian ratios. Age-matched littermate WT and *Apcdd1*^-/-^ mice were used for experiments. PCR primers for the *Apcdd1* wild-type allele are forward (5′- GAGTGTCCCCGACTCCGACTCT -3′) and reverse (5′- ATGTGTTGAGTTATTCCCGGAAG -3′). PCR primers for the *Apcdd1* mutant allele are forward (5′- GCCCATCTGGGATAGTACATGTG -3′) and reverse (5′- CCTGGACTGAGGGCCATAGGTAAGAGG -3′).

#### *Tcf/Lef*::H2B-eGFP mice genotyping

Generation and validation of *Tcf/Lef*::H2B-eGFP transgenic mice (BL/6J background) was described before (Lengfeld et al., 2017). PCR primers for the *GFP* allele are forward (5′- CCCTGAAGTTCATCTGCACCAC -3′) and reverse (5′- TTCTCGTTGGGGTCTTTGCTC -3′).

#### *Lama2^Lacz^*^/+^ mice genotyping

*Lama2^Lacz^*^/+^ mice (B6.129S1(Cg)-*Lama2^tm1Eeng^*/J, RRID:IMSR_JAX:013786 (Kuang et al., 1998)) were obtained from the Jackson Laboratory (Bar Harbor, ME). Breeding between *Lama2^Lacz^*^/+^ mice gave birth in Mendelian ratios, and age-matched littermate WT, *Lama2^Lacz^*^/+^ and *Lama2^Lacz^*^/*Lacz*^ (*Lama2*^-/-^) mice were used for experiments. PCR primers for the *Lama2* wild-type and mutant alleles are forward (5′- ACTGCCCTTTCTCACCCACCCTT -3′) and reverse (WT: 5′- GTTGATGCGCTTGGGAC -3′; *Lacz*: 5′- CGACAGTATCGGCCTCAG -3′). *Lama2*^-/-^ pups died around P14 due to complications associated with merosin (Laminin α2)-dependent congenital muscular dystrophy.

### Primary vascular brain pericyte (PC) and endothelial cell (EC) cultures

Primary brain vascular PCs (iXCells; San Diego, CA; Cat# 10MU-014) and ECs (Cell Biologics; Chicago, IL; Cat# C57-6023) were cultured in dishes were coated with Poly-D-Lysine (PDL; 1 mg/ml) and Collagen IV (20 μg/ml) and pericytes were seeded onto the coated plates. Cells were cultured with pericyte (IxCells; San Diego, CA, Cat# MD-0030) or endothelial (Cell Biologics; Chicago, IL; Cat# M1168) growth media. Media were changed every 48 hours until the cells were 80% confluent. Next, the cells were detached with 0.25% Trypsin and plated on 24-well glass plates coated with PDL and Collagen IV. Media were changed every 24 hours. CHIR-99021 = 1 μM (Selleck Chemicals; Houston, TX; Cat# S1263) or DMSO (control) were added to the media after 24 hours and repeated every 24 hours for 2 days. Then the media were aspirated, and the cells were fixed with 2% PFA for 10 minutes and processed for immunohistochemistry. Briefly, cells were blocked for 2 h at room temperature followed by primary antibody incubation overnight at 4°C. Following 1X PBS washes, cells were incubated with secondary antibodies for 4 h at room temperature. Cells were imaged using a Zeiss LSM700 confocal microscope and analyzed using Fiji.

### Bulk RNA-sequencing and assignment of putative cellular sources for differentially- expressed transcripts

Total cellular RNA was extracted from P10 and P14 WT and *Apcdd1*^-/-^ retinas using QIAGEN RNeasy Mini Kit. 8 retinas from 4 pups were combined for each sample in each genotype. Concentrations of isolated RNA samples was ascertained using Nanodrop Spectrophotometer (ND-100). RNA samples were sent to either New York Genome Center (P10) or Genewiz (P14) for sequencing. cDNA libraries were prepared from RNA samples and sequenced to generate read counts for individual RNA species. Differential expression analyses were performed for these RNAs using the Bioconductor package DESeq2 in R. Cutoffs of the Benjamini-Hochberg adjusted p values (padj) ≤0.05 and |log_2_ fold change| ≥0.5 were used to identify statistically significant, differentially-expressed genes. Putative cellular sources for differentially expressed transcripts between P10 or P14 *Apcdd1*^-/-^ versus WT retinas were determined from the existing retinal single cell RNA sequencing gene expression database (https://singlecell.broadinstitute.org/single_cell/study/SCP301/c57b6-wild-type-p14-retina-by-drop-seq#study-summary) (Macosko et al., 2015). Any gene showing mean expression and percentage of a specific cell type expressing that gene as >0 was assigned to that specific cell type. Genes that showed expression in multiple retinal cell types in the database were assigned to all the cell types they were expressed in. Volcano plots for each gene category were constructed using the R software package and the top upregulated and downregulated genes as well as other specific genes of interest within each plot were identified (see **Supplemental Data Sheet 1-5**).

### *In vivo* tracer injection

Mice were anesthetized using isoflurane and injected with 1% Biocytin-5(and-6-)- Tetramethylrhodamine (Biocytin-TMR: ThermoFisher Scientific; Waltham, MA, Cat# T12921). After 30 minutes in circulation, mice were re-anesthetized and perfused with 1X phosphate buffer saline (PBS) via the cardiac route. Retinas and livers were isolated and fixed in 4% paraformaldehyde (PFA) for 2 hours prior to immunostaining with FITC-conjugated Lectin. Biocytin-TMR fluorescence in the livers of corresponding animals were used as internal controls for measurement of biocytin-TMR fluorescence in the retina as described (Lengfeld et al., 2017; Mazzoni et al., 2017).

### Immunofluorescence

Unless otherwise specified, isolated tissues (retina, brain and liver) were fixed in 4% PFA for 2 hours for immunostaining. The exceptions were retinal flat-mount staining for Laminin chains and Integrin subunits, in which cases retinas were fixed for 10 minutes in 2% PFA. For retinal flat-mounts, tissues were washed with 1X PBS following PFA fixation and blocked overnight in blocking buffer (5% goat or donkey serum, 0.3% Triton X-100 in 1× PBS) at 4°C. Samples were then incubated with primary antibodies in antibody-diluting solution (5% goat or donkey serum; 0.01% Triton X-100 in 1X PBS) for 48h at 4°C, washed, and incubated with secondary antibodies for 24 h. For radial section staining, samples were washed in 1 × PBS, cryoprotected in 30% sucrose overnight, embedded in optimal cutting temperature compound (Tissue-Tek; Torrance, CA, USA), and sectioned at a thickness of 12 µm. Tissue sections were blocked for 2 h at room temperature followed by primary antibody incubation overnight at 4°C. Following 1X PBS washes, sections were incubated with secondary antibodies for 4 h at room temperature. Samples were imaged in room temperature using a Zeiss LSM700 confocal microscope [ZEISS 20X air (<u>Objective Plan-Apochromat 20x/0.8 M27</u>; numerical aperture: 0.8) or 40X water (<u>Objective C-Apochromat 40x/1.2 W autocorr M27</u>; numerical aperture: 1.2) objectives; ZEN image acquisition software] and analyzed using Fiji software as described before (Mazzoni et al., 2017).

### Antibodies used in immunofluorescence

Primary antibodies used were rat anti-CD31 (1:250; BD Biosciences; San Jose, CA; Cat# 553370, RRID:AB_394816), rabbit anti-CD31 (1:250; Abcam; Cambridge, MA; Cat# AB28364, RRID:AB_726362), goat anti-Pdgfrβ (1:200; R&D Systems; Minneapolis, MN; Cat# AF1042, RRID:AB_2162633), FITC-conjugated rabbit anti-LEF1 (1:100; Cell Signaling Technology; Danvers, MA; Cat# 2230s, RRID:AB_823558), rabbit anti-GFP (1:500; ThermoFisher Scientific; Waltham, MA; Cat# A11122, RRID:AB_221569), rabbit anti-NG2 (1:250; Millipore; Burlington, MA; Cat# AB5320, RRID:AB_11213678), goat anti-β-galactosidase (1:250; Abcam; Cambridge, MA; Cat# AB12081, RRID:AB_725684), rat anti-Laminin α2 (1:200; Abcam; Cambridge, MA; Cat# AB11576, RRID:AB_298180), rat anti-Laminin β1 (1:200; Abcam; Cambridge, MA; Cat# AB44941, RRID:AB_775971), rabbit anti-Laminin β2 (1:1000; a gift from Dr. William J Brunken; SUNY Upstate Medical University, Syracuse, NY), rabbit anti-GFAP (1:1000; Millipore; Burlington, MA, Cat# AB5804, RRID:AB_2109645), rat anti-Pdgfrα (1:100; BD Biosciences; San Jose, CA; Cat# 558774, RRID:AB_397117), rabbit anti-Aquaporin-4 (1:250; Sigma-Aldrich; St. Louis, MO; Cat# A5971, RRID:AB_258270), rat anti-Integrin α6 (1:100; Millipore; Burlington, MA; Cat# MAB1982, RRID:AB_2128296) and FITC-conjugated mouse anti-Integrin α2 (1:100; Santa-Cruz Biotechnology; Dallas, TX; Cat# sc-74466, RRID:AB_1124939). Secondary antibodies used were goat anti-rat AlexaFluor 594, goat anti-rabbit AlexaFluor 594, goat anti-rat AlexaFluor 488, goat anti-rabbit AlexaFluor 488, donkey anti-rat AlexaFluor 594, donkey anti-rabbit AlexaFluor 594, donkey anti-rat AlexaFluor 488, donkey anti-rabbit AlexaFluor 488, donkey anti-rat AlexaFluor 647, donkey anti-rabbit AlexaFluor 647, donkey anti-goat AlexaFluor 594 and donkey anti-goat AlexaFluor 488 (1:250; ThermoFisher Scientific; Waltham, MA). Other reagents used were Fluorescein *Griffonia simplicifolia*-Lectin I (Isolectin B4; 1:250; Vector Laboratories; Burlingame, CA; Cat# FL-1201, RRID:AB_2314663).

### RNA *in situ* hybridization

Antisense mRNA probe for *Lama2* (Laminin α2) was prepared using TOM6004-MGC premier ORF clone for *Lama2* (Transomic Technologies; Huntsville, AL; Clone ID: BC172647, Accession: BC172647). Preparation and validation of antisense mRNA probes for full length *Apcdd1* and *Pdgfr*β was described before(Mazzoni et al., 2017). Fluorescent *in situ* hybridizations using these probes were performed as described before(Mazzoni et al., 2017). In case of fluorescent *in situ* hybridization along with immunolabeling with an endothelial or mural cell marker, the samples were immunostained for Caveolin-1 (1:500; Abcam; Cambridge, MA; Cat# AB18199, RRID:AB_444314) or NG2, respectively, following *in situ* hybridization for *Apcdd1* or *Lama2*. The samples were imaged using Zeiss LSM700 confocal microscope and analyzed using Fiji software.

### Western blotting

Retinas were collected from mice after perfusion with PBS. For each sample, 8 retinas from four mice were pulled together. Collected tissue was homogenized in lysis buffer containing protease and phosphatase inhibitors using a Dounce homogenizer followed by sonication. Protein levels in P10, P14 and P18 retinas were measured by fluorescent Western blot analysis [4-15% or 7.5% Mini-PROTEAN TGX gels (Bio-Rad)] and quantitation was performed using the LICOR (LICOR, Lincoln, NE) system, as described(Mazzoni et al., 2017). Western blot analysis used the following primary antibodies: rat anti-Laminin α2 (1:200; Abcam; Cambridge, MA; Cat# AB11576, RRID:AB_298180), rabbit anti-Laminin (1:2500; Sigma-Aldrich; St. Louis, MO; Cat# L9393, RRID:AB_477163; primarily recognizes Lama1), rabbit anti-Aquaporin-4 (1:250; Sigma-Aldrich; St. Louis, MO; Cat# A5971, RRID:AB_258270) and anti-β-actin (1:10000; Novus Biologicals; Centennial CO; Cat# NB600-501, RRID:AB_10077656). IRDyes 680 and 800 (1:20000; LI-COR) were used as secondary antibodies. Fluorescent quantification of protein levels was done using the Odyssey SA infrared imaging system. Values are displayed as protein levels corrected with β-actin and normalized to wild-type control values.

### Image acquisition and analysis

For retinal flat-mounts, each measurement was made in at least 3 quadrants per sample. For sections, at least three sections were imaged per sample. In each case, multiple representative images were acquired and averaged. To measure the number of Lef1^+^ and nGFP^+^ mural cells (**Fig. 1**), area positive for Pdgfrβ was measured from each image in Fiji software (National Institutes of Health, Bethesda, MD, USA) using signal masking and the number of Lef1^+^ and nGFP^+^ nuclei within that Pdgfrβ^+^ area was quantified and normalized to unit area. To measure *Apcdd1* expression in ECc and mural cells (**Fig. 2**), number of Apcdd1^+^/Pdgfrβ^+^ or Apcdd1^+^/Cav1^+^ cells were quantified and presented as the percentage of total Pdgfrβ^+^ or Cav1^+^ cells, respectively. To measure *Lama2* mRNA expression in mural cells (**Fig. 3**), mean fluorescence intensity (M.F.I) of *Lama2* puncta in individual mural cells were calculated with histogram analysis in Fiji software. To measure deposition of Laminins in the vBM (**Fig. 3**, Sup. Fig. 3, 5), mean fluorescence intensity (M.F.I) ratios (Laminin:CD31) in individual vessel segments were calculated with histogram analysis in Fiji software. Data were presented as values normalized to the WT average values. To measure Lef1 expression *in vitro* (**Sup. Fig. 4**), the number of Lef1^+^ nuclei were quantified as the percentage of total nuclei. To measure laminin deposition *in vitro* (**Sup. Fig. 4**), Laminin M.F.I was quantified from each image and normalized to the number of total nuclei. The intensity of biocytin-TMR outside of retinal vessels (tracer leakage; **Fig. 4**) was measured as described (Mazzoni et al., 2017). Biocytin-TMR signal intensities in the retina and cerebella were normalized to those of livers. To measure the relative expression of GFAP (**Fig. 5**), the M.F.I were calculated for each field and data was presented as values normalized to the WT. To measure Aqp-4^+^ astrocyte end-feet coverage of arteries and veins (**Fig. 5**), percentage of total vessel length surrounded by Aqp-4^+^ astrocyte end-feet in individual arterial and venous segments were calculated. To measure astrocytic expression of Integrin α6 (**Fig. 6**), M.F.I ratios (Integrin α6/Gfap) in each field was calculated. Data were presented as values normalized to the WT average values.

### Statistical analyses

Samples from 3-7 different animals (for *in vivo* experiments) and 3-4 separate experiments (for *in vitro* experiments) were used for statistical analysis. To test for statistical significance of any differences, either one-way ANOVA or 2-tailed, unpaired Student’s *t* test was performed. A value of *P <* 0.05 was considered statistically significant.

## SUMMARY OF SUPPLEMENTAL MATERIALS

Sup. Fig. 1 shows that Norrin/β-catenin signaling is activated in retinal and cerebellar ECs and vascular mural cells, but not in retinal astrocytes. Sup. Fig. 2 shows that Lama2 (*Lama2*) is predominantly expressed in retinal PCs, but not in ECs, and is upregulated in the *Apcdd1*^-/-^ retina. Sup. Fig. 3 shows that Mural cell-derived Lama2 expression and deposition are upregulated in the *Apcdd1^-/-^* cerebellar vascular BM. Sup. Fig. 4 shows that Lama2 expression is upregulated in primary brain PCs, but not in ECs, upon Norrin/β-catenin activation. Sup. Fig. 5 shows that Laminin-211 is likely the most affected isoform in the *Apcdd1^-/-^* retina. Sup. Fig. 6 shows decreased astrocyte end-feet polarization in the *Lama2^-/-^* retina. Sup. Fig. 7 shows increased Aqp-4 expression in astrocyte end-feet and astrocytic Integrin α6 expression in the *Apcdd1^-/-^* cerebella. Sup. data sheet 1 list of differentially expressed genes (significant and non-significant) between wild-type and *Apcdd1*^-/-^ retinas at P10 (P10 Raw data) and P14 (P14 Raw Data). Sup. data sheet 2 lists significant differentially expressed putative endothelial genes between wild-type and *Apcdd1*^-/-^ retinas at P10 and P14. Sup. data sheet 3 lists significant differentially expressed putative pericyte genes between wild-type and *Apcdd1*^-/-^ retinas at P10 and P14. Sup. data sheet 4 lists significant differentially expressed extracellular matrix (ECM) genes between wild-type and *Apcdd1*^-/-^ retinas at P10 and P14. Sup. data sheet 5 lists significant differentially expressed astrocyte maturity genes between wild-type and *Apcdd1*^-/-^ retinas at P10 and P14.

## Supporting information

Supplementary figures 1-6 and data sheets 1-5

## ACKNOWLEDGEMENT

Saptarshi Biswas, Tyler Cutforth and Dritan Agalliu conceived the project. Saptarshi Biswas, Sanjid Shahriar, Nicholas P. Giangreco, Panos Arvanitis and Markus Winkler performed the experiments and analyses. Saptarshi Biswas, Sanjid Shahriar, Nicholas P. Tatonetti, Tyler Cutforth, William J Brunken and Dritan Agalliu wrote the manuscript. Saptarshi Biswas, Sanjid Shahriar, Nicholas P. Giangreco and Dritan Agalliu prepared figures and figure legends.

## COMPETING INTERESTS

The authors declare that the research was conducted without any commercial or financial relationships that can be construed as potential conflicts of interest.

## FUNDING

Saptarshi Biswas, Sanjid Shahriar, Tyler Cutforth and Dritan Agalliu are supported by grants from the NIH/NIMH (National Institute of Health/National Institute of Mental Health; R01 MH112849), NIH/NINDS (National Institute of Health/National Institute of neurological Disorders and Stroke; R01 NS107344), the National Multiple Sclerosis (MS) Society (RG-1901-33218) and the Leducq Foundation (15CDV-02), and Tyler Cutforth and Dritan Agalliu are partially supported by unrestricted gifts from Newport Equities LLC (Limited Liability Company) and PANDAS Network to the Division of Cerebrovascular Diseases and Stroke, Department of Neurology, Columbia University Irving Medical center. William. J. Brunken is supported by grants from the NIH/NEI (National Institute of Health/National Eye Institute; R01-Ey12676) and in part by funds from an unrestricted grant from Research to Prevent Blindness to the Department of Ophthalmology and Visual Sciences, Upstate Medical University. Nicholas P. Giangreco and Nicholas P. Tatonetti are supported by a grant from the NIH/NIGMS (National Institute of Health/National Institute of General Medical Sciences; R35GM131905).

## DATA AVAILABILITY

C57B6 wild-type P14 retina by drop-seq (Macosko et al., 2015); DOI: 10.1016/j.cell.2015.05.002; (Link: https://singlecell.broadinstitute.org/single_cell/study/SCP301/c57b6-wild-type-p14-retina-by-drop-seq#study-summary)

Database of gene expression in adult mouse brain and lung vascular and perivascular cells (He et al., 2018; Vanlandewijck et al., 2018); DOI: 10.1038/sdata.2018.160; DOI: 10.1038/nature25739; (Link: http://betsholtzlab.org/VascularSingleCells/database.html)

## MATERIALS AND CORRESPONDENCE

The materials are available upon request to Dritan Agalliu.

## Notes

https://singlecell.broadinstitute.org/single_cell/study/SCP301/c57b6-wild-type-p14-retina-by-drop-seq#study-summary

http://betsholtzlab.org/VascularSingleCells/database.html

### Competing Interest Statement

The authors have declared no competing interest.

