## Supplementary figures 1-6 and data sheets 1-5 for "Mural norrin/β-catenin signaling regulates Lama2 expression to promote neurovascular unit assembly"

Mailing address:
Columbia University Medical Center
650 West 168th Street
Black Building Room 310
New York, NY 10032

ORCID: 0000-0002-5375-4143

**RUNNING TITLE**

Mural Wnt signaling regulates Lama2

#### SUPPLEMENTARY FIGURES AND FIGURE LEGENDS

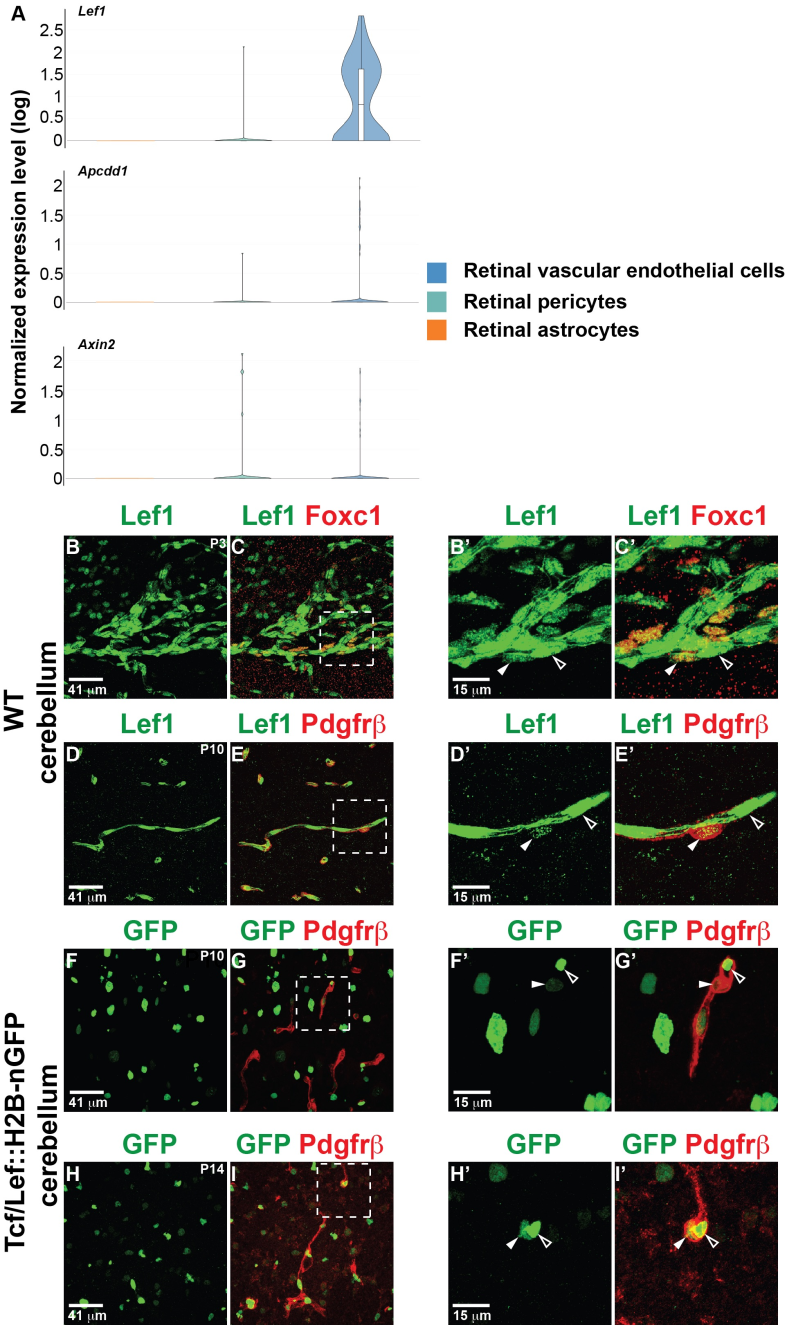


**Supplementary Fig. 1 (related to Figs. 1, 2). Norrin/β-catenin signaling is activated in CNS endothelial and mural cells, but not in astrocytes. A)** Analyses of a P14 WT retinal single cell RNAseq database (see Results and Materials and Methods sections) to explore expression profiles of downstream Norrin/β-catenin targets (*Lef1*, *Apcdd1*, *Axin2*) in endothelial cells (ECs), pericytes (PCs) and astrocytes. **B-C')** Postnatal day (P) 3 wild-type (WT) cerebellar sections were stained for Lef1 (green) and Foxc1 [red; immature PC marker]. **D-E')** P10 WT cerebellar sections were stained for Lef1 (green) and Pdgfrβ (red; PC marker). **F-I')** P10 and P14 Tcf/Lef::H2B-nGFP cerebellar sections were stained for GFP (green) and Pdgfrβ (red). In all images empty arrowheads point to either Lef1$^+$ or GFP$^+$ ECs, whereas solid arrowheads point to either Lef1$^+$ or GFP$^+$ mural cells. Scale bars: B-I = 41 μm; B'-I' = 20 μm.

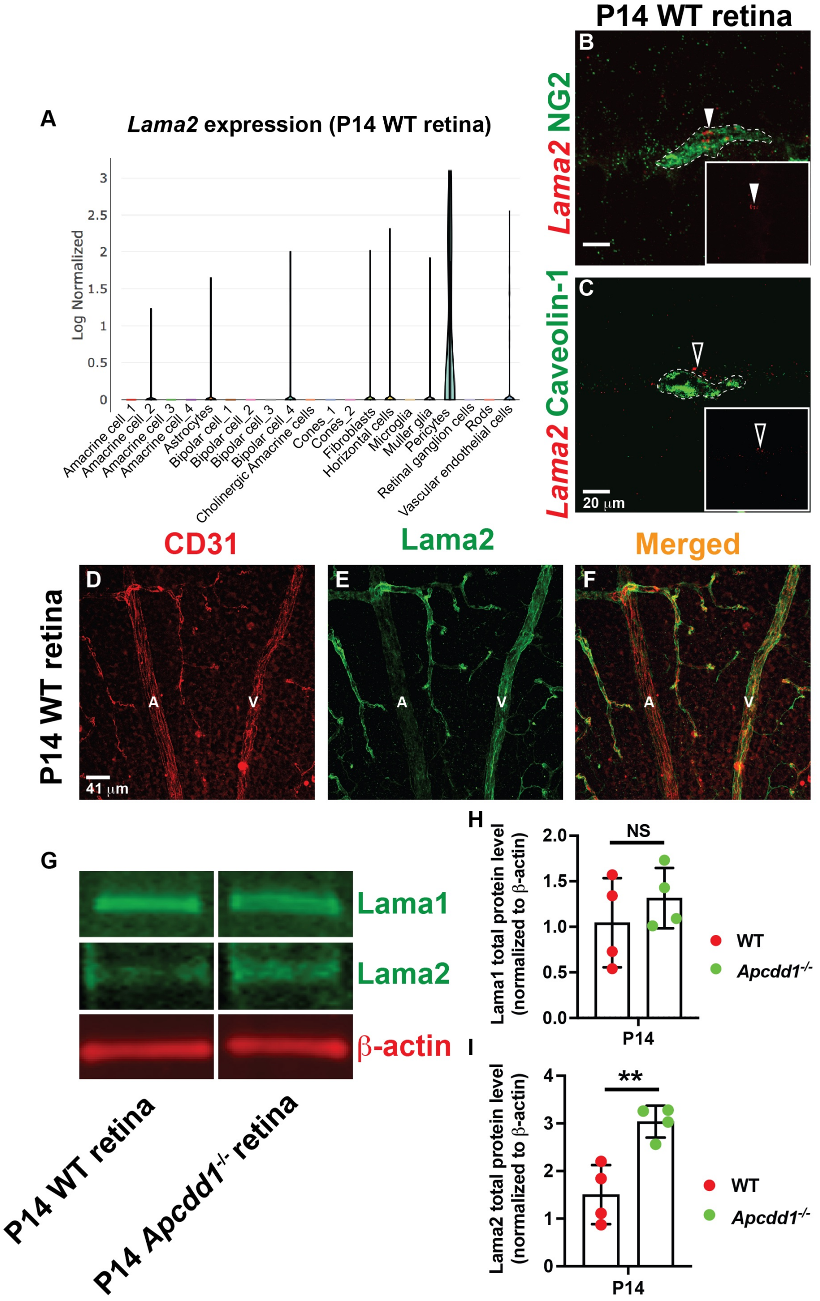


**Supplementary Fig. 2 (related to Fig. 3). Lama2 (*Lama2*) is predominantly expressed in retinal PCs, but not in ECs, and is upregulated in the *Apcdd1*$^{-/-}$ retina. A)** Expression profile of *Lama2* in P14 WT retinal cells from the published P14 retina single cell RNAseq database. **B)** Fluorescent *in situ* hybridization of P14 WT retinal section with antisense probes against *Lama2* (red) and immunolabeling with NG2 (green; mural cell marker). Solid arrowheads point to *Lama2* expression in mural cells (Inset: *Lama2* single channel image). **C)** Fluorescent *in situ* hybridization of P14 WT retinal section with antisense probes against *Lama2* (red) and immunolabeling with Caveolin-1 (green). Empty arrowheads point to *Lama2* expression outside ECs (Inset: *Lama2* single channel image). **D-F)** P14 WT retinal flat mounts were stained for CD31 (red) and Lama2 (green). A = artery, V = vein. **G)** P14 WT and *Apcdd1*$^{-/-}$ whole retinal lysates were probed for Lama1 or Lama2 (green bands) and β-actin (red bands) by western blot. **H-I)** Quantification of Lama1 and Lama2 protein levels (normalized to β-actin) by western blotting (n=4 samples/ group). Student's t-test **p<0.02, NS: not significant. Error bars: SD. Scale bars: B, C = 20 μm; D-F = 41 μm.

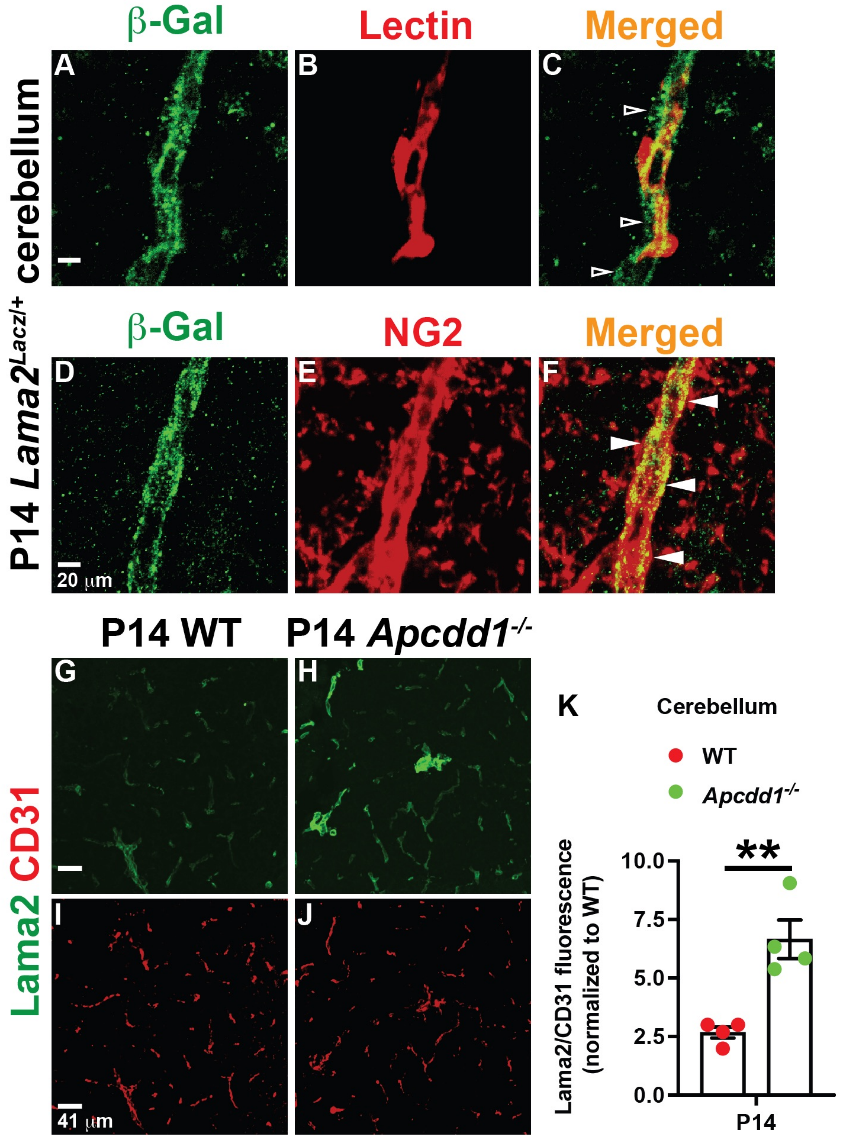

**Supplementary Fig. 3 (related to Fig. 3). Mural cell-derived Lama2 expression and deposition are upregulated in the *Apcdd1$^{-/-}$* cerebellar vascular basement membrane**. **A-F)** P14 *Lama2$^{Lacz/+}$* cerebellar sections were stained for β-galactosidase (β-gal; green) and either Lectin (**A-C;** red) or NG2 (**D-F;** red). Empty arrowheads point to β-Gal$^{-}$ ECs. Solid arrowheads point to β-Gal$^{+}$ mural cells. **G-J)** P14 WT and *Apcdd1$^{-/-}$* cerebellar sections were immunolabelled for Lama2 (green) and CD31 (red). **K)** Quantification of the ratio of Lama2 / CD31 mean fluorescence intensity (M.F.I) in P14 blood vessels, normalized to the WT values (n=4 mice/group); Students' t-test **p<0.02. Error bars: SEM. Scale bars: **A-F** = 20 µm; **G-J** = 41 µm.

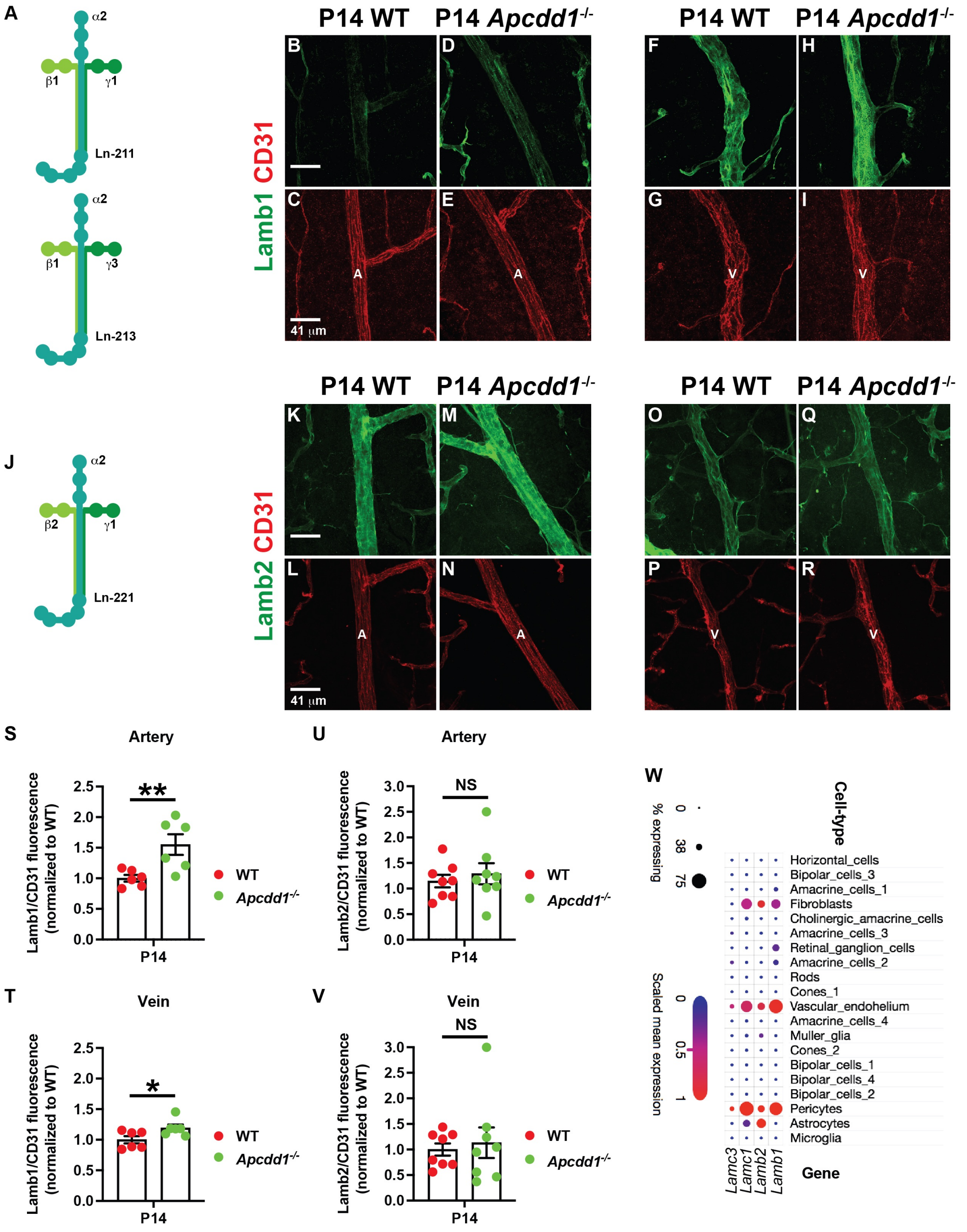


**Supplementary Fig. 4 (related to Fig. 3). Laminin-211 is likely the most affected isoform in the *Apcdd1*$^{-/-}$ retina. A)** Schematic diagram of two Laminin heterotrimers (211 and 213) containing α2 and β1. **B-I)** P14 WT and *Apcdd1*$^{-/-}$ retinal flat-mounts were labelled for Lamb1 chain (green) and CD31 (red). **J)** Schematic diagram of one α2- and β2-containing Laminin heterotrimer (221). **K-R)** P14 WT and *Apcdd1*$^{-/-}$ retinal flat-mounts were labelled for Lamb2 chain (green) and CD31 (red). **S, T)** Quantification of the ratio of Lamb1/CD31 fluorescence intensities in arteries and veins at P14 normalized to the WT average values (6 arteries or veins analyzed from n=3 mice/group). **U, V)** Quantification of the ratio of Lamb2/CD31 fluorescence intensities in arteries and veins at P14 normalized to the WT average values (8 arteries and 7 veins analyzed from n=4 mice/group). A: artery, V: vein. **W)** Analyses of the published P14 WT retinal single cell RNAseq database (see Results and Materials and Methods sections) to explore the expression profiles of *Lamb1*, *Lamb1*, *Lamc1* and *Lamc3*. Student's t-test *p<0.05, **p<0.02, NS: not significant. Error bars: SEM. Scale bars: 41 μm.

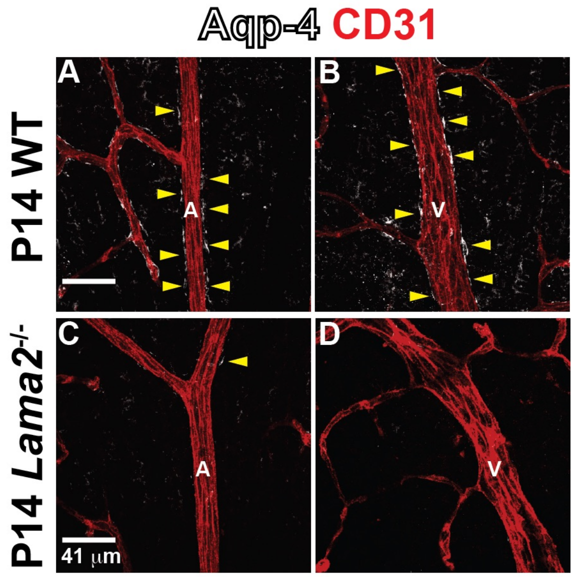


**Supplementary Fig. 5 (related to Fig. 6). Decreased astrocyte endfeet polarization in the *Lama2*$^{-/-}$ retina. A-D)** P14 WT and *Lama2*$^{-/-}$ retinal flat-mounts were labelled for Aqp-4 (white) and CD31 (red). Yellow arrowheads point at Aqp-4$^{+}$ astrocyte endfeet around blood vessels in the retina. A: arteries and V: veins. Scale bars = 41 μm.

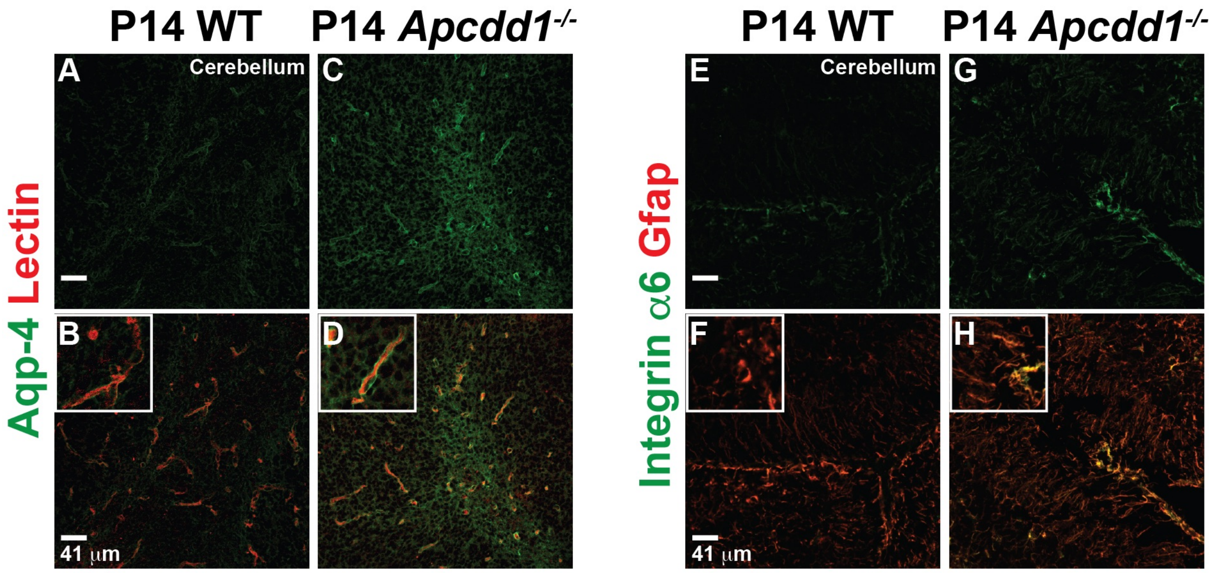


**Supplementary Fig. 6 (related to Figs. 6 and 7): Increased Aqp-4 expression in astrocyte endfeet and astrocytic Integrin α6 expression in the *Apcdd1*$^{-/-}$ cerebella. A-D)** P14 WT and *Apcdd1*$^{-/-}$ cerebellar sections were labelled for Aqp-4 (green) and Lectin (red). Insets show higher magnification images of single vessels. **E-H)** P10 WT and *Apcdd1*$^{-/-}$ cerebellar sections were labelled for Integrin α6 (green) and GFAP (red). Insets show higher magnification images. Scale bar = 41 μm.

**Supplementary data sheet 1 (related to Figs. 2, 3, 6 and 7)**. List of significantly differentially expressed genes between wild-type and *Apcdd1*$^{-/-}$ retinas at P10 and P14. Negative values mean that the gene is lower in *Apcdd1*$^{-/-}$ compared to wild-type retinas and positive values mean that the gene is higher in *Apcdd1*$^{-/-}$ compared to wild-type retinas.

**Supplementary data sheet 2 (related to Fig. 2)**. List of significant differentially expressed putative endothelial genes between wild-type and *Apcdd1*$^{-/-}$ retinas at P10 and P14. The genes are ranked by $log_2$Fold Change from the lowest to the highest value. Negative values mean that the gene is lower in *Apcdd1*$^{-/-}$

compared to wild-type retinas and positive values mean that the gene is higher in *Apcdd1*$^{-/-}$ compared to wild-type retinas.

**Supplementary data sheet 3 (related to Figs. 2 and 3)**. List of significant differentially expressed putative pericyte genes between wild-type and *Apcdd1*$^{-/-}$ retinas at P10 and P14. The genes are listed by $log_2$Fold Change from the lowest to the highest value. Negative values mean that the gene is lower in *Apcdd1*$^{-/-}$ compared to wild-type retinas and positive values mean that the gene is higher in *Apcdd1*$^{-/-}$ compared to wild-type retinas.

**Supplementary data sheet 4 (related to Figs. 2, 3 and 7)**. List of significant differentially expressed extracellular matrix genes (ECM) between wild-type and *Apcdd1*$^{-/-}$ retinas at P10 and P14. The genes are listed by $log_2$Fold Change from the lowest to the highest value. Negative values mean that the gene is lower in *Apcdd1*$^{-/-}$ compared to wild-type retinas and positive values mean that the gene is higher in *Apcdd1*$^{-/-}$ compared to wild-type retinas.

**Supplementary data sheet 5 (related to Fig. 6)**. List of significant differentially expressed astrocyte maturity genes between wild-type and *Apcdd1*$^{-/-}$ retinas at P10 and P14. The genes are listed by $log_2$Fold Change from the lowest to the highest value. Negative values mean that the gene is lower in *Apcdd1*$^{-/-}$ compared to wild-type retinas and positive values mean that the gene is higher in *Apcdd1*$^{-/-}$ compared to wild-type retinas.

#### Supplementary data sheet 1

**Significantly differentially expressed genes (P10 Apcdd1-/- vs Wt retina)**

| Gene name | Ensembl_ID | log2FoldChange | padj |
|---|---|---|---|
| Gm6563 | ENSMUSG00000051255.5 | -0.502134576 | 0.00755658 |
| Prr19 | ENSMUSG00000058741.3 | -0.500490366 | 0.00731154 |
| Slc37a2 | ENSMUSG00000032122.11 | -0.507351488 | 0.00639986 |
| A830052D11Rik | ENSMUSG00000097413.1 | -0.513085712 | 0.00610216 |
| Ccdc175 | ENSMUSG00000021086.4 | 0.50581448 | 0.00590605 |
| Kynu | ENSMUSG00000026866.13 | 0.51085217 | 0.00590605 |
| Marc1 | ENSMUSG00000026621.10 | 0.504203607 | 0.00577374 |
| Cfap58 | ENSMUSG00000046585.8 | -0.518345122 | 0.00553964 |
| Kcnh4 | ENSMUSG00000035355.12 | -0.520070847 | 0.00536188 |
| Hpx | ENSMUSG00000030895.8 | 0.511147457 | 0.00523578 |
| Mtif3 | ENSMUSG00000016510.8 | 0.515350411 | 0.00510159 |
| Gm10031 | ENSMUSG00000101523.1 | -0.507369603 | 0.00452498 |
| Ccr1 | ENSMUSG00000025804.4 | -0.515623594 | 0.00423573 |
| Mss51 | ENSMUSG00000021815.3 | -0.53247028 | 0.0040759 |
| 4732414G09Rik | ENSMUSG00000086943.2 | -0.511597957 | 0.00403714 |
| Cyp4f15 | ENSMUSG00000073424.6 | -0.532144786 | 0.00378494 |
| Fam184b | ENSMUSG00000015879.8 | 0.50365433 | 0.00377965 |
| Rec114 | ENSMUSG00000074269.7 | -0.511656783 | 0.00377965 |
| Obox6 | ENSMUSG00000041583.8 | 0.54167456 | 0.00329751 |
| Gpc2 | ENSMUSG00000029510.12 | -0.53040727 | 0.00318171 |
| Fmo2 | ENSMUSG00000040170.10 | 0.518539501 | 0.00294134 |
| Klhdc7a | ENSMUSG00000078234.6 | -0.540045307 | 0.00288449 |
| Lrrc23 | ENSMUSG00000030125.8 | -0.506381095 | 0.00275934 |
| Cntd1 | ENSMUSG00000078653.4 | -0.548503477 | 0.00273465 |
| Paox | ENSMUSG00000025464.11 | 0.515487464 | 0.00228097 |
| Tfb1m | ENSMUSG00000036983.6 | -0.520045041 | 0.00221225 |
| Gdpd3 | ENSMUSG00000030703.7 | -0.557088638 | 0.00218732 |
| Nat1 | ENSMUSG00000025588.3 | -0.550123941 | 0.00214166 |
| Mas1 | ENSMUSG00000068037.7 | -0.541068873 | 0.00212712 |
| Olfml2a | ENSMUSG00000046618.7 | -0.558525393 | 0.0020246 |
| Gyltl1b | ENSMUSG00000040434.13 | -0.557600523 | 0.00197267 |
| Cthrc1 | ENSMUSG00000054196.6 | -0.505566406 | 0.00185974 |
| Clspn | ENSMUSG00000042489.12 | -0.523881538 | 0.0017353 |
| Gm10320 | ENSMUSG00000092116.1 | 0.56879311 | 0.00173513 |
| Proca1 | ENSMUSG00000044122.11 | -0.542594545 | 0.00168878 |
| Glo1 | ENSMUSG00000024026.9 | -0.567615031 | 0.00168522 |
| Aspdh | ENSMUSG00000038704.7 | -0.565314656 | 0.00157255 |
| 9130227L01Rik | ENSMUSG00000099950.1 | 0.543254546 | 0.0014652 |
| Dmbx1 | ENSMUSG00000028707.12 | -0.545653083 | 0.00119043 |
| Arhgef33 | ENSMUSG00000054901.5 | -0.509338923 | 0.00110588 |
| Fam181a | ENSMUSG00000096753.4 | 0.579424257 | 0.00110584 |
| Csrnp1 | ENSMUSG00000032515.7 | 0.561867851 | 0.00106739 |

| | | | |
|---|---|---|---|
| Prss38 | ENSMUSG00000049291.6 | -0.500575561 | 0.00106665 |
| Palm3 | ENSMUSG00000047986.9 | 0.519732332 | 0.00104406 |
| Spef2 | ENSMUSG00000072663.8 | -0.505044659 | 9.89E-04 |
| Tspan11 | ENSMUSG00000030351.4 | 0.587390452 | 8.80E-04 |
| Ankrd23 | ENSMUSG00000067653.8 | -0.575135622 | 8.73E-04 |
| Arhgef39 | ENSMUSG00000051517.11 | -0.59500779 | 8.73E-04 |
| Csf3r | ENSMUSG00000028859.11 | -0.504144961 | 7.95E-04 |
| Cr2 | ENSMUSG00000026616.11 | -0.59135973 | 7.80E-04 |
| A830019P07Rik | ENSMUSG00000094707.1 | 0.593644258 | 7.29E-04 |
| Gm9913 | ENSMUSG00000053615.1 | -0.578050597 | 6.99E-04 |
| Cwc22 | ENSMUSG00000027014.11 | 0.527087346 | 6.99E-04 |
| D330050G23Rik | ENSMUSG00000085316.1 | -0.567367287 | 6.66E-04 |
| Scrg1 | ENSMUSG00000031610.3 | -0.593423015 | 6.43E-04 |
| Hdhd3 | ENSMUSG00000038422.2 | -0.558639635 | 6.02E-04 |
| Npy4r | ENSMUSG00000048337.3 | -0.531717427 | 5.88E-04 |
| Pde6a | ENSMUSG00000024575.12 | 0.537561876 | 5.14E-04 |
| Rab32 | ENSMUSG00000019832.3 | -0.599232353 | 4.81E-04 |
| Slc29a1 | ENSMUSG00000023942.12 | 0.622051 | 4.47E-04 |
| Slco1a5 | ENSMUSG00000063975.10 | 0.60385568 | 4.44E-04 |
| Cfi | ENSMUSG00000058952.9 | 0.564369837 | 4.01E-04 |
| Lct | ENSMUSG00000026354.8 | -0.574149308 | 3.94E-04 |
| Poteg | ENSMUSG00000063932.4 | -0.575400961 | 3.93E-04 |
| Cntn3 | ENSMUSG00000030075.8 | -0.504108614 | 3.78E-04 |
| Tekt1 | ENSMUSG00000020799.13 | 0.606824802 | 3.53E-04 |
| Gm6471 | ENSMUSG00000073781.3 | -0.630870089 | 3.48E-04 |
| Entpd3 | ENSMUSG00000041608.7 | 0.546849483 | 3.47E-04 |
| Rsph9 | ENSMUSG00000023966.5 | -0.548567717 | 3.36E-04 |
| Cd1d2 | ENSMUSG00000041750.10 | 0.588685848 | 3.36E-04 |
| 2010107G12Rik | ENSMUSG00000029847.10 | -0.542367984 | 3.30E-04 |
| Rab11fip1 | ENSMUSG00000031488.10 | -0.595587245 | 3.27E-04 |
| Itga2 | ENSMUSG00000015533.8 | 0.574918927 | 3.19E-04 |
| Zfp541 | ENSMUSG00000078796.3 | -0.567307023 | 3.12E-04 |
| Pmfbp1 | ENSMUSG00000031727.7 | 0.518814566 | 3.09E-04 |
| Postn | ENSMUSG00000027750.13 | -0.543828886 | 2.39E-04 |
| Best1 | ENSMUSG00000037418.5 | -0.536555626 | 2.39E-04 |
| Zfp868 | ENSMUSG00000060427.12 | -0.603405255 | 2.34E-04 |
| Oscar | ENSMUSG00000054594.10 | 0.620657121 | 2.10E-04 |
| Adamts19 | ENSMUSG00000053441.4 | -0.638591756 | 2.06E-04 |
| Ermap | ENSMUSG00000028644.13 | -0.633234357 | 1.65E-04 |
| Unc93a | ENSMUSG00000067049.7 | -0.535613729 | 1.63E-04 |
| RP23-363M4.1 | ENSMUSG00000105039.1 | 0.650039433 | 1.61E-04 |
| Psg29 | ENSMUSG00000023159.7 | -0.641232349 | 1.49E-04 |
| Gm26779 | ENSMUSG00000097140.1 | -0.646382264 | 1.46E-04 |
| Ccdc114 | ENSMUSG00000040189.12 | -0.648012128 | 1.36E-04 |
| C030034I22Rik | ENSMUSG00000073374.3 | -0.621455268 | 1.25E-04 |

| | | | |
|---|---|---|---|
| Efhc1 | ENSMUSG00000041809.5 | -0.642002398 | 1.24E-04 |
| F730016J06Rik | ENSMUSG00000086425.4 | -0.544476101 | 1.21E-04 |
| Trim17 | ENSMUSG00000036964.11 | 0.557900847 | 1.13E-04 |
| Tnfsf13 | ENSMUSG00000089669.4 | -0.522757812 | 1.12E-04 |
| Dsg2 | ENSMUSG00000044393.12 | -0.655643633 | 9.87E-05 |
| Calml4 | ENSMUSG00000032246.10 | 0.529095506 | 9.36E-05 |
| Ccdc24 | ENSMUSG00000078588.7 | -0.658904859 | 9.34E-05 |
| Il12rb2 | ENSMUSG00000018341.9 | -0.554418421 | 8.96E-05 |
| Skor2 | ENSMUSG00000091519.1 | -0.621550075 | 8.95E-05 |
| Pomc | ENSMUSG00000020660.5 | 0.591905084 | 8.81E-05 |
| Col17a1 | ENSMUSG00000025064.11 | -0.662756196 | 8.40E-05 |
| B3gnt3 | ENSMUSG00000031803.7 | 0.580885549 | 8.08E-05 |
| Slc25a41 | ENSMUSG00000011486.11 | 0.677771828 | 7.74E-05 |
| Eva1c | ENSMUSG00000039903.13 | 0.53885811 | 7.69E-05 |
| Gm13154 | ENSMUSG00000065999.10 | -0.545652494 | 7.68E-05 |
| Scand1 | ENSMUSG00000046229.9 | 0.532850733 | 7.23E-05 |
| Mbl2 | ENSMUSG00000024863.5 | 0.503491946 | 7.23E-05 |
| Trim30d | ENSMUSG00000057596.10 | -0.638266854 | 7.06E-05 |
| Dctn3 | ENSMUSG00000028447.8 | -0.503775922 | 6.96E-05 |
| Gm15446 | ENSMUSG00000090015.5 | -0.546254959 | 6.57E-05 |
| Ncapg | ENSMUSG00000015880.10 | 0.686658145 | 6.49E-05 |
| 1600014C23Rik | ENSMUSG00000094690.1 | 0.683367738 | 6.08E-05 |
| Egln3 | ENSMUSG00000035105.5 | -0.614971745 | 5.91E-05 |
| Col20a1 | ENSMUSG00000016356.14 | -0.681575818 | 5.80E-05 |
| Lama2 | ENSMUSG00000019899.12 | 0.523332443 | 5.57E-05 |
| Pla2g2c | ENSMUSG00000028750.9 | -0.654897698 | 5.53E-05 |
| Mapk15 | ENSMUSG00000063704.9 | -0.680991762 | 4.98E-05 |
| Fyb | ENSMUSG00000022148.12 | -0.547277017 | 4.56E-05 |
| Gm2115 | ENSMUSG00000097789.1 | 0.513578578 | 4.41E-05 |
| Adra1d | ENSMUSG00000027335.9 | 0.558922969 | 4.29E-05 |
| Lcn2 | ENSMUSG00000026822.11 | 0.6348563 | 4.08E-05 |
| Cpne7 | ENSMUSG00000034796.11 | 0.52250988 | 3.53E-05 |
| Calcrl | ENSMUSG00000059588.10 | -0.567152965 | 3.14E-05 |
| Hs3st3a1 | ENSMUSG00000047759.6 | 0.643886902 | 2.69E-05 |
| Gfra3 | ENSMUSG00000024366.7 | 0.716455619 | 2.50E-05 |
| Tescl | ENSMUSG00000055826.4 | -0.560422728 | 2.24E-05 |
| Met | ENSMUSG00000009376.12 | -0.721766551 | 2.13E-05 |
| Hebp2 | ENSMUSG00000019853.5 | 0.554960147 | 1.97E-05 |
| Steap1 | ENSMUSG00000015652.6 | 0.726113222 | 1.93E-05 |
| Gm1698 | ENSMUSG00000074404.5 | -0.71350095 | 1.91E-05 |
| Ndufa4l2 | ENSMUSG00000040280.9 | 0.596559115 | 1.88E-05 |
| Acaca | ENSMUSG00000020532.15 | -0.514300121 | 1.77E-05 |
| Acot1 | ENSMUSG00000072949.5 | -0.599754484 | 1.69E-05 |
| Pcsk9 | ENSMUSG00000044254.6 | -0.641180299 | 1.66E-05 |
| Col24a1 | ENSMUSG00000028197.4 | -0.705233479 | 1.50E-05 |

| | | | |
|---|---|---|---|
| Podn | ENSMUSG00000028600.12 | -0.537021417 | 1.47E-05 |
| Gnmt | ENSMUSG00000002769.8 | 0.644758008 | 1.39E-05 |
| Kif20b | ENSMUSG00000024795.10 | -0.540782613 | 1.28E-05 |
| Hao2 | ENSMUSG00000027870.8 | 0.714855169 | 1.20E-05 |
| Fam198b | ENSMUSG00000027955.13 | -0.629250772 | 1.04E-05 |
| Hist1h2bc | ENSMUSG00000018102.4 | 0.518366657 | 9.64E-06 |
| Ccl28 | ENSMUSG00000074715.2 | 0.695051602 | 9.21E-06 |
| Cbs | ENSMUSG00000024039.11 | -0.731102677 | 8.40E-06 |
| Fancg | ENSMUSG00000028453.7 | 0.578822885 | 8.06E-06 |
| 4930452B06Rik | ENSMUSG00000021747.11 | 0.635208436 | 7.63E-06 |
| Cbr3 | ENSMUSG00000022947.7 | 0.569771491 | 7.63E-06 |
| Npb | ENSMUSG00000044034.8 | 0.754670852 | 7.40E-06 |
| Echdc2 | ENSMUSG00000028601.15 | -0.579606855 | 6.40E-06 |
| Coa4 | ENSMUSG00000044881.6 | -0.572886459 | 6.23E-06 |
| Aass | ENSMUSG00000029695.10 | -0.709445625 | 5.38E-06 |
| Kcnd1 | ENSMUSG00000009731.4 | -0.511103682 | 5.13E-06 |
| Gm17315 | ENSMUSG00000097791.1 | -0.68790215 | 5.11E-06 |
| Cubn | ENSMUSG00000026726.10 | -0.669188353 | 4.99E-06 |
| Ifi30 | ENSMUSG00000031838.7 | -0.533926718 | 4.35E-06 |
| H2-Q2 | ENSMUSG00000091705.5 | 0.754539233 | 3.90E-06 |
| Gm11273 | ENSMUSG00000079941.1 | 0.735022373 | 3.76E-06 |
| Sspn | ENSMUSG00000030255.10 | 0.518189587 | 3.58E-06 |
| Gm14133 | ENSMUSG00000087029.1 | -0.627595046 | 3.57E-06 |
| Arhgap8 | ENSMUSG00000078954.6 | 0.586548095 | 3.52E-06 |
| Esrp1 | ENSMUSG00000040728.12 | -0.741728078 | 3.25E-06 |
| Sla2 | ENSMUSG00000027636.8 | 0.75971919 | 3.22E-06 |
| Glb1l | ENSMUSG00000026200.10 | -0.52057035 | 3.21E-06 |
| Ccdc63 | ENSMUSG00000043036.10 | 0.639125343 | 3.18E-06 |
| Prcp | ENSMUSG00000061119.6 | -0.529616729 | 3.18E-06 |
| Nanos2 | ENSMUSG00000051965.7 | 0.782437197 | 3.07E-06 |
| Moxd1 | ENSMUSG00000020000.7 | 0.530092546 | 2.90E-06 |
| Fbn1 | ENSMUSG00000027204.10 | 0.502610325 | 2.84E-06 |
| Itgam | ENSMUSG00000030786.15 | -0.566123884 | 2.21E-06 |
| 2410127L17Rik | ENSMUSG00000024726.9 | -0.764106889 | 1.85E-06 |
| Gm26583 | ENSMUSG00000097476.1 | -0.794375272 | 1.68E-06 |
| Lair1 | ENSMUSG00000055541.14 | 0.639780951 | 1.67E-06 |
| Ccdc57 | ENSMUSG00000048445.6 | -0.529072505 | 1.53E-06 |
| Nos3 | ENSMUSG00000028978.9 | -0.796499121 | 1.44E-06 |
| Smoc2 | ENSMUSG00000023886.9 | -0.610068581 | 1.44E-06 |
| Cyp4f37 | ENSMUSG00000062464.5 | -0.67106102 | 1.44E-06 |
| Perm1 | ENSMUSG00000078486.3 | 0.804321432 | 1.34E-06 |
| Syt10 | ENSMUSG00000063260.2 | 0.508987026 | 1.30E-06 |
| Slc37a1 | ENSMUSG00000024036.12 | 0.735607972 | 6.62E-07 |
| Gm10263 | ENSMUSG00000066407.3 | 0.823619817 | 6.57E-07 |
| Aldh4a1 | ENSMUSG00000028737.12 | -0.523780759 | 5.67E-07 |

| | | | |
|---|---|---|---|
| 2610305D13Rik | ENSMUSG00000066000.9 | -0.672329997 | 5.44E-07 |
| F5 | ENSMUSG00000026579.8 | -0.766123391 | 4.83E-07 |
| Mlxip | ENSMUSG00000038342.12 | -0.542653171 | 4.56E-07 |
| Folh1 | ENSMUSG00000001773.10 | 0.632504083 | 4.04E-07 |
| Rbm46 | ENSMUSG00000033882.12 | -0.832240761 | 3.81E-07 |
| Mob3b | ENSMUSG00000073910.7 | -0.560039248 | 3.61E-07 |
| Otor | ENSMUSG00000027416.2 | -0.80750483 | 3.58E-07 |
| Rps26-ps1 | ENSMUSG00000059775.4 | -0.708361276 | 2.86E-07 |
| Zfp963 | ENSMUSG00000092260.4 | -0.661727206 | 2.57E-07 |
| Gm14399 | ENSMUSG00000090093.5 | -0.815420581 | 2.26E-07 |
| Gm4980 | ENSMUSG00000096606.2 | 0.52839801 | 2.10E-07 |
| Myo7a | ENSMUSG00000030761.12 | -0.663300448 | 1.78E-07 |
| BC055402 | ENSMUSG00000101429.1 | -0.852827485 | 1.62E-07 |
| Rnaset2b | ENSMUSG00000094724.4 | 0.85925806 | 1.55E-07 |
| Nr6a1 | ENSMUSG00000063972.10 | 0.572026651 | 1.46E-07 |
| Hspa1b | ENSMUSG00000090877.3 | 0.623392612 | 1.44E-07 |
| Cfap46 | ENSMUSG00000049571.13 | -0.58113407 | 1.36E-07 |
| Kcng4 | ENSMUSG00000045246.8 | 0.552550016 | 1.17E-07 |
| Dtd1 | ENSMUSG00000027430.9 | 0.502929076 | 1.13E-07 |
| Ccl21a | ENSMUSG00000094686.1 | 0.729913972 | 9.26E-08 |
| Cadps2 | ENSMUSG00000017978.15 | 0.600107433 | 8.30E-08 |
| Ryr1 | ENSMUSG00000030592.13 | -0.64549873 | 8.12E-08 |
| Gm17546 | ENSMUSG00000078648.2 | -0.730978945 | 6.99E-08 |
| Plekhn1 | ENSMUSG00000078485.2 | -0.64394736 | 5.54E-08 |
| Fibcd1 | ENSMUSG00000026841.7 | 0.639152405 | 5.22E-08 |
| Gm10020 | ENSMUSG00000057262.1 | -0.803408915 | 5.22E-08 |
| Lhx1os | ENSMUSG00000087211.4 | -0.838481212 | 4.42E-08 |
| Tectb | ENSMUSG00000024979.10 | 0.869376405 | 4.22E-08 |
| Lgals4 | ENSMUSG00000053964.13 | -0.803315253 | 4.20E-08 |
| Irak1bp1 | ENSMUSG00000032251.9 | 0.591567773 | 3.92E-08 |
| Nsa2 | ENSMUSG00000060739.7 | -0.586434751 | 3.68E-08 |
| Tmem86b | ENSMUSG00000045282.6 | -0.647063711 | 3.37E-08 |
| Gstp1 | ENSMUSG00000060803.5 | -0.509636682 | 2.59E-08 |
| Cep250 | ENSMUSG00000038241.13 | 0.544480873 | 2.36E-08 |
| Vil1 | ENSMUSG00000026175.9 | 0.901565397 | 2.35E-08 |
| Lipg | ENSMUSG00000053846.4 | -0.539872317 | 2.12E-08 |
| Synj2 | ENSMUSG00000023805.13 | 0.550627223 | 1.45E-08 |
| Enpp2 | ENSMUSG00000022425.12 | 0.644753235 | 1.44E-08 |
| Kif26a | ENSMUSG00000021294.7 | 0.563616612 | 1.35E-08 |
| Ndufb8 | ENSMUSG00000025204.7 | 0.525620867 | 1.17E-08 |
| Apobec2 | ENSMUSG00000040694.3 | 0.556535956 | 1.15E-08 |
| Hist1h2al | ENSMUSG00000091383.1 | -0.86919983 | 1.15E-08 |
| Fcgr1 | ENSMUSG00000015947.7 | -0.857379202 | 1.02E-08 |
| Zkscan1 | ENSMUSG00000029729.9 | -0.524481448 | 8.48E-09 |
| Fras1 | ENSMUSG00000034687.5 | 0.500390861 | 7.06E-09 |

| | | | |
|---|---|---|---|
| H2-DMa | ENSMUSG00000037649.9 | -0.688043181 | 6.73E-09 |
| Vwa5b1 | ENSMUSG00000028753.9 | 0.820764642 | 6.34E-09 |
| Slc16a1 | ENSMUSG00000032902.1 | 0.625083567 | 5.66E-09 |
| Nmnat3 | ENSMUSG00000032456.10 | -0.886217671 | 4.87E-09 |
| Ctss | ENSMUSG00000038642.7 | -0.582298919 | 3.88E-09 |
| Samhd1 | ENSMUSG00000027639.13 | -0.536330446 | 3.35E-09 |
| D630003M21Rik | ENSMUSG00000037813.10 | -0.941527345 | 3.00E-09 |
| Ankrd35 | ENSMUSG00000038354.10 | 0.717295619 | 1.41E-09 |
| Hspe1 | ENSMUSG00000073676.4 | 0.587872204 | 1.39E-09 |
| Gsg1 | ENSMUSG00000030206.10 | 0.553993064 | 1.37E-09 |
| Cpa2 | ENSMUSG00000071553.7 | 0.78639328 | 7.20E-10 |
| Fam111a | ENSMUSG00000024691.10 | -0.845829137 | 6.96E-10 |
| Slc7a11 | ENSMUSG00000027737.7 | -0.781181809 | 6.31E-10 |
| Tmem59l | ENSMUSG00000035964.7 | 0.538314991 | 5.75E-10 |
| Ano6 | ENSMUSG00000064210.6 | -0.558441208 | 5.54E-10 |
| Fxyd6 | ENSMUSG00000066705.6 | 0.666125749 | 4.50E-10 |
| Cdc37l1 | ENSMUSG00000024780.6 | 0.554913032 | 3.78E-10 |
| Gabrr3 | ENSMUSG00000074991.4 | -0.571578245 | 3.78E-10 |
| Ocel1 | ENSMUSG00000002396.8 | 0.846620483 | 3.62E-10 |
| Adamts10 | ENSMUSG00000024299.13 | -0.527822837 | 2.46E-10 |
| Itgb3bp | ENSMUSG00000028549.14 | -0.734683823 | 1.89E-10 |
| P3h4 | ENSMUSG00000006931.12 | -0.639639403 | 1.23E-10 |
| Fbxo43 | ENSMUSG00000048230.5 | 0.985933506 | 1.06E-10 |
| Myo1d | ENSMUSG00000035441.11 | -0.563983532 | 8.91E-11 |
| Bche | ENSMUSG00000027792.8 | 0.727919092 | 8.57E-11 |
| Fam69c | ENSMUSG00000047992.7 | 0.715502695 | 7.69E-11 |
| 9430015G10Rik | ENSMUSG00000059939.10 | -0.557713735 | 7.01E-11 |
| Gnat3 | ENSMUSG00000028777.8 | 0.973565305 | 5.27E-11 |
| A930004D18Rik | ENSMUSG00000054057.2 | 0.59671149 | 4.38E-11 |
| A230087F16Rik | ENSMUSG00000097381.1 | 1.039264351 | 2.88E-11 |
| Ppwd1 | ENSMUSG00000021713.8 | 0.532106469 | 2.82E-11 |
| Itga6 | ENSMUSG00000027111.12 | 0.595201244 | 2.48E-11 |
| Aqp4 | ENSMUSG00000024411.9 | 0.632105433 | 2.05E-11 |
| A530016L24Rik | ENSMUSG00000043122.6 | 0.989749583 | 1.68E-11 |
| H2-K1 | ENSMUSG00000061232.12 | 0.920807862 | 8.63E-12 |
| Upp2 | ENSMUSG00000026839.13 | -0.892776198 | 8.24E-12 |
| Gm26614 | ENSMUSG00000097240.1 | -1.05364663 | 6.54E-12 |
| Ltbp4 | ENSMUSG00000040488.13 | -0.650522532 | 3.94E-12 |
| Zfp955b | ENSMUSG00000096910.1 | -0.601453694 | 3.81E-12 |
| Trpv1 | ENSMUSG00000005952.12 | -1.09462044 | 3.44E-12 |
| Gm8730 | ENSMUSG00000063696.6 | 1.093857891 | 2.91E-12 |
| Gm13157 | ENSMUSG00000078495.7 | -1.036922572 | 2.34E-12 |
| Haus5 | ENSMUSG00000078762.7 | -1.043704145 | 1.84E-12 |
| Ccr6 | ENSMUSG00000040899.10 | -1.00329604 | 1.60E-12 |
| Zcwpw1 | ENSMUSG00000037108.10 | -0.569040104 | 1.32E-12 |

| | | | |
|---|---|---|---|
| Map1lc3a | ENSMUSG00000027602.6 | 0.567596126 | 7.46E-13 |
| Cfap70 | ENSMUSG00000039543.12 | 0.896883427 | 7.20E-13 |
| Gngt2 | ENSMUSG00000038811.10 | -0.999841961 | 7.20E-13 |
| 4930503L19Rik | ENSMUSG00000044906.4 | -0.812095597 | 2.59E-13 |
| Krt24 | ENSMUSG00000020913.3 | -0.932280827 | 1.97E-13 |
| Samd15 | ENSMUSG00000090812.5 | -0.763320161 | 1.17E-13 |
| Rab27b | ENSMUSG00000024511.12 | 0.595012147 | 1.04E-13 |
| Fbxo44 | ENSMUSG00000029001.12 | 0.611905657 | 7.68E-14 |
| Ring1 | ENSMUSG00000024325.8 | -0.676803013 | 7.62E-14 |
| Igj | ENSMUSG00000067149.6 | 0.690412896 | 6.94E-14 |
| Rfxank | ENSMUSG00000036120.9 | 0.575214029 | 5.28E-14 |
| Vcam1 | ENSMUSG00000027962.11 | 0.787927077 | 5.20E-14 |
| Gm8909 | ENSMUSG00000073402.8 | 1.169875406 | 3.44E-14 |
| Hexdc | ENSMUSG00000039307.13 | -0.614404354 | 1.58E-14 |
| Nudt19 | ENSMUSG00000034875.5 | 0.741253607 | 1.36E-14 |
| Cfap126 | ENSMUSG00000026649.11 | 1.155168057 | 1.27E-14 |
| Tuba4a | ENSMUSG00000026202.10 | 0.819390438 | 1.23E-14 |
| Dxo | ENSMUSG00000040482.12 | -0.576322828 | 8.40E-15 |
| Adamts7 | ENSMUSG00000032363.12 | -0.897427501 | 6.58E-15 |
| Zfp729a | ENSMUSG00000021510.10 | -0.864569769 | 4.93E-15 |
| AI467606 | ENSMUSG00000045165.5 | 1.159507773 | 2.79E-15 |
| Bpifb6 | ENSMUSG00000068009.8 | -1.058688047 | 2.20E-15 |
| Ccna2 | ENSMUSG00000027715.6 | 0.972399884 | 1.38E-15 |
| Frzb | ENSMUSG00000027004.3 | 0.621250918 | 4.41E-16 |
| Cyp4f16 | ENSMUSG00000048440.12 | 0.943522257 | 2.52E-16 |
| Rps4l | ENSMUSG00000063171.3 | -0.884004126 | 1.62E-16 |
| Dusp4 | ENSMUSG00000031530.6 | 0.625665551 | 1.12E-16 |
| Trim68 | ENSMUSG00000073968.3 | -0.968771797 | 9.27E-17 |
| Pdk1 | ENSMUSG00000006494.8 | 0.525088303 | 9.20E-17 |
| Sh2d2a | ENSMUSG00000028071.9 | -1.202344098 | 7.31E-17 |
| Hist2h2bb | ENSMUSG00000050936.5 | -1.290682591 | 5.69E-17 |
| Pus7 | ENSMUSG00000057541.11 | -0.680157052 | 5.03E-17 |
| Stx3 | ENSMUSG00000041488.12 | -0.621444848 | 5.03E-17 |
| Alox5ap | ENSMUSG00000060063.6 | -1.26184423 | 3.28E-17 |
| Pgap2 | ENSMUSG00000030990.14 | -0.941898609 | 2.78E-17 |
| 4921501E09Rik | ENSMUSG00000023350.4 | -1.235009884 | 1.79E-17 |
| St7 | ENSMUSG00000029534.14 | -0.749963871 | 6.75E-18 |
| Zfp955a | ENSMUSG00000094441.1 | 0.733697406 | 6.33E-18 |
| Stk19 | ENSMUSG00000061207.8 | -0.816553797 | 3.61E-18 |
| Ulk4 | ENSMUSG00000040936.11 | -0.822923588 | 2.74E-18 |
| Gm21887 | ENSMUSG00000095562.4 | 1.341637907 | 1.37E-18 |
| Frem2 | ENSMUSG00000037016.8 | 0.899741099 | 6.28E-19 |
| Gm12866 | ENSMUSG00000066060.5 | -1.30025955 | 5.86E-19 |
| Slc35e4 | ENSMUSG00000048807.2 | 0.950351019 | 2.14E-19 |
| Anxa1 | ENSMUSG00000024659.11 | -1.320170542 | 2.00E-19 |

| | | | |
|---|---|---|---|
| Fndc1 | ENSMUSG00000071984.7 | -0.924101471 | 1.47E-19 |
| Gem | ENSMUSG00000028214.10 | -0.917873857 | 1.21E-19 |
| Ccl27a | ENSMUSG00000073888.9 | 1.081868704 | 5.91E-20 |
| Rgs22 | ENSMUSG00000037627.12 | 0.735004069 | 3.69E-20 |
| Rreb1 | ENSMUSG00000039087.13 | -0.718518234 | 1.93E-20 |
| Ccdc171 | ENSMUSG00000052407.13 | -1.118809169 | 1.83E-20 |
| Mt3 | ENSMUSG00000031760.8 | 1.033404476 | 3.23E-22 |
| Clstn2 | ENSMUSG00000032452.9 | 0.71564947 | 2.37E-22 |
| Krt18 | ENSMUSG00000023043.6 | 1.062922846 | 1.25E-22 |
| Ckb | ENSMUSG00000001270.8 | 0.598642551 | 9.61E-23 |
| Zfyve28 | ENSMUSG00000037224.12 | -0.974003967 | 3.81E-23 |
| Fcrls | ENSMUSG00000015852.10 | -1.29889118 | 2.97E-23 |
| BC022687 | ENSMUSG00000037594.7 | 0.764367756 | 3.65E-24 |
| Bfsp2 | ENSMUSG00000032556.10 | 1.541379993 | 1.54E-24 |
| Draxin | ENSMUSG00000029005.4 | -0.892353752 | 8.47E-25 |
| Trim12c | ENSMUSG00000057143.11 | -1.440058456 | 6.77E-26 |
| Stag3 | ENSMUSG00000036928.11 | -1.1329751 | 3.56E-26 |
| A930001A20Rik | ENSMUSG00000098008.1 | 1.554374355 | 2.08E-26 |
| Malt1 | ENSMUSG00000032688.7 | -0.712375007 | 1.17E-26 |
| Tmem144 | ENSMUSG00000027956.8 | -1.241818682 | 1.62E-27 |
| H2-Q4 | ENSMUSG00000035929.11 | 1.275125296 | 5.65E-28 |
| Gm7120 | ENSMUSG00000074634.8 | 1.369980756 | 9.01E-29 |
| Vcp | ENSMUSG00000028452.7 | 0.876030174 | 3.18E-29 |
| Zfp738 | ENSMUSG00000048280.14 | 0.853811659 | 6.58E-30 |
| 4930447C04Rik | ENSMUSG00000021098.11 | -1.082528793 | 1.49E-30 |
| Duxbl1 | ENSMUSG00000048502.11 | -1.692593535 | 1.30E-30 |
| Apcdd1 | ENSMUSG00000071847.9 | -0.837383081 | 6.75E-31 |
| Ceacam10 | ENSMUSG00000054169.6 | -1.644244678 | 6.75E-31 |
| Gm21967 | ENSMUSG00000094114.1 | 1.137401022 | 2.61E-31 |
| Trp53cor1 | ENSMUSG00000085912.1 | -1.666927018 | 2.31E-33 |
| Tmem169 | ENSMUSG00000026188.8 | 0.72417622 | 6.04E-34 |
| Aurka | ENSMUSG00000027496.12 | -1.411782715 | 2.31E-35 |
| Lrpprc | ENSMUSG00000024120.9 | 0.777528424 | 2.77E-38 |
| Il11ra1 | ENSMUSG00000073889.7 | 1.323589324 | 1.59E-40 |
| Fam210b | ENSMUSG00000027495.4 | -0.913504807 | 1.15E-40 |
| Gm9847 | ENSMUSG00000050974.3 | 1.995253564 | 4.67E-41 |
| Syt15 | ENSMUSG00000041479.10 | 1.409765644 | 5.88E-47 |
| Polr2k | ENSMUSG00000045996.9 | -1.610521968 | 1.71E-49 |
| Cyp4f13 | ENSMUSG00000024055.11 | -1.551764604 | 8.76E-50 |
| Olfml3 | ENSMUSG00000027848.12 | -1.766066708 | 3.43E-53 |
| Tagap | ENSMUSG00000033450.7 | -2.230848477 | 3.43E-53 |
| Ush1c | ENSMUSG00000030838.14 | -1.908074887 | 4.09E-55 |
| Gm1840 | ENSMUSG00000043192.3 | 1.573031178 | 1.81E-59 |
| Tcte2 | ENSMUSG00000038347.11 | 1.678681332 | 2.26E-69 |
| Plcd4 | ENSMUSG00000026173.12 | 1.743047481 | 1.15E-76 |

| | | | |
|---|---|---|---|
| H2-D1 | ENSMUSG00000073411.8 | 2.388354934 | 1.33E-186 |
| Tpm3-rs7 | ENSMUSG00000058126.6 | -3.570676155 | 2.29E-236 |

**Significantly differentially expressed genes (P14 Apcdd1-/- vs Wt retina)**

| Gene name | Ensembl_ID | log2FoldChange | padj |
|---|---|---|---|
| NA | ENSMUSG00000086344 | 8.108683749 | 1.66E-12 |
| Gm9847 | ENSMUSG00000050974 | 7.990955642 | 1.63E-12 |
| Gm18562 | ENSMUSG00000110123 | 7.58618079 | 4.37E-11 |
| Gm11342 | ENSMUSG00000084788 | 7.540029838 | 7.98E-17 |
| Arhgap8 | ENSMUSG00000078954 | 7.382382014 | 7.41E-78 |
| Rps3a3 | ENSMUSG00000059751 | 7.352887321 | 6.01E-201 |
| Gm14014 | ENSMUSG00000087225 | 6.910329915 | 3.34E-09 |
| Gm10874 | ENSMUSG00000075591 | 6.341061371 | 2.81E-69 |
| Gm20775 | ENSMUSG00000075359 | 6.295301189 | 5.86E-08 |
| Gm14719 | ENSMUSG00000082192 | 6.188593798 | 1.23E-44 |
| Tgif2-ps2 | ENSMUSG00000045842 | 6.066846631 | 2.92E-07 |
| A830036E02I | ENSMUSG00000084890 | 5.955163028 | 2.14E-09 |
| Gm6166 | ENSMUSG00000074280 | 5.401659686 | 5.67E-20 |
| Gm15772 | ENSMUSG00000062353 | 5.06680896 | 6.02E-30 |
| Gm42764 | ENSMUSG00000106173 | 4.849741248 | 3.37E-12 |
| Gm20305 | ENSMUSG00000103979 | 4.665814571 | 9.16E-16 |
| Slc17a1 | ENSMUSG00000021335 | 4.580235931 | 3.88E-14 |
| Gm10654 | ENSMUSG00000074252 | 4.388337902 | 8.00E-11 |
| Gm5436 | ENSMUSG00000042962 | 4.288417897 | 8.00E-88 |
| Rps3a2 | ENSMUSG00000062611 | 4.100287793 | 7.03E-44 |
| Gm8116 | ENSMUSG00000059422 | 4.084389723 | 2.59E-10 |
| Rnaset2b | ENSMUSG00000094724 | 4.066275029 | 8.84E-74 |
| Gm8730 | ENSMUSG00000063696 | 3.926164214 | 1.22E-14 |
| A230087F16I | ENSMUSG00000097381 | 3.856282894 | 1.02E-07 |
| Slc17a3 | ENSMUSG00000036083 | 3.82965835 | 4.03E-16 |
| Cd1d2 | ENSMUSG00000041750 | 3.690347587 | 3.35E-11 |
| Lbp | ENSMUSG00000016024 | 3.66432372 | 9.50E-86 |
| Cwc22 | ENSMUSG00000027014 | 3.609834916 | 4.13E-24 |
| Aspg | ENSMUSG00000037686 | 3.581579065 | 2.38E-08 |
| Gm4134 | ENSMUSG00000092571 | 3.556426917 | 8.40E-28 |
| 1500004A13I | ENSMUSG00000098912 | 3.497544379 | 5.21E-16 |
| Nov | ENSMUSG00000037362 | 3.312603856 | 2.75E-10 |
| Gm11273 | ENSMUSG00000079941 | 3.294379441 | 8.49E-68 |
| A530016L24I | ENSMUSG00000043122 | 3.223445694 | 8.51E-13 |
| Gm6768 | ENSMUSG00000021908 | 3.103662176 | 3.17E-17 |
| Cbr2 | ENSMUSG00000025150 | 3.101106367 | 4.49E-06 |
| Gm6877 | ENSMUSG00000083557 | 3.02337037 | 1.49E-06 |
| Gm13577 | ENSMUSG00000082127 | 3.017930391 | 1.98E-08 |
| Got2-ps1 | ENSMUSG00000080935 | 2.994476611 | 6.12E-31 |
| Ttc30a2 | ENSMUSG00000075272 | 2.975271113 | 5.53E-29 |
| B3gnt3 | ENSMUSG00000031803 | 2.852536932 | 2.49E-10 |
| Foxe3 | ENSMUSG00000044518 | 2.792321784 | 0.00933981 |

| | | | |
|---|---|---|---|
| Gm14026 | ENSMUSG00000082274 | 2.759021312 | 6.61E-23 |
| Oscar | ENSMUSG00000054594 | 2.693880227 | 1.05E-05 |
| Gm5801 | ENSMUSG00000058581 | 2.688879965 | 5.09E-08 |
| Gm14165 | ENSMUSG00000081434 | 2.679349452 | 2.17E-06 |
| Gm9816 | ENSMUSG00000085783 | 2.643101239 | 5.04E-27 |
| 4930458D05I | ENSMUSG00000087611 | 2.623931069 | 4.62E-17 |
| Rab11b-ps2 | ENSMUSG00000095690 | 2.525983839 | 7.72E-30 |
| Mrc1 | ENSMUSG00000026712 | 2.420444055 | 2.52E-09 |
| Spp1 | ENSMUSG00000029304 | 2.415034388 | 0.00525567 |
| Dydc2 | ENSMUSG00000021791 | 2.410831382 | 2.43E-06 |
| 4930481B07I | ENSMUSG00000085956 | 2.38558657 | 8.45E-41 |
| Rps2-ps13 | ENSMUSG00000081684 | 2.36071515 | 0.005733 |
| Wnt10a | ENSMUSG00000026167 | 2.360052333 | 3.88E-05 |
| Gja8 | ENSMUSG00000049908 | 2.342451849 | 5.08E-05 |
| Kcnk15 | ENSMUSG00000035238 | 2.304550952 | 9.82E-11 |
| Glycam1 | ENSMUSG00000022491 | 2.165587869 | 0.00376477 |
| AI506816 | ENSMUSG00000105987 | 2.156008286 | 1.76E-18 |
| A930001A20 | ENSMUSG00000098008 | 2.14032304 | 2.70E-11 |
| Arsi | ENSMUSG00000036412 | 2.135457449 | 4.90E-31 |
| Gm3608 | ENSMUSG00000067017 | 2.124032948 | 1.15E-24 |
| Slc25a41 | ENSMUSG00000011486 | 2.113241363 | 2.22E-09 |
| R3hdml | ENSMUSG00000078949 | 2.099107796 | 2.43E-05 |
| Tcte2 | ENSMUSG00000038347 | 2.097197284 | 4.76E-54 |
| H2-D1 | ENSMUSG00000073411 | 2.090209147 | 1.85E-98 |
| Slc22a6 | ENSMUSG00000024650 | 2.084040114 | 0.0004524 |
| Tspan11 | ENSMUSG00000030351 | 2.06253516 | 1.44E-12 |
| Colec11 | ENSMUSG00000036655 | 2.031282409 | 0.00119876 |
| Gm266 | ENSMUSG00000010529 | 1.949282941 | 0.0009726 |
| Epo | ENSMUSG00000029711 | 1.940260699 | 0.00071721 |
| Insl6 | ENSMUSG00000050957 | 1.936304803 | 0.00240117 |
| Srd5a2 | ENSMUSG00000038541 | 1.933484384 | 4.52E-05 |
| Gm4349 | ENSMUSG00000097532 | 1.891216928 | 0.00094119 |
| Hs3st3a1 | ENSMUSG00000047759 | 1.890463021 | 8.29E-18 |
| D430036J16I | ENSMUSG00000097466 | 1.861112937 | 3.59E-07 |
| Tbx22 | ENSMUSG00000031241 | 1.847598504 | 0.00189848 |
| Cdh1 | ENSMUSG00000000303 | 1.846869256 | 0.0002214 |
| Igfals | ENSMUSG00000046070 | 1.824903006 | 0.00046619 |
| Gm13778 | ENSMUSG00000086495 | 1.822342499 | 2.70E-05 |
| 2610528A11I | ENSMUSG00000096001 | 1.81643662 | 9.66E-06 |
| Gm12758 | ENSMUSG00000085105 | 1.809558805 | 3.44E-26 |
| Tinagl1 | ENSMUSG00000028776 | 1.794251824 | 1.48E-10 |
| Gstm2 | ENSMUSG00000040562 | 1.789107874 | 0.00303107 |
| Nos3 | ENSMUSG00000028978 | 1.787594364 | 2.17E-06 |
| Pamr1 | ENSMUSG00000027188 | 1.751883706 | 9.44E-09 |
| Trim5 | ENSMUSG00000060441 | 1.729971267 | 2.69E-06 |

| | | | |
|---|---|---|---|
| Gm15482 | ENSMUSG00000086711 | 1.727767562 | 1.16E-06 |
| Bpifb5 | ENSMUSG00000038572 | 1.725865307 | 0.00122265 |
| Fosb | ENSMUSG00000003545 | 1.70050788 | 0.02595726 |
| Cfi | ENSMUSG00000058952 | 1.696312259 | 0.0003064 |
| Myl2 | ENSMUSG00000013936 | 1.690203416 | 0.00246464 |
| Il11ra1 | ENSMUSG00000073889 | 1.670649256 | 8.13E-52 |
| E030042O20 | ENSMUSG00000087222 | 1.644744395 | 1.53E-08 |
| Mfap4 | ENSMUSG00000042436 | 1.624074261 | 0.00057869 |
| Ms4a6d | ENSMUSG00000024679 | 1.623513616 | 0.00325965 |
| Masp2 | ENSMUSG00000028979 | 1.606786781 | 0.005733 |
| Vil1 | ENSMUSG00000026175 | 1.598407979 | 5.61E-12 |
| Ctla2b | ENSMUSG00000074874 | 1.593284887 | 0.00294843 |
| Klf1 | ENSMUSG00000054191 | 1.588669015 | 0.00018716 |
| Ltbp2 | ENSMUSG00000002020 | 1.588659515 | 0.0001187 |
| Rspo1 | ENSMUSG00000028871 | 1.582372761 | 0.00074649 |
| Slc38a8 | ENSMUSG00000034224 | 1.581811898 | 2.70E-06 |
| 4930481A15I | ENSMUSG00000086938 | 1.572222025 | 0.00025393 |
| F13a1 | ENSMUSG00000039109 | 1.568425028 | 8.19E-05 |
| H60b | ENSMUSG00000075297 | 1.564967446 | 0.00124559 |
| Gm24265 | ENSMUSG00000096243 | 1.558007124 | 0.01861728 |
| H2-Q4 | ENSMUSG00000035929 | 1.552829397 | 4.48E-42 |
| Npb | ENSMUSG00000044034 | 1.551665791 | 3.43E-07 |
| Gm37986 | ENSMUSG00000104212 | 1.545348399 | 0.0043398 |
| Best2 | ENSMUSG00000052819 | 1.541727726 | 0.00457197 |
| Gm4742 | ENSMUSG00000105461 | 1.54090925 | 0.00377074 |
| Galnt12 | ENSMUSG00000039774 | 1.535217752 | 5.40E-05 |
| Gm26587 | ENSMUSG00000097470 | 1.534229259 | 1.23E-15 |
| 1810010K12I | ENSMUSG00000056329 | 1.534024623 | 3.29E-24 |
| Gm43718 | ENSMUSG00000104905 | 1.533808809 | 0.01250795 |
| Serpinb6b | ENSMUSG00000042842 | 1.528406514 | 4.50E-07 |
| Ccdc175 | ENSMUSG00000021086 | 1.517951017 | 0.00035461 |
| Adh1 | ENSMUSG00000074207 | 1.515519746 | 0.00396103 |
| Perp | ENSMUSG00000019851 | 1.515513357 | 1.85E-26 |
| Dbh | ENSMUSG00000000889 | 1.506612744 | 0.00957474 |
| Gm21887 | ENSMUSG00000095562 | 1.50022612 | 1.92E-13 |
| Slc26a11 | ENSMUSG00000039908 | 1.498214654 | 1.34E-43 |
| AI467606 | ENSMUSG00000045165 | 1.496868568 | 3.21E-10 |
| Kcne1 | ENSMUSG00000039639 | 1.494998998 | 0.00335182 |
| Dio3 | ENSMUSG00000075707 | 1.48746969 | 2.61E-09 |
| Tfpi | ENSMUSG00000027082 | 1.484475813 | 9.95E-14 |
| Agtr1a | ENSMUSG00000049115 | 1.475187374 | 0.00443047 |
| Gm31805 | ENSMUSG00000110661 | 1.454195225 | 0.00027134 |
| Optc | ENSMUSG00000010311 | 1.451802056 | 0.00459598 |
| Ribc2 | ENSMUSG00000022431 | 1.450659173 | 0.04209493 |
| Gm45246 | ENSMUSG00000109645 | 1.448005459 | 0.00907163 |

| | | | |
|---|---|---|---|
| 4930405A21I | ENSMUSG00000086638 | 1.444489465 | 3.04E-27 |
| Rmrp | ENSMUSG00000088088 | 1.439868881 | 0.0005277 |
| Cyp2c39 | ENSMUSG00000025003 | 1.439439876 | 0.01016389 |
| Crhbp | ENSMUSG00000021680 | 1.436138821 | 2.33E-06 |
| 5830417I10R | ENSMUSG00000078684 | 1.435522429 | 1.68E-22 |
| Gm13453 | ENSMUSG00000083240 | 1.434421187 | 3.63E-20 |
| Adgrg6 | ENSMUSG00000039116 | 1.432002884 | 2.20E-11 |
| Gm45844 | ENSMUSG00000110105 | 1.431699835 | 2.34E-07 |
| Myom2 | ENSMUSG00000031461 | 1.425060051 | 0.00444023 |
| Gm7879 | ENSMUSG00000033036 | 1.414235518 | 0.00218171 |
| Abi3bp | ENSMUSG00000035258 | 1.4084498 | 2.72E-06 |
| Fbln1 | ENSMUSG00000006369 | 1.382214811 | 0.00788349 |
| Pitx3 | ENSMUSG00000025229 | 1.381978192 | 0.0062771 |
| Hsd17b2 | ENSMUSG00000031844 | 1.380397299 | 6.13E-10 |
| Syt15 | ENSMUSG00000041479 | 1.374086256 | 2.57E-30 |
| Apoc1 | ENSMUSG00000040564 | 1.37036324 | 2.95E-07 |
| Gm43031 | ENSMUSG00000106535 | 1.370127609 | 1.98E-08 |
| Lyz2 | ENSMUSG00000069516 | 1.369320898 | 0.00720403 |
| Gm4353 | ENSMUSG00000091900 | 1.36114119 | 6.64E-05 |
| Gm10320 | ENSMUSG00000092116 | 1.358390397 | 2.07E-15 |
| Pla2g4a | ENSMUSG00000056220 | 1.334802516 | 1.81E-07 |
| Klhdc7a | ENSMUSG00000078234 | 1.330178357 | 3.91E-06 |
| Orm2 | ENSMUSG00000061540 | 1.329458148 | 1.43E-06 |
| Gm9967 | ENSMUSG00000055323 | 1.322407183 | 9.20E-13 |
| Gm43622 | ENSMUSG00000104988 | 1.313669339 | 0.00030451 |
| Fos | ENSMUSG00000021250 | 1.30794222 | 0.0354153 |
| Fmod | ENSMUSG00000041559 | 1.306987968 | 0.00155253 |
| Mme | ENSMUSG00000027820 | 1.295479039 | 0.00517758 |
| Adgre1 | ENSMUSG00000004730 | 1.291605459 | 0.01488638 |
| Slc38a4 | ENSMUSG00000022464 | 1.290902481 | 4.75E-05 |
| Phex | ENSMUSG00000057457 | 1.288830124 | 0.03350989 |
| S100a6 | ENSMUSG00000001025 | 1.288018029 | 7.85E-12 |
| Tmem267 | ENSMUSG00000074634 | 1.28684665 | 7.00E-30 |
| Col4a6 | ENSMUSG00000031273 | 1.283940675 | 0.00198635 |
| Frem2 | ENSMUSG00000037016 | 1.279804121 | 1.59E-07 |
| Gfap | ENSMUSG00000020932 | 1.276516329 | 0.01551296 |
| Fbln2 | ENSMUSG00000064080 | 1.269181266 | 4.85E-09 |
| C430049B03I | ENSMUSG00000087365 | 1.265729531 | 0.02173653 |
| Esyt3 | ENSMUSG00000037681 | 1.263137435 | 0.00368496 |
| 1500015O10 | ENSMUSG00000026051 | 1.258426491 | 0.00092532 |
| Gfra3 | ENSMUSG00000024366 | 1.257932278 | 2.95E-05 |
| Stk26 | ENSMUSG00000031112 | 1.257858655 | 0.00276612 |
| Ndc80 | ENSMUSG00000024056 | 1.257749786 | 0.03031214 |
| Mia | ENSMUSG00000089661 | 1.25594918 | 0.01810423 |
| Bcl3 | ENSMUSG00000053175 | 1.249762073 | 0.00302196 |

| | | | |
|---|---|---|---|
| NA | ENSMUSG00000094114 | 1.247984495 | 1.24E-34 |
| Kifc1 | ENSMUSG00000079553 | 1.247358144 | 4.48E-14 |
| Btnl9 | ENSMUSG00000040283 | 1.242633882 | 0.00020549 |
| Ptafr | ENSMUSG00000056529 | 1.239189115 | 0.01105472 |
| Zic4 | ENSMUSG00000036972 | 1.235011777 | 4.36E-05 |
| Wnt7b | ENSMUSG00000022382 | 1.232728086 | 3.46E-07 |
| Gm35040 | ENSMUSG00000108616 | 1.231033678 | 0.01958516 |
| Ifitm1 | ENSMUSG00000025491 | 1.229138913 | 0.00504677 |
| Slit3 | ENSMUSG00000056427 | 1.226470085 | 0.00217061 |
| Vcan | ENSMUSG00000021614 | 1.222609123 | 0.00673527 |
| Serping1 | ENSMUSG00000023224 | 1.220599464 | 4.85E-09 |
| Gm26804 | ENSMUSG00000097263 | 1.219630795 | 0.03713708 |
| Gstm1 | ENSMUSG00000058135 | 1.218773352 | 5.65E-09 |
| Gm3555 | ENSMUSG00000089791 | 1.218120356 | 0.00836628 |
| Rab38 | ENSMUSG00000030559 | 1.212798165 | 9.02E-08 |
| Gm16306 | ENSMUSG00000089780 | 1.210269839 | 0.0497447 |
| Stac | ENSMUSG00000032502 | 1.208317116 | 3.35E-07 |
| Slc13a4 | ENSMUSG00000029843 | 1.207039433 | 0.03239385 |
| Ism2 | ENSMUSG00000050671 | 1.201608452 | 5.67E-05 |
| Notum | ENSMUSG00000042988 | 1.199079514 | 3.84E-07 |
| Gja5 | ENSMUSG00000057123 | 1.189320041 | 0.0002979 |
| Hspb2 | ENSMUSG00000038086 | 1.187442123 | 0.00334527 |
| 2310043M15 | ENSMUSG00000097768 | 1.186969156 | 0.03758076 |
| Cyp11a1 | ENSMUSG00000032323 | 1.181360576 | 0.02203963 |
| Gm12355 | ENSMUSG00000078134 | 1.177217487 | 0.00838472 |
| Colec12 | ENSMUSG00000036103 | 1.176767216 | 7.62E-08 |
| Klk8 | ENSMUSG00000064023 | 1.172902887 | 0.00089172 |
| Ccdc122 | ENSMUSG00000034795 | 1.171298628 | 0.00879772 |
| Lrpprc | ENSMUSG00000024120 | 1.16849876 | 3.48E-65 |
| A730049H05 | ENSMUSG00000048636 | 1.168396939 | 0.00831325 |
| Gkn3 | ENSMUSG00000030048 | 1.166167435 | 0.02898672 |
| Rpl14-ps1 | ENSMUSG00000046721 | 1.166030624 | 7.38E-09 |
| Col6a3 | ENSMUSG00000048126 | 1.165729253 | 0.00155246 |
| Gnat3 | ENSMUSG00000028777 | 1.16433841 | 0.00033438 |
| Mpzl2 | ENSMUSG00000032092 | 1.163440989 | 0.04773487 |
| 1010001B22 | ENSMUSG00000097863 | 1.161817764 | 0.02280054 |
| Cpz | ENSMUSG00000036596 | 1.161286685 | 8.27E-05 |
| Pbk | ENSMUSG00000022033 | 1.160164893 | 0.01447432 |
| Tnfaip8l3 | ENSMUSG00000074345 | 1.160156429 | 0.00055663 |
| Ltc4s | ENSMUSG00000020377 | 1.155019326 | 0.00703393 |
| Msx1 | ENSMUSG00000048450 | 1.151656255 | 7.62E-05 |
| Pzp | ENSMUSG00000030359 | 1.144112268 | 0.01934523 |
| Fbxl7 | ENSMUSG00000043556 | 1.138323426 | 0.00269157 |
| Tcim | ENSMUSG00000056313 | 1.1358336 | 4.30E-13 |
| Aldh3a1 | ENSMUSG00000019102 | 1.135168077 | 0.0109155 |

| | | | |
|---|---|---|---|
| Cd86 | ENSMUSG00000022901 | 1.129455789 | 0.0193409 |
| A330040F15I | ENSMUSG00000086213 | 1.129377978 | 2.60E-10 |
| Gpr137b | ENSMUSG00000021306 | 1.128420356 | 2.64E-24 |
| Gm32736 | ENSMUSG00000106483 | 1.126698669 | 0.0317984 |
| Serpine1 | ENSMUSG00000037411 | 1.124201947 | 1.73E-07 |
| H2-Q2 | ENSMUSG00000091705 | 1.123528441 | 1.87E-06 |
| Gm29593 | ENSMUSG00000099440 | 1.123241777 | 0.00608742 |
| Spink13 | ENSMUSG00000073551 | 1.120839523 | 0.00050239 |
| Gm43766 | ENSMUSG00000105868 | 1.119508672 | 0.02587904 |
| Myh3 | ENSMUSG00000020908 | 1.119324177 | 0.00016378 |
| Sox1ot | ENSMUSG00000047935 | 1.119293856 | 0.01018901 |
| Gm11413 | ENSMUSG00000086321 | 1.117339398 | 0.03698411 |
| Folr1 | ENSMUSG00000001827 | 1.113616067 | 0.00043138 |
| Gm13212 | ENSMUSG00000078502 | 1.112132223 | 0.01870552 |
| Plau | ENSMUSG00000021822 | 1.112059483 | 0.00834209 |
| Mindy4b-ps | ENSMUSG00000101860 | 1.111850305 | 1.53E-08 |
| Islr | ENSMUSG00000037206 | 1.111026998 | 0.00801283 |
| Egfl8 | ENSMUSG00000015467 | 1.110587307 | 7.79E-09 |
| Cd274 | ENSMUSG00000016496 | 1.110569221 | 0.00267594 |
| Cd300a | ENSMUSG00000034652 | 1.110200568 | 0.00403279 |
| Scd4 | ENSMUSG00000050195 | 1.106568611 | 0.03517101 |
| Gm26880 | ENSMUSG00000096979 | 1.105990007 | 0.01052169 |
| Slc16a12 | ENSMUSG00000009378 | 1.100564726 | 3.74E-05 |
| 6030498E09I | ENSMUSG00000051361 | 1.100450449 | 0.04821641 |
| Gm45407 | ENSMUSG00000109873 | 1.099616109 | 0.00472962 |
| Atad3aos | ENSMUSG00000054514 | 1.095477214 | 0.0001068 |
| Cpsf4l | ENSMUSG00000018727 | 1.094196758 | 0.00210323 |
| Lrp8os2 | ENSMUSG00000073779 | 1.091045187 | 2.85E-07 |
| Gm45285 | ENSMUSG00000109732 | 1.089013305 | 0.04937629 |
| Aif1 | ENSMUSG00000024397 | 1.087313638 | 0.00035311 |
| Ccna2 | ENSMUSG00000027715 | 1.087289009 | 3.26E-12 |
| Ccl28 | ENSMUSG00000074715 | 1.086391655 | 1.08E-06 |
| Hao2 | ENSMUSG00000027870 | 1.085158784 | 0.00010033 |
| Penk | ENSMUSG00000045573 | 1.083956414 | 2.09E-32 |
| H2-K1 | ENSMUSG00000061232 | 1.081186286 | 1.32E-18 |
| Gm9625 | ENSMUSG00000097906 | 1.080343236 | 0.01482196 |
| Gm32585 | ENSMUSG00000105039 | 1.07481856 | 3.90E-05 |
| 1-Mar | ENSMUSG00000026621 | 1.074507753 | 0.00154706 |
| Serpinf1 | ENSMUSG00000000753 | 1.067388633 | 1.65E-10 |
| Eif4ebp1 | ENSMUSG00000031490 | 1.066726141 | 3.37E-08 |
| Dtl | ENSMUSG00000037474 | 1.066053113 | 0.01548692 |
| Wnt16 | ENSMUSG00000029671 | 1.064196984 | 0.00565366 |
| Rps19-ps6 | ENSMUSG00000096942 | 1.059951505 | 1.62E-06 |
| Nudt19 | ENSMUSG00000034875 | 1.059135142 | 3.77E-43 |
| Hrct1 | ENSMUSG00000071001 | 1.057520905 | 0.00186559 |

| | | | |
|---|---|---|---|
| Cldn19 | ENSMUSG00000066058 | 1.053228165 | 9.32E-05 |
| Gm5874 | ENSMUSG00000071568 | 1.049592617 | 0.01170605 |
| Pglyrp1 | ENSMUSG00000030413 | 1.049464701 | 7.00E-06 |
| Ogn | ENSMUSG00000021390 | 1.047494775 | 2.03E-05 |
| Mt3 | ENSMUSG00000031760 | 1.044478444 | 2.70E-17 |
| Wfdc1 | ENSMUSG00000023336 | 1.042895268 | 5.75E-09 |
| Tnxb | ENSMUSG00000033327 | 1.042242071 | 2.26E-05 |
| Pcolce | ENSMUSG00000029718 | 1.038902907 | 5.17E-08 |
| Cybrd1 | ENSMUSG00000027015 | 1.036211782 | 0.00377074 |
| Tmprss11e | ENSMUSG00000054537 | 1.035805779 | 0.00296115 |
| Tnni1 | ENSMUSG00000026418 | 1.035181733 | 0.00139115 |
| Hist1h2bg | ENSMUSG00000058385 | 1.034574331 | 0.021369 |
| Fzd1 | ENSMUSG00000044674 | 1.033822772 | 4.64E-14 |
| Rpph1 | ENSMUSG00000092837 | 1.02539214 | 0.00011072 |
| Lama2 | ENSMUSG00000019899 | 1.024020241 | 1.41E-05 |
| Blnk | ENSMUSG00000061132 | 1.023036479 | 0.00130357 |
| Otx1 | ENSMUSG00000005917 | 1.02285007 | 0.00054814 |
| Btk | ENSMUSG00000031264 | 1.022074131 | 0.03035691 |
| Zfp185 | ENSMUSG00000031351 | 1.02155113 | 2.71E-05 |
| Lix1 | ENSMUSG00000047786 | 1.021479733 | 9.74E-09 |
| Gm5841 | ENSMUSG00000103921 | 1.020322444 | 0.04937629 |
| Col10a1 | ENSMUSG00000039462 | 1.019678078 | 0.00447051 |
| Htra3 | ENSMUSG00000029096 | 1.018546679 | 3.55E-08 |
| Trpm5 | ENSMUSG00000009246 | 1.016581043 | 0.00284813 |
| S100a4 | ENSMUSG00000001020 | 1.016141116 | 0.00157684 |
| D630039A03 | ENSMUSG00000052117 | 1.014544535 | 0.00373486 |
| Uckl1os | ENSMUSG00000010492 | 1.012004955 | 6.89E-09 |
| Wdr86 | ENSMUSG00000055235 | 1.008316018 | 0.00101625 |
| Atp4a | ENSMUSG00000005553 | 1.004164941 | 0.00535631 |
| Zfp738 | ENSMUSG00000048280 | 1.001088578 | 1.64E-19 |
| Gm13068 | ENSMUSG00000085403 | 0.997406642 | 4.87E-28 |
| Lair1 | ENSMUSG00000055541 | 0.997093315 | 2.04E-07 |
| Kctd4 | ENSMUSG00000046523 | 0.993144452 | 5.66E-05 |
| Hspg2 | ENSMUSG00000028763 | 0.991144861 | 0.02626044 |
| Car3 | ENSMUSG00000027559 | 0.98942727 | 7.68E-07 |
| Apbb1ip | ENSMUSG00000026786 | 0.988410881 | 0.03571352 |
| Fcgr2b | ENSMUSG00000026656 | 0.986262353 | 0.03730561 |
| Cntf | ENSMUSG00000079415 | 0.983599894 | 0.00443428 |
| Gm17322 | ENSMUSG00000097604 | 0.983489847 | 7.87E-05 |
| C3ar1 | ENSMUSG00000040552 | 0.980828606 | 0.03643526 |
| Igf2bp1 | ENSMUSG00000013415 | 0.977306549 | 0.00838845 |
| Cmtm7 | ENSMUSG00000032436 | 0.960927924 | 3.62E-05 |
| Mgst1 | ENSMUSG00000008540 | 0.959356606 | 3.44E-09 |
| Fbxo43 | ENSMUSG00000048230 | 0.956066473 | 0.00154328 |
| Prokr1 | ENSMUSG00000049409 | 0.954122176 | 0.01474203 |

| | | | |
|---|---|---|---|
| Zfp493 | ENSMUSG00000090659 | 0.953585951 | 3.95E-05 |
| Rgs13 | ENSMUSG00000051079 | 0.953430566 | 0.00203825 |
| 9930120I10R | ENSMUSG00000108120 | 0.948153158 | 0.0425777 |
| Creb3l4 | ENSMUSG00000027938 | 0.947938211 | 0.00740055 |
| Ifitm2 | ENSMUSG00000060591 | 0.946441783 | 3.76E-11 |
| C1rl | ENSMUSG00000038527 | 0.945839495 | 0.02986711 |
| Gm13835 | ENSMUSG00000086922 | 0.940846281 | 4.33E-06 |
| Gypc | ENSMUSG00000090523 | 0.939084001 | 0.00338326 |
| Fgfr3 | ENSMUSG00000054252 | 0.933561021 | 0.000725 |
| Nid1 | ENSMUSG00000005397 | 0.931835878 | 0.00610575 |
| Hpgds | ENSMUSG00000029919 | 0.931074286 | 0.01135332 |
| Cd68 | ENSMUSG00000018774 | 0.929906885 | 0.03330249 |
| Tbx21 | ENSMUSG00000001444 | 0.92929326 | 0.00034148 |
| Cdc37l1 | ENSMUSG00000024780 | 0.928987836 | 5.59E-35 |
| Pon3 | ENSMUSG00000029759 | 0.928811902 | 2.02E-05 |
| Slc16a10 | ENSMUSG00000019838 | 0.928419879 | 3.17E-05 |
| Gpr37l1 | ENSMUSG00000026424 | 0.926549157 | 0.00091889 |
| Rsg1 | ENSMUSG00000073733 | 0.923682673 | 0.02187159 |
| Cyp4f16 | ENSMUSG00000048440 | 0.9225911 | 1.62E-06 |
| Slc5a5 | ENSMUSG00000000792 | 0.922457361 | 3.32E-10 |
| Rdh1 | ENSMUSG00000089789 | 0.922002531 | 0.01105472 |
| Mdfi | ENSMUSG00000032717 | 0.921753899 | 0.000209 |
| Fstl1 | ENSMUSG00000022816 | 0.921403692 | 6.23E-12 |
| Sgo1 | ENSMUSG00000023940 | 0.920370035 | 0.02793278 |
| Rpl35a | ENSMUSG00000060636 | 0.918976055 | 2.28E-19 |
| Anxa5 | ENSMUSG00000027712 | 0.918951093 | 9.78E-19 |
| Calca | ENSMUSG00000030669 | 0.915925701 | 0.01241533 |
| Cfap70 | ENSMUSG00000039543 | 0.910461366 | 8.84E-05 |
| Rnase4 | ENSMUSG00000021876 | 0.909971145 | 0.0035933 |
| C2cd6 | ENSMUSG00000072295 | 0.908800873 | 0.00244161 |
| Slc35f3 | ENSMUSG00000057060 | 0.906079169 | 2.18E-05 |
| Iqcg | ENSMUSG00000035578 | 0.905265677 | 1.03E-19 |
| Angptl2 | ENSMUSG00000004105 | 0.904556735 | 0.00012213 |
| Cyp2j9 | ENSMUSG00000015224 | 0.904282761 | 0.00034649 |
| Gm43625 | ENSMUSG00000105636 | 0.902541137 | 6.86E-05 |
| Gja1 | ENSMUSG00000050953 | 0.901632998 | 0.01521701 |
| Aldh1a7 | ENSMUSG00000024747 | 0.891878054 | 3.72E-13 |
| Tlr2 | ENSMUSG00000027995 | 0.88907075 | 0.00661575 |
| Crocc2 | ENSMUSG00000084989 | 0.888971486 | 0.0118343 |
| Gpr34 | ENSMUSG00000040229 | 0.887932318 | 0.0224597 |
| F2rl1 | ENSMUSG00000021678 | 0.887774549 | 0.01206253 |
| Nr4a3 | ENSMUSG00000028341 | 0.887441293 | 0.00341143 |
| Kcns3 | ENSMUSG00000043673 | 0.886680271 | 0.00478429 |
| Rac2 | ENSMUSG00000033220 | 0.884789389 | 0.0447272 |
| Dmrta2 | ENSMUSG00000047143 | 0.882271707 | 0.02614811 |

| | | | |
|---|---|---|---|
| Acss1 | ENSMUSG00000027452 | 0.875799031 | 0.02092699 |
| Calml4 | ENSMUSG00000032246 | 0.875545226 | 6.12E-09 |
| Itga6 | ENSMUSG00000027111 | 0.874797958 | 1.43E-09 |
| Nat8f4 | ENSMUSG00000068299 | 0.873416101 | 0.00148062 |
| Ccdc80 | ENSMUSG00000022665 | 0.872369723 | 5.02E-05 |
| Spon2 | ENSMUSG00000037379 | 0.871073605 | 1.75E-10 |
| Smagp | ENSMUSG00000053559 | 0.867269392 | 0.01443947 |
| Fcer1g | ENSMUSG00000058715 | 0.865797897 | 0.03970787 |
| Gm26954 | ENSMUSG00000098233 | 0.864878146 | 1.64E-10 |
| Rnf43 | ENSMUSG00000034177 | 0.863923071 | 0.01995216 |
| Emp2 | ENSMUSG00000022505 | 0.86273157 | 6.51E-11 |
| Tmem37 | ENSMUSG00000050777 | 0.862412199 | 3.40E-05 |
| Ctgf | ENSMUSG00000019997 | 0.861809412 | 6.81E-07 |
| Gm12247 | ENSMUSG00000081223 | 0.860326465 | 0.00018732 |
| Slc15a2 | ENSMUSG00000022899 | 0.860240247 | 6.17E-34 |
| Gm14296 | ENSMUSG00000074527 | 0.859641869 | 0.00707754 |
| Hacd4 | ENSMUSG00000028497 | 0.855949023 | 0.00012545 |
| Sfrp1 | ENSMUSG00000031548 | 0.854869892 | 0.00551294 |
| Akr1c13 | ENSMUSG00000021213 | 0.853215972 | 0.04125202 |
| Zic1 | ENSMUSG00000032368 | 0.852381046 | 0.0001068 |
| Mtap | ENSMUSG00000062937 | 0.85216434 | 3.44E-07 |
| Sspn | ENSMUSG00000030255 | 0.850147071 | 6.08E-07 |
| Gfpt2 | ENSMUSG00000020363 | 0.84786018 | 0.00227043 |
| Dtd1 | ENSMUSG00000027430 | 0.846058865 | 6.59E-14 |
| Zfp457 | ENSMUSG00000055341 | 0.844589414 | 0.00068412 |
| Rarres2 | ENSMUSG00000009281 | 0.843843295 | 6.75E-05 |
| Bche | ENSMUSG00000027792 | 0.843789906 | 1.55E-09 |
| Enpp1 | ENSMUSG00000037370 | 0.842126522 | 0.00013925 |
| Txnip | ENSMUSG00000038393 | 0.841887213 | 0.0001187 |
| Hist1h2bc | ENSMUSG00000018102 | 0.841745908 | 9.94E-09 |
| Ltbp1 | ENSMUSG00000001870 | 0.837608829 | 0.00043299 |
| Tmsb15l | ENSMUSG00000072955 | 0.833469337 | 0.01489633 |
| Lgals3 | ENSMUSG00000050335 | 0.832827307 | 2.20E-08 |
| Plekhf1 | ENSMUSG00000074170 | 0.82859868 | 4.08E-05 |
| Gm17634 | ENSMUSG00000097482 | 0.827418623 | 0.00260948 |
| Sla | ENSMUSG00000022372 | 0.826749206 | 0.03030304 |
| Psmc3ip | ENSMUSG00000019303 | 0.824511175 | 0.0002005 |
| Ndufa4l2 | ENSMUSG00000040280 | 0.822001567 | 3.38E-05 |
| Myo1f | ENSMUSG00000024300 | 0.821625103 | 0.04926767 |
| Atp1a2 | ENSMUSG00000007097 | 0.820246224 | 1.13E-08 |
| Paqr6 | ENSMUSG00000041423 | 0.818377537 | 0.00277284 |
| Zfp429 | ENSMUSG00000078994 | 0.817660767 | 0.00786133 |
| Ccnd2 | ENSMUSG00000000184 | 0.817209782 | 9.45E-05 |
| Rhod | ENSMUSG00000041845 | 0.817197662 | 1.31E-07 |
| Gm37313 | ENSMUSG00000104097 | 0.817109521 | 0.02886891 |

| | | | |
|---|---|---|---|
| Fcgr3 | ENSMUSG00000059498 | 0.814258259 | 0.02231543 |
| AI464131 | ENSMUSG00000046312 | 0.81252743 | 4.13E-07 |
| Rab7b | ENSMUSG00000052688 | 0.81181878 | 0.00039541 |
| Gm3898 | ENSMUSG00000096322 | 0.810166619 | 8.43E-05 |
| Gm14295 | ENSMUSG00000078877 | 0.809744425 | 0.00049405 |
| Rassf3 | ENSMUSG00000025795 | 0.809144703 | 5.25E-09 |
| Mstn | ENSMUSG00000026100 | 0.806427884 | 2.09E-06 |
| Col18a1 | ENSMUSG00000001435 | 0.805157116 | 0.0404106 |
| Rapsn | ENSMUSG00000002104 | 0.804644219 | 0.00243454 |
| Ptgfr | ENSMUSG00000028036 | 0.80074999 | 0.04027961 |
| Fbxl8 | ENSMUSG00000033313 | 0.800534482 | 0.00883114 |
| Hist2h3c2 | ENSMUSG00000081058 | 0.800008035 | 0.00041462 |
| Pdk1 | ENSMUSG00000006494 | 0.79908467 | 3.10E-24 |
| P2ry12 | ENSMUSG00000036353 | 0.79892026 | 0.000531 |
| Nupr1 | ENSMUSG00000030717 | 0.798888859 | 0.00013923 |
| Fancg | ENSMUSG00000028453 | 0.798175135 | 2.84E-07 |
| Mki67 | ENSMUSG00000031004 | 0.798122757 | 0.01128756 |
| Cfap126 | ENSMUSG00000026649 | 0.796036414 | 2.95E-05 |
| 4930523C07I | ENSMUSG00000090394 | 0.794644732 | 0.00343206 |
| Igfbp7 | ENSMUSG00000036256 | 0.792993356 | 4.05E-05 |
| Cd44 | ENSMUSG00000005087 | 0.789963393 | 1.08E-05 |
| Egfem1 | ENSMUSG00000063600 | 0.786861441 | 1.37E-08 |
| Plcd4 | ENSMUSG00000026173 | 0.786165265 | 1.29E-11 |
| Antxr1 | ENSMUSG00000033420 | 0.780545812 | 4.67E-09 |
| Calr4 | ENSMUSG00000028558 | 0.776928381 | 0.0240195 |
| Elf4 | ENSMUSG00000031103 | 0.775297859 | 0.04406086 |
| Gm20633 | ENSMUSG00000093553 | 0.77158042 | 0.00192018 |
| Ass1 | ENSMUSG00000076441 | 0.771089837 | 4.51E-19 |
| Hpx | ENSMUSG00000030895 | 0.770027029 | 0.01568274 |
| Tdrd5 | ENSMUSG00000060985 | 0.767799587 | 0.00499933 |
| Zbtb8b | ENSMUSG00000048485 | 0.763629978 | 1.70E-08 |
| Tnfrsf10b | ENSMUSG00000022074 | 0.762717692 | 0.00133978 |
| Col4a5 | ENSMUSG00000031274 | 0.759786081 | 0.00018587 |
| Psmb9 | ENSMUSG00000096727 | 0.759530414 | 0.01497576 |
| Ifitm3 | ENSMUSG00000025492 | 0.758566168 | 3.84E-06 |
| Gm4366 | ENSMUSG00000107383 | 0.757621996 | 2.75E-15 |
| Slc29a1 | ENSMUSG00000023942 | 0.756884661 | 0.01853492 |
| Gm11961 | ENSMUSG00000056288 | 0.756697766 | 0.00499933 |
| Fhdc1 | ENSMUSG00000041842 | 0.756364193 | 2.49E-08 |
| Loxl1 | ENSMUSG00000032334 | 0.755923928 | 0.00031677 |
| Gm20632 | ENSMUSG00000093577 | 0.753285783 | 7.87E-11 |
| Tspo | ENSMUSG00000041736 | 0.750605322 | 1.70E-07 |
| Scn7a | ENSMUSG00000034810 | 0.750396252 | 3.04E-06 |
| 1700067K01I | ENSMUSG00000046408 | 0.748049018 | 0.04980118 |
| Cenpf | ENSMUSG00000026605 | 0.74756641 | 0.01869037 |

| | | | |
|---|---|---|---|
| Clec18a | ENSMUSG00000033633 | 0.745123856 | 0.0005277 |
| Nkd2 | ENSMUSG00000021567 | 0.742514976 | 0.0003696 |
| Gm44250 | ENSMUSG00000107881 | 0.739956093 | 0.00209473 |
| Bmp4 | ENSMUSG00000021835 | 0.739085424 | 0.00065171 |
| P2ry6 | ENSMUSG00000048779 | 0.738977948 | 0.04436024 |
| Metap1d | ENSMUSG00000041921 | 0.736408035 | 1.61E-07 |
| Gm37904 | ENSMUSG00000103073 | 0.735835274 | 0.019106 |
| Ccnb1 | ENSMUSG00000041431 | 0.73518044 | 0.03373068 |
| Tk1 | ENSMUSG00000025574 | 0.734829134 | 0.01490851 |
| Rny3 | ENSMUSG00000064945 | 0.733877377 | 0.04018942 |
| Gprc5c | ENSMUSG00000051043 | 0.732522334 | 0.00212863 |
| Rps2-ps10 | ENSMUSG00000091957 | 0.731131602 | 0.000471 |
| Chmp4c | ENSMUSG00000027536 | 0.730690371 | 0.02383059 |
| Gm42555 | ENSMUSG00000104701 | 0.73009742 | 0.0005242 |
| Krt18 | ENSMUSG00000023043 | 0.729153852 | 2.70E-11 |
| Palm3 | ENSMUSG00000047986 | 0.724745238 | 0.00167122 |
| Cebpb | ENSMUSG00000056501 | 0.724255096 | 0.00386757 |
| Lrrc66 | ENSMUSG00000067206 | 0.721972688 | 0.00010973 |
| Gm1840 | ENSMUSG00000043192 | 0.719574951 | 8.05E-10 |
| Tmem59l | ENSMUSG00000035964 | 0.719335521 | 1.80E-12 |
| Sema3d | ENSMUSG00000040254 | 0.718050242 | 0.00015882 |
| Slc11a1 | ENSMUSG00000026177 | 0.717908445 | 0.0100912 |
| Slc25a45 | ENSMUSG00000024818 | 0.716710543 | 0.02842455 |
| Cebpa | ENSMUSG00000034957 | 0.713447857 | 0.01568274 |
| Lpar2 | ENSMUSG00000031861 | 0.713346058 | 0.00099848 |
| Gpc4 | ENSMUSG00000031119 | 0.711313638 | 1.54E-06 |
| 1110019D14 | ENSMUSG00000097616 | 0.709362493 | 0.00959898 |
| Cp | ENSMUSG00000003617 | 0.708888869 | 3.86E-08 |
| Lrrc55 | ENSMUSG00000075224 | 0.70729457 | 0.00476568 |
| Ror2 | ENSMUSG00000021464 | 0.706898931 | 0.02525958 |
| Ckm | ENSMUSG00000030399 | 0.706549802 | 0.03610278 |
| Myl9 | ENSMUSG00000067818 | 0.705971152 | 8.49E-12 |
| Rn7sk | ENSMUSG00000065037 | 0.704412707 | 0.00389739 |
| Hexb | ENSMUSG00000021665 | 0.703611509 | 0.00013925 |
| Mecom | ENSMUSG00000027684 | 0.702202731 | 0.00547585 |
| Mei4 | ENSMUSG00000043289 | 0.70196545 | 0.04712595 |
| Kitl | ENSMUSG00000019966 | 0.701416132 | 1.05E-15 |
| Frk | ENSMUSG00000019779 | 0.701322807 | 0.02386451 |
| Nme1 | ENSMUSG00000037601 | 0.700971446 | 4.71E-13 |
| Fam60a | ENSMUSG00000039985 | 0.700824501 | 3.67E-05 |
| Cyba | ENSMUSG00000006519 | 0.696929368 | 1.70E-05 |
| Kdelr3 | ENSMUSG00000010830 | 0.695671608 | 0.004485 |
| Gm37090 | ENSMUSG00000103183 | 0.695042011 | 0.00533924 |
| Bace2 | ENSMUSG00000040605 | 0.694630768 | 0.0001508 |
| Ctsh | ENSMUSG00000032359 | 0.694126793 | 1.35E-07 |

| | | | |
|---|---|---|---|
| Tectb | ENSMUSG00000024979 | 0.693755946 | 0.03500763 |
| Aldh1a1 | ENSMUSG00000053279 | 0.691998349 | 1.75E-06 |
| Gpx8 | ENSMUSG00000021760 | 0.691421425 | 1.06E-14 |
| 9630028B13I | ENSMUSG00000051295 | 0.689510987 | 9.54E-08 |
| Hhex | ENSMUSG00000024986 | 0.689495158 | 0.02739009 |
| Selenop | ENSMUSG00000064373 | 0.68869521 | 6.97E-06 |
| Agmo | ENSMUSG00000050103 | 0.68752633 | 0.04928382 |
| Vcp | ENSMUSG00000028452 | 0.687173204 | 4.64E-24 |
| Dse | ENSMUSG00000039497 | 0.684687083 | 0.0008346 |
| Zfp811 | ENSMUSG00000055202 | 0.684367366 | 4.55E-06 |
| 9630001P10I | ENSMUSG00000097825 | 0.68388658 | 0.0030967 |
| Efemp1 | ENSMUSG00000020467 | 0.682933524 | 1.98E-10 |
| S100a10 | ENSMUSG00000041959 | 0.68257472 | 5.82E-09 |
| Podn | ENSMUSG00000028600 | 0.682271271 | 0.00156611 |
| Fam107a | ENSMUSG00000021750 | 0.682192083 | 0.03033048 |
| Pcolce2 | ENSMUSG00000015354 | 0.681615662 | 1.56E-07 |
| Cpxm1 | ENSMUSG00000027408 | 0.677600783 | 6.55E-08 |
| Pqlc3 | ENSMUSG00000045679 | 0.675354149 | 0.02309991 |
| Gm20139 | ENSMUSG00000111052 | 0.674789738 | 6.88E-07 |
| Abca9 | ENSMUSG00000041797 | 0.673314256 | 0.00305537 |
| Stbd1 | ENSMUSG00000047963 | 0.673154294 | 1.30E-05 |
| Mrap2 | ENSMUSG00000042761 | 0.673132697 | 4.22E-07 |
| Nexn | ENSMUSG00000039103 | 0.67303593 | 0.04275988 |
| Cd38 | ENSMUSG00000029084 | 0.672723992 | 0.00582417 |
| Atp1b3 | ENSMUSG00000032412 | 0.670376039 | 3.75E-10 |
| Matn2 | ENSMUSG00000022324 | 0.668640936 | 0.00193311 |
| Gm10069 | ENSMUSG00000059659 | 0.667391256 | 0.00907035 |
| Ahnak | ENSMUSG00000069833 | 0.667311148 | 0.00172803 |
| P2ry2 | ENSMUSG00000032860 | 0.666811246 | 0.01632443 |
| S100a11 | ENSMUSG00000027907 | 0.666569361 | 0.00081494 |
| Tnfaip8l2 | ENSMUSG00000013707 | 0.66553547 | 0.01473078 |
| Tal1 | ENSMUSG00000028717 | 0.665334461 | 0.03271533 |
| Crabp2 | ENSMUSG00000004885 | 0.664680983 | 0.00390948 |
| Lgi2 | ENSMUSG00000039252 | 0.663732033 | 1.65E-06 |
| Crip1 | ENSMUSG00000006360 | 0.663030455 | 0.01127085 |
| Rasef | ENSMUSG00000043003 | 0.661868881 | 0.01693709 |
| Adm | ENSMUSG00000030790 | 0.661363993 | 0.01284728 |
| Klhl6 | ENSMUSG00000043008 | 0.661117263 | 0.01709747 |
| Pxdn | ENSMUSG00000020674 | 0.660743742 | 3.41E-06 |
| Scd1 | ENSMUSG00000037071 | 0.659453835 | 7.65E-08 |
| A730056A06 | ENSMUSG00000097756 | 0.659170592 | 0.0099021 |
| 4930412C18I | ENSMUSG00000085558 | 0.655692056 | 0.02046971 |
| Emp3 | ENSMUSG00000040212 | 0.655145496 | 0.00931323 |
| Rpl22l1 | ENSMUSG00000039221 | 0.650451604 | 4.31E-07 |
| Spi1 | ENSMUSG00000002111 | 0.646500862 | 0.04149998 |

| | | | |
|---|---|---|---|
| Fgfr2 | ENSMUSG00000030849 | 0.644932484 | 0.00478497 |
| Fbln7 | ENSMUSG00000027386 | 0.644634516 | 0.00013735 |
| Rcn3 | ENSMUSG00000019539 | 0.644219313 | 7.98E-06 |
| Gm17057 | ENSMUSG00000090500 | 0.643484895 | 0.01697597 |
| Gm44751 | ENSMUSG00000109244 | 0.641512665 | 0.04487654 |
| Gm16062 | ENSMUSG00000087249 | 0.640190737 | 0.0023925 |
| Ramp1 | ENSMUSG00000034353 | 0.639035946 | 0.0002462 |
| Tst | ENSMUSG00000044986 | 0.638926669 | 0.00318156 |
| Glb1 | ENSMUSG00000045594 | 0.637938811 | 1.19E-09 |
| Hist1h4i | ENSMUSG00000060639 | 0.634084522 | 0.00754005 |
| A930037H05 | ENSMUSG00000109408 | 0.633833661 | 3.31E-05 |
| Il1r1 | ENSMUSG00000026072 | 0.6334096 | 4.40E-05 |
| Tec | ENSMUSG00000029217 | 0.633350154 | 0.01065947 |
| Hmcn1 | ENSMUSG00000066842 | 0.633256495 | 0.02942465 |
| Zcchc9 | ENSMUSG00000021621 | 0.632953226 | 2.35E-12 |
| Tcp11l1 | ENSMUSG00000027175 | 0.632854828 | 0.00316604 |
| Mfap2 | ENSMUSG00000060572 | 0.63265134 | 2.16E-05 |
| Gm45472 | ENSMUSG00000109669 | 0.632445292 | 6.13E-07 |
| Gpr68 | ENSMUSG00000047415 | 0.631588593 | 0.00240332 |
| 1700027J19F | ENSMUSG00000063838 | 0.63140802 | 0.01284224 |
| Gm26840 | ENSMUSG00000097728 | 0.630630878 | 0.0124636 |
| Colgalt2 | ENSMUSG00000032649 | 0.630489529 | 1.14E-10 |
| Ackr3 | ENSMUSG00000044337 | 0.630385513 | 4.88E-05 |
| C330013E15I | ENSMUSG00000097093 | 0.63006406 | 0.02898672 |
| Col9a1 | ENSMUSG00000026147 | 0.629593123 | 7.79E-07 |
| Fxyd6 | ENSMUSG00000066705 | 0.629202382 | 8.84E-05 |
| Hspe1 | ENSMUSG00000073676 | 0.628098425 | 3.07E-07 |
| Baiap2l1 | ENSMUSG00000038859 | 0.62797116 | 0.04558007 |
| Frmpd4 | ENSMUSG00000049176 | 0.627944821 | 0.00048614 |
| Mmp2 | ENSMUSG00000031740 | 0.627884766 | 5.10E-06 |
| Ech1 | ENSMUSG00000053898 | 0.62665877 | 1.54E-14 |
| En2 | ENSMUSG00000039095 | 0.624080247 | 0.02401918 |
| Sstr2 | ENSMUSG00000047904 | 0.623758705 | 3.84E-06 |
| Smco4 | ENSMUSG00000058173 | 0.623694266 | 0.01154656 |
| Peg10 | ENSMUSG00000092035 | 0.623013619 | 0.00397014 |
| Zfp605 | ENSMUSG00000023284 | 0.622729561 | 7.03E-05 |
| Spry3 | ENSMUSG00000061654 | 0.622626015 | 5.90E-05 |
| Slc8b1 | ENSMUSG00000032754 | 0.620628192 | 3.31E-05 |
| Cpa2 | ENSMUSG00000071553 | 0.619404216 | 1.24E-06 |
| Tnfaip8 | ENSMUSG00000062210 | 0.619110067 | 4.28E-10 |
| Itpr3 | ENSMUSG00000042644 | 0.61858566 | 0.0174185 |
| Ctsc | ENSMUSG00000030560 | 0.617365811 | 1.74E-08 |
| Sgsh | ENSMUSG00000005043 | 0.617302765 | 2.44E-07 |
| Pmfbp1 | ENSMUSG00000031727 | 0.615881341 | 0.01378475 |
| Tnfrsf12a | ENSMUSG00000023905 | 0.614999667 | 0.00097249 |

| | | | |
|---|---|---|---|
| Mlc1 | ENSMUSG00000035805 | 0.614044318 | 1.18E-14 |
| Hebp2 | ENSMUSG00000019853 | 0.613551677 | 1.41E-05 |
| Pign | ENSMUSG00000056536 | 0.613465901 | 2.29E-06 |
| Slc6a13 | ENSMUSG00000030108 | 0.612542905 | 0.00025929 |
| Rrm2 | ENSMUSG00000020649 | 0.610398915 | 0.0262763 |
| Tuba4a | ENSMUSG00000026202 | 0.610048055 | 0.04878346 |
| Tmbim1 | ENSMUSG00000006301 | 0.607409978 | 4.22E-09 |
| Idi1 | ENSMUSG00000058258 | 0.60564272 | 1.58E-08 |
| Gm6863 | ENSMUSG00000043483 | 0.604298112 | 0.01811444 |
| Ptgs1 | ENSMUSG00000047250 | 0.601733294 | 0.040691 |
| Folh1 | ENSMUSG00000001773 | 0.601686986 | 0.00879315 |
| Ppic | ENSMUSG00000024538 | 0.601602874 | 6.70E-06 |
| Gm17207 | ENSMUSG00000090561 | 0.601025377 | 0.00445304 |
| S1pr3 | ENSMUSG00000067586 | 0.600515479 | 6.13E-05 |
| Lysmd1 | ENSMUSG00000053769 | 0.600452799 | 3.40E-07 |
| Kcnip4 | ENSMUSG00000029088 | 0.599714577 | 3.31E-16 |
| Igf2 | ENSMUSG00000048583 | 0.598124804 | 3.30E-17 |
| Iyd | ENSMUSG00000019762 | 0.596884847 | 0.02762146 |
| Arhgdib | ENSMUSG00000030220 | 0.596405679 | 0.00076883 |
| Rsph3b | ENSMUSG00000023806 | 0.595813226 | 0.00050865 |
| NA | ENSMUSG00000102306 | 0.594892768 | 0.02560094 |
| Traf3ip3 | ENSMUSG00000037318 | 0.594348455 | 0.04342472 |
| Csf1 | ENSMUSG00000014599 | 0.591530538 | 0.00459021 |
| Gm13340 | ENSMUSG00000083563 | 0.591083513 | 0.01305195 |
| Ctsk | ENSMUSG00000028111 | 0.589596718 | 0.04154892 |
| Arpc1b | ENSMUSG00000029622 | 0.589486804 | 1.01E-06 |
| Gm28438 | ENSMUSG00000101939 | 0.587380304 | 0.01041907 |
| Gm10033 | ENSMUSG00000110444 | 0.586599499 | 2.20E-05 |
| Sdc1 | ENSMUSG00000020592 | 0.586488032 | 0.02211185 |
| Mrc2 | ENSMUSG00000020695 | 0.585414443 | 0.00307837 |
| Apoe | ENSMUSG00000002985 | 0.584954013 | 8.91E-11 |
| Insm2 | ENSMUSG00000045440 | 0.584921791 | 0.02800657 |
| Hfe | ENSMUSG00000006611 | 0.584042062 | 3.75E-09 |
| Ccdc180 | ENSMUSG00000035539 | 0.584002294 | 0.01939658 |
| Ppp1r3b | ENSMUSG00000046794 | 0.583899861 | 0.0067462 |
| Iqgap2 | ENSMUSG00000021676 | 0.582390192 | 0.00552437 |
| Olfml2a | ENSMUSG00000046618 | 0.581403347 | 0.04022057 |
| 4632428C04I | ENSMUSG00000097184 | 0.581332777 | 0.03787327 |
| Fign | ENSMUSG00000075324 | 0.581331339 | 0.01173101 |
| Ecm1 | ENSMUSG00000028108 | 0.580854177 | 0.0005673 |
| Fzd10 | ENSMUSG00000081683 | 0.580662822 | 0.00039502 |
| Itih5 | ENSMUSG00000025780 | 0.580188802 | 0.00051732 |
| Sqor | ENSMUSG00000005803 | 0.579175402 | 0.004745 |
| Irak1bp1 | ENSMUSG00000032251 | 0.578062185 | 0.00297862 |
| Tgif1 | ENSMUSG00000047407 | 0.577446318 | 0.00316987 |

| | | | |
|---|---|---|---|
| Pomc | ENSMUSG00000020660 | 0.577220493 | 0.00048614 |
| BC022687 | ENSMUSG00000037594 | 0.573564711 | 2.75E-08 |
| mt-Te | ENSMUSG00000064369 | 0.572205155 | 0.00262433 |
| Zfp36 | ENSMUSG00000044786 | 0.570392991 | 0.00100244 |
| Cst3 | ENSMUSG00000027447 | 0.56866207 | 1.95E-08 |
| Moxd1 | ENSMUSG00000020000 | 0.56861417 | 2.45E-07 |
| Rwdd3 | ENSMUSG00000028133 | 0.568262512 | 4.43E-05 |
| Arhgap25 | ENSMUSG00000030047 | 0.568110115 | 0.04662058 |
| Rab27b | ENSMUSG00000024511 | 0.567343773 | 1.13E-12 |
| Sptssb | ENSMUSG00000043461 | 0.567068281 | 0.00091115 |
| Cnmd | ENSMUSG00000022025 | 0.566117118 | 0.00215004 |
| Igfbp2 | ENSMUSG00000039323 | 0.565420238 | 6.10E-06 |
| Zfp758 | ENSMUSG00000044501 | 0.564847382 | 0.00052589 |
| Cmtm8 | ENSMUSG00000041012 | 0.564191223 | 0.00043585 |
| Tnfrsf1b | ENSMUSG00000028599 | 0.562224688 | 0.03163298 |
| Fam114a1 | ENSMUSG00000029185 | 0.561421077 | 0.00083438 |
| Col9a2 | ENSMUSG00000028626 | 0.558834875 | 2.32E-05 |
| Nhsl1 | ENSMUSG00000039835 | 0.55861205 | 0.0002128 |
| Ifi30 | ENSMUSG00000031838 | 0.558556317 | 5.86E-05 |
| 2900097C17I | ENSMUSG00000102869 | 0.558278691 | 1.15E-06 |
| Gas5 | ENSMUSG00000053332 | 0.557821064 | 8.24E-06 |
| Gm15500 | ENSMUSG00000086583 | 0.556513801 | 0.04826202 |
| Bambi | ENSMUSG00000024232 | 0.556284457 | 5.04E-07 |
| Cgn | ENSMUSG00000068876 | 0.555910899 | 0.0001018 |
| Itm2a | ENSMUSG00000031239 | 0.555554936 | 2.49E-09 |
| Kcnq1ot1 | ENSMUSG00000101609 | 0.554989057 | 0.03655776 |
| Il10rb | ENSMUSG00000022969 | 0.554585863 | 0.00083842 |
| Gm16055 | ENSMUSG00000089890 | 0.554098261 | 0.00481377 |
| Lyn | ENSMUSG00000042228 | 0.551013622 | 0.00372338 |
| Crh | ENSMUSG00000049796 | 0.5497751 | 0.01305814 |
| C920021L13I | ENSMUSG00000080727 | 0.549631776 | 0.03695817 |
| Gng11 | ENSMUSG00000032766 | 0.546729958 | 0.0007345 |
| Stk33 | ENSMUSG00000031027 | 0.546412948 | 0.00233087 |
| Rbm47 | ENSMUSG00000070780 | 0.545173701 | 0.04336136 |
| Gm17750 | ENSMUSG00000098087 | 0.544899975 | 0.00978129 |
| Bhlhe40 | ENSMUSG00000030103 | 0.544757488 | 5.03E-07 |
| Mterf1b | ENSMUSG00000053178 | 0.543877945 | 0.03937757 |
| Cep250 | ENSMUSG00000038241 | 0.543653654 | 5.04E-09 |
| Gm43172 | ENSMUSG00000107374 | 0.543243085 | 0.03083071 |
| Aqp4 | ENSMUSG00000024411 | 0.542527436 | 5.67E-05 |
| Icam1 | ENSMUSG00000037405 | 0.542302439 | 0.04059287 |
| Crim1 | ENSMUSG00000024074 | 0.540888707 | 0.00030928 |
| Sri | ENSMUSG00000003161 | 0.539828322 | 1.73E-13 |
| F630040K05I | ENSMUSG00000097727 | 0.538887151 | 0.00046711 |
| Herc6 | ENSMUSG00000029798 | 0.538353623 | 0.03254669 |

| | | | |
|---|---|---|---|
| A930004D18 | ENSMUSG00000054057 | 0.538188127 | 9.96E-06 |
| Six2 | ENSMUSG00000024134 | 0.537989701 | 0.03093853 |
| Mrps6 | ENSMUSG00000039680 | 0.536482086 | 2.96E-06 |
| Tmem169 | ENSMUSG00000026188 | 0.535783106 | 5.97E-07 |
| Arsk | ENSMUSG00000021592 | 0.535215243 | 0.00014244 |
| Sparc | ENSMUSG00000018593 | 0.534698888 | 2.70E-21 |
| Zfp422-ps | ENSMUSG00000091515 | 0.534111854 | 0.00068412 |
| Golm1 | ENSMUSG00000021556 | 0.533589325 | 0.00732682 |
| Cnbd2 | ENSMUSG00000038085 | 0.532103702 | 1.64E-07 |
| Thbs2 | ENSMUSG00000023885 | 0.530587028 | 0.01705699 |
| Cd320 | ENSMUSG00000002308 | 0.529756015 | 8.03E-07 |
| Inhba | ENSMUSG00000041324 | 0.528448515 | 0.03451206 |
| Gusb | ENSMUSG00000025534 | 0.528349244 | 0.00012925 |
| Top2a | ENSMUSG00000020914 | 0.52800728 | 0.03019222 |
| Rorc | ENSMUSG00000028150 | 0.524979983 | 0.03055004 |
| Zfp697 | ENSMUSG00000050064 | 0.524334735 | 0.00223375 |
| Tagln2 | ENSMUSG00000026547 | 0.524144417 | 0.00486834 |
| Fkbp9 | ENSMUSG00000029781 | 0.522430293 | 8.57E-07 |
| Ahrr | ENSMUSG00000021575 | 0.520039073 | 0.0013477 |
| Zic3 | ENSMUSG00000067860 | 0.518548366 | 0.03190567 |
| Pmp22 | ENSMUSG00000018217 | 0.518310451 | 2.05E-08 |
| Cd63 | ENSMUSG00000025351 | 0.517079516 | 9.67E-10 |
| Cgnl1 | ENSMUSG00000032232 | 0.51676679 | 0.00055365 |
| Fcgrt | ENSMUSG00000003420 | 0.516472919 | 0.00124559 |
| Cavin1 | ENSMUSG00000004044 | 0.514344264 | 0.00014323 |
| Wnt5b | ENSMUSG00000030170 | 0.513667319 | 0.00013723 |
| Spata9 | ENSMUSG00000021590 | 0.513617493 | 0.04503926 |
| Marveld2 | ENSMUSG00000021636 | 0.51315534 | 0.00948718 |
| Plp2 | ENSMUSG00000031146 | 0.512758728 | 0.02421786 |
| Thbd | ENSMUSG00000074743 | 0.512581936 | 0.02810901 |
| Frzb | ENSMUSG00000027004 | 0.512270524 | 6.74E-15 |
| Anxa2 | ENSMUSG00000032231 | 0.511944245 | 5.15E-07 |
| Nrros | ENSMUSG00000052384 | 0.511454803 | 0.00881317 |
| Hist1h2be | ENSMUSG00000047246 | 0.509442917 | 0.02409815 |
| Pkp1 | ENSMUSG00000026413 | 0.509011914 | 0.00243454 |
| Dglucy | ENSMUSG00000021185 | 0.508376243 | 7.36E-05 |
| Prelp | ENSMUSG00000041577 | 0.506744501 | 0.00071721 |
| Slc22a8 | ENSMUSG00000063796 | 0.504174693 | 0.00518332 |
| Efna1 | ENSMUSG00000027954 | 0.503281376 | 0.00013067 |
| Bub1b | ENSMUSG00000040084 | 0.50266146 | 0.00547585 |
| Mfsd2a | ENSMUSG00000028655 | 0.501647802 | 8.20E-05 |
| Suclg2 | ENSMUSG00000061838 | 0.501563008 | 0.00092556 |
| Ptpn14 | ENSMUSG00000026604 | 0.501187067 | 0.01918108 |
| Gm8292 | ENSMUSG00000100215 | 0.501074827 | 0.0461822 |
| Lipa | ENSMUSG00000024781 | 0.500909301 | 1.13E-08 |

| | | | |
|---|---|---|---|
| Hspb8 | ENSMUSG00000041548 | 0.500554422 | 0.00388364 |
| Stab1 | ENSMUSG00000042286 | 0.500200658 | 0.00898819 |
| Gm13301 | ENSMUSG00000096862 | -7.294763834 | 3.76E-11 |
| Catspere | ENSMUSG00000102483 | -6.824757103 | 1.20E-09 |
| Gm42047 | ENSMUSG00000110631 | -6.721601858 | 3.90E-21 |
| Gm12407 | ENSMUSG00000095298 | -6.714470485 | 1.89E-09 |
| 1700085C21I | ENSMUSG00000100890 | -6.70747632 | 2.65E-08 |
| Gm9522 | ENSMUSG00000098021 | -6.32299058 | 1.05E-26 |
| Gm7292 | ENSMUSG00000104222 | -6.217580958 | 1.89E-113 |
| Tpm3-rs7 | ENSMUSG00000058126 | -5.885759703 | 8.49E-93 |
| Gm8399 | ENSMUSG00000043889 | -5.694662662 | 0 |
| Gm11127 | ENSMUSG00000079492 | -5.352743469 | 6.19E-14 |
| Gm11585 | ENSMUSG00000082045 | -5.235502253 | 1.73E-07 |
| Gm12892 | ENSMUSG00000083679 | -5.068012148 | 0 |
| Vcp-rs | ENSMUSG00000083327 | -5.024231388 | 0 |
| Gm35082 | ENSMUSG00000108866 | -4.776307246 | 1.02E-20 |
| Arhgap33os | ENSMUSG00000062132 | -4.672837623 | 6.88E-16 |
| Gm11942 | ENSMUSG00000094344 | -4.478195026 | 1.97E-72 |
| 4921501E09I | ENSMUSG00000023350 | -4.471447607 | 0.00086635 |
| Gm6728 | ENSMUSG00000091408 | -4.296820229 | 1.43E-10 |
| Rpsa-ps10 | ENSMUSG00000047676 | -3.895177598 | 0.00548932 |
| Gdpd3 | ENSMUSG00000030703 | -3.704505962 | 2.39E-45 |
| Sh2d2a | ENSMUSG00000028071 | -3.680911951 | 0.00077388 |
| Gm9347 | ENSMUSG00000110126 | -3.540814632 | 9.93E-09 |
| Ccr6 | ENSMUSG00000040899 | -3.448504498 | 9.24E-06 |
| Rbpj-ps3 | ENSMUSG00000079575 | -3.393153537 | 2.12E-13 |
| 1500026H17I | ENSMUSG00000097383 | -3.309866964 | 0.02393967 |
| Gm13306 | ENSMUSG00000073877 | -3.215934236 | 2.71E-18 |
| Amd-ps4 | ENSMUSG00000019836 | -3.202566876 | 1.32E-12 |
| Trp53cor1 | ENSMUSG00000085912 | -3.184060013 | 4.59E-11 |
| Rpl15-ps2 | ENSMUSG00000098915 | -3.166919805 | 0.02221781 |
| Gm5921 | ENSMUSG00000074034 | -3.045184674 | 1.28E-11 |
| Rpl17-ps5 | ENSMUSG00000081855 | -3.016755944 | 4.91E-14 |
| Tagap | ENSMUSG00000033450 | -2.88803798 | 1.65E-22 |
| Tescl | ENSMUSG00000055826 | -2.752634827 | 5.85E-08 |
| Npy4r | ENSMUSG00000048337 | -2.702920105 | 1.70E-05 |
| Pax5 | ENSMUSG00000014030 | -2.642748817 | 2.62E-11 |
| Grifin | ENSMUSG00000036586 | -2.543442328 | 0.00070019 |
| Aire | ENSMUSG00000000731 | -2.535662954 | 7.46E-22 |
| Selenbp2 | ENSMUSG00000068877 | -2.529096681 | 1.13E-11 |
| Gm8146 | ENSMUSG00000103260 | -2.502093411 | 1.96E-05 |
| Hmgb1-ps7 | ENSMUSG00000092281 | -2.501931115 | 0.02325758 |
| 1700006J14F | ENSMUSG00000034764 | -2.466006201 | 2.81E-08 |
| Gm14204 | ENSMUSG00000086496 | -2.459083461 | 1.73E-68 |
| Gm10031 | ENSMUSG00000101523 | -2.271834639 | 1.50E-23 |

| | | | |
|---|---|---|---|
| Gm42743 | ENSMUSG00000050936 | -2.220641725 | 2.39E-39 |
| D630003M21 | ENSMUSG00000037813 | -2.216174515 | 2.13E-11 |
| Alox5ap | ENSMUSG00000060063 | -2.168444026 | 1.84E-49 |
| Upk3b | ENSMUSG00000042985 | -2.163918795 | 4.31E-14 |
| Atp8b5 | ENSMUSG00000028457 | -2.137706744 | 3.80E-10 |
| Ceacam10 | ENSMUSG00000054169 | -2.076757478 | 8.29E-08 |
| Gm15387 | ENSMUSG00000082585 | -2.040987739 | 6.70E-06 |
| Hmgb1-ps3 | ENSMUSG00000081455 | -2.013334884 | 0.00033326 |
| C4a | ENSMUSG00000015451 | -1.996908206 | 3.50E-36 |
| Gm9726 | ENSMUSG00000094935 | -1.943883833 | 1.04E-10 |
| Bpifb6 | ENSMUSG00000068009 | -1.909418097 | 8.93E-22 |
| Gm30238 | ENSMUSG00000103898 | -1.900913486 | 1.16E-13 |
| Cfap99 | ENSMUSG00000109572 | -1.889697648 | 7.53E-19 |
| Tmod4 | ENSMUSG00000005628 | -1.876385392 | 9.44E-09 |
| Pkd1l3 | ENSMUSG00000048827 | -1.858133046 | 1.26E-06 |
| Bcan | ENSMUSG00000004892 | -1.846246177 | 1.64E-24 |
| Gm15513 | ENSMUSG00000086291 | -1.810571249 | 2.20E-22 |
| Clic3 | ENSMUSG00000015093 | -1.801625099 | 6.34E-14 |
| Tmem51os1 | ENSMUSG00000073728 | -1.761232841 | 0.00152747 |
| Dennd1c | ENSMUSG00000002668 | -1.741063004 | 1.99E-05 |
| Gm18180 | ENSMUSG00000099342 | -1.733313138 | 1.80E-07 |
| Lpar3 | ENSMUSG00000036832 | -1.698154261 | 0.00323232 |
| Gm6472 | ENSMUSG00000095597 | -1.690574921 | 9.13E-15 |
| Gm26614 | ENSMUSG00000097240 | -1.677890834 | 2.62E-08 |
| Gm9057 | ENSMUSG00000105156 | -1.673227499 | 0.00050239 |
| Piwil1 | ENSMUSG00000029423 | -1.673181988 | 0.00255825 |
| Gm16432 | ENSMUSG00000091476 | -1.662871718 | 1.56E-24 |
| Gpr150 | ENSMUSG00000045509 | -1.661508142 | 0.0045879 |
| Tmem144 | ENSMUSG00000027956 | -1.637307682 | 1.11E-29 |
| Gm37277 | ENSMUSG00000103280 | -1.633200673 | 3.42E-06 |
| Ryr1 | ENSMUSG00000030592 | -1.618968077 | 8.77E-21 |
| Padi1 | ENSMUSG00000025329 | -1.601660342 | 1.76E-05 |
| Hyi | ENSMUSG00000006395 | -1.596799205 | 9.74E-29 |
| Gm26512 | ENSMUSG00000097157 | -1.585940567 | 0.00218614 |
| Padi6 | ENSMUSG00000040935 | -1.526364755 | 0.00131541 |
| Ulk4 | ENSMUSG00000040936 | -1.512493436 | 6.90E-30 |
| Lgals4 | ENSMUSG00000053964 | -1.505215908 | 5.94E-20 |
| Klhl30 | ENSMUSG00000026308 | -1.501676763 | 0.00123272 |
| Efhc1 | ENSMUSG00000041809 | -1.499155219 | 4.47E-08 |
| Cyp4f13 | ENSMUSG00000024055 | -1.491463641 | 2.30E-26 |
| Gm14584 | ENSMUSG00000083798 | -1.484501663 | 1.71E-11 |
| 6330537M06 | ENSMUSG00000110575 | -1.483386128 | 2.00E-09 |
| Alms1-ps2 | ENSMUSG00000090098 | -1.48237257 | 0.02046521 |
| Prox1os | ENSMUSG00000079045 | -1.480993108 | 1.64E-19 |
| Gm14267 | ENSMUSG00000085682 | -1.466714999 | 0.00502677 |

| | | | |
|---|---|---|---|
| Prss56 | ENSMUSG00000036480 | -1.461253009 | 1.23E-36 |
| Scrg1 | ENSMUSG00000031610 | -1.455936507 | 0.00050254 |
| Gpr137b-ps | ENSMUSG00000097715 | -1.455830473 | 6.00E-24 |
| Hist1h2al | ENSMUSG00000091383 | -1.450278602 | 1.14E-47 |
| Gm14169 | ENSMUSG00000086118 | -1.447448981 | 4.54E-08 |
| Polr2k | ENSMUSG00000045996 | -1.431334143 | 1.92E-35 |
| Gm12751 | ENSMUSG00000062554 | -1.425958204 | 1.95E-11 |
| Rsph9 | ENSMUSG00000023966 | -1.417479732 | 6.24E-27 |
| Lypd5 | ENSMUSG00000030484 | -1.405624335 | 0.00759046 |
| Gje1 | ENSMUSG00000019867 | -1.39676902 | 0.00547585 |
| Gm5540 | ENSMUSG00000048916 | -1.383906842 | 0.0212456 |
| Gm26770 | ENSMUSG00000097253 | -1.37349649 | 0.00699934 |
| Gm26668 | ENSMUSG00000097410 | -1.372495916 | 0.00023261 |
| Gm15687 | ENSMUSG00000086643 | -1.353573047 | 4.74E-07 |
| Col20a1 | ENSMUSG00000016356 | -1.3512778 | 4.47E-30 |
| Vdr | ENSMUSG00000022479 | -1.343643941 | 0.00482542 |
| C430042M11 | ENSMUSG00000097254 | -1.339390463 | 0.01442969 |
| Pax7 | ENSMUSG00000028736 | -1.321886245 | 6.19E-06 |
| Fndc1 | ENSMUSG00000071984 | -1.320665706 | 9.39E-14 |
| Zfyve28 | ENSMUSG00000037224 | -1.319742401 | 1.32E-75 |
| Meioc | ENSMUSG00000051455 | -1.319298548 | 0.0227899 |
| Prss33 | ENSMUSG00000049620 | -1.319141085 | 1.66E-05 |
| Mapk15 | ENSMUSG00000063704 | -1.308741848 | 1.17E-05 |
| Gm14286 | ENSMUSG00000087445 | -1.308525362 | 0.00475449 |
| AW046200 | ENSMUSG00000110419 | -1.305158199 | 0.00011021 |
| Gm45889 | ENSMUSG00000109936 | -1.304010625 | 0.00033242 |
| Gm45629 | ENSMUSG00000110010 | -1.302380807 | 2.70E-11 |
| Gm10499 | ENSMUSG00000073403 | -1.298869031 | 0.00674917 |
| Gm17021 | ENSMUSG00000091661 | -1.296248307 | 0.03646719 |
| Gm26847 | ENSMUSG00000097526 | -1.280083316 | 2.39E-05 |
| Gm44702 | ENSMUSG00000109420 | -1.279657118 | 0.00605416 |
| Gm13031 | ENSMUSG00000087698 | -1.276731245 | 0.00287005 |
| Ccdc171 | ENSMUSG00000052407 | -1.253616039 | 1.24E-15 |
| Card14 | ENSMUSG00000013483 | -1.252460382 | 7.08E-06 |
| Itga11 | ENSMUSG00000032243 | -1.247250916 | 0.01880939 |
| Mir1968 | ENSMUSG00000088054 | -1.236036598 | 0.00413949 |
| A930015D03 | ENSMUSG00000092368 | -1.231831946 | 0.00504677 |
| Gm23130 | ENSMUSG00000077440 | -1.228003689 | 0.04297446 |
| Rec114 | ENSMUSG00000074269 | -1.224076894 | 4.59E-05 |
| Gm11772 | ENSMUSG00000085501 | -1.219785612 | 3.74E-06 |
| Loxl4 | ENSMUSG00000025185 | -1.210780998 | 2.83E-14 |
| Endou | ENSMUSG00000022468 | -1.209691799 | 0.00120927 |
| Trpv1 | ENSMUSG00000005952 | -1.2080826 | 0.02243415 |
| Trib1 | ENSMUSG00000032501 | -1.207504839 | 2.64E-17 |
| Gm26594 | ENSMUSG00000097189 | -1.20738694 | 0.00203875 |

| | | | |
|---|---|---|---|
| Stk19 | ENSMUSG00000061207 | -1.205132554 | 4.60E-38 |
| Pcsk9 | ENSMUSG00000044254 | -1.203442461 | 6.58E-40 |
| Terb1 | ENSMUSG00000052616 | -1.192319577 | 0.00162489 |
| Htr3a | ENSMUSG00000032269 | -1.190080128 | 3.13E-14 |
| Xlr3a | ENSMUSG00000057836 | -1.189701148 | 0.00298479 |
| Lime1 | ENSMUSG00000090077 | -1.179521373 | 0.01058496 |
| Gm47113 | ENSMUSG00000111465 | -1.177757358 | 0.00019249 |
| Gm39323 | ENSMUSG00000111229 | -1.174711105 | 0.00101885 |
| Ush1c | ENSMUSG00000030838 | -1.171989637 | 3.41E-06 |
| Cbs | ENSMUSG00000024039 | -1.169093342 | 1.42E-10 |
| Ccdc24 | ENSMUSG00000078588 | -1.166297595 | 4.11E-36 |
| A930006K02 | ENSMUSG00000084866 | -1.157844713 | 1.08E-05 |
| Aurka | ENSMUSG00000027496 | -1.145364461 | 4.55E-16 |
| Vpreb3 | ENSMUSG00000000903 | -1.143787171 | 0.00473116 |
| Gm15631 | ENSMUSG00000085067 | -1.138546495 | 2.95E-05 |
| Gm11762 | ENSMUSG00000087675 | -1.133048776 | 8.24E-05 |
| Gm33056 | ENSMUSG00000110917 | -1.132928482 | 0.03032882 |
| Rreb1 | ENSMUSG00000039087 | -1.128640345 | 1.15E-12 |
| A630095N17 | ENSMUSG00000096094 | -1.126684608 | 0.0053095 |
| Gm6277 | ENSMUSG00000097440 | -1.124762321 | 4.85E-09 |
| Trim12c | ENSMUSG00000057143 | -1.122692805 | 0.0001936 |
| Acr | ENSMUSG00000022622 | -1.118083297 | 0.02817838 |
| A930038B10 | ENSMUSG00000097310 | -1.116065224 | 4.55E-06 |
| Gem | ENSMUSG00000028214 | -1.112609495 | 3.03E-39 |
| Rec8 | ENSMUSG00000002324 | -1.109584903 | 0.00623423 |
| Prss32 | ENSMUSG00000048992 | -1.096024948 | 0.02512388 |
| Rpl36-ps9 | ENSMUSG00000106476 | -1.095617909 | 0.04412286 |
| Tecta | ENSMUSG00000037705 | -1.093762806 | 4.16E-05 |
| Ccr9 | ENSMUSG00000029530 | -1.088619931 | 0.00034125 |
| Prob1 | ENSMUSG00000073600 | -1.075814715 | 2.15E-17 |
| Gm6356 | ENSMUSG00000091400 | -1.072991208 | 0.00118866 |
| H1foo | ENSMUSG00000042279 | -1.06734791 | 0.03855162 |
| A730081D07 | ENSMUSG00000086693 | -1.064802579 | 0.03534763 |
| 1810021B22 | ENSMUSG00000087331 | -1.064209272 | 1.93E-05 |
| Calcrl | ENSMUSG00000059588 | -1.06255826 | 2.70E-17 |
| 4831440E17 | ENSMUSG00000097236 | -1.062115332 | 0.00744673 |
| Bspry | ENSMUSG00000028392 | -1.057315552 | 2.01E-06 |
| Gm42848 | ENSMUSG00000106888 | -1.057098091 | 0.04708363 |
| 1700069B07 | ENSMUSG00000053081 | -1.056613739 | 0.04859086 |
| AC149090.1 | ENSMUSG00000095041 | -1.055907779 | 1.14E-14 |
| U2af1l4 | ENSMUSG00000109378 | -1.052960924 | 1.47E-06 |
| Kremen2 | ENSMUSG00000040680 | -1.049513437 | 1.16E-05 |
| Krt24 | ENSMUSG00000020913 | -1.047204235 | 7.32E-06 |
| Lama3 | ENSMUSG00000024421 | -1.044360535 | 1.33E-10 |
| Stag3 | ENSMUSG00000036928 | -1.04385042 | 1.36E-08 |

| | | | |
|---|---|---|---|
| Col11a2 | ENSMUSG00000024330 | -1.039438789 | 7.89E-20 |
| Gm17315 | ENSMUSG00000097791 | -1.034008684 | 1.15E-06 |
| Zfp97 | ENSMUSG00000095990 | -1.02956866 | 1.16E-05 |
| Ermap | ENSMUSG00000028644 | -1.023189266 | 0.00045866 |
| Rnf207 | ENSMUSG00000058498 | -1.019467089 | 1.26E-16 |
| H2-Q10 | ENSMUSG00000067235 | -1.001866951 | 0.03051375 |
| Arhgef39 | ENSMUSG00000051517 | -1.000101395 | 0.00036361 |
| BC055402 | ENSMUSG00000101429 | -0.997031005 | 3.17E-05 |
| Wdr66 | ENSMUSG00000029442 | -0.995215137 | 1.40E-06 |
| Fam110a | ENSMUSG00000027459 | -0.987190883 | 3.62E-42 |
| Catsper4 | ENSMUSG00000048003 | -0.986859874 | 0.00053914 |
| Il9r | ENSMUSG00000020279 | -0.984858471 | 5.77E-11 |
| Plvap | ENSMUSG00000034845 | -0.980562775 | 9.56E-11 |
| Samd15 | ENSMUSG00000090812 | -0.979549551 | 2.39E-07 |
| Gm44954 | ENSMUSG00000109205 | -0.978818791 | 0.00563037 |
| Cfap46 | ENSMUSG00000049571 | -0.966580009 | 2.91E-08 |
| Crxos | ENSMUSG00000074365 | -0.966523022 | 3.90E-21 |
| 4930503L19R | ENSMUSG00000044906 | -0.963648015 | 1.48E-15 |
| Sspo | ENSMUSG00000029797 | -0.961063415 | 0.00388867 |
| Gm42741 | ENSMUSG00000106905 | -0.958300657 | 2.26E-10 |
| Acot1 | ENSMUSG00000072949 | -0.953252782 | 5.42E-12 |
| Samd11 | ENSMUSG00000096351 | -0.952704051 | 1.55E-19 |
| Sugct | ENSMUSG00000055137 | -0.944677079 | 0.00389739 |
| Ero1lb | ENSMUSG00000057069 | -0.944305596 | 2.77E-23 |
| Cyp2t4 | ENSMUSG00000078787 | -0.934181833 | 0.00019002 |
| Mycbpap | ENSMUSG00000039110 | -0.930716815 | 2.92E-07 |
| Lhx1os | ENSMUSG00000087211 | -0.928418758 | 4.04E-06 |
| Gna14 | ENSMUSG00000024697 | -0.925743307 | 0.00370627 |
| Gm5868 | ENSMUSG00000060204 | -0.918457464 | 5.70E-06 |
| Cdh17 | ENSMUSG00000028217 | -0.914123376 | 0.00074668 |
| Ceacam2 | ENSMUSG00000054385 | -0.912548578 | 0.02947708 |
| Ptpmt1 | ENSMUSG00000063235 | -0.909807886 | 9.70E-13 |
| 6430531B16R | ENSMUSG00000073795 | -0.90786053 | 0.02284304 |
| 9030624G23 | ENSMUSG00000073158 | -0.906482548 | 0.00162311 |
| Nudt8 | ENSMUSG00000024869 | -0.903205204 | 5.64E-05 |
| Dctn3 | ENSMUSG00000028447 | -0.900143271 | 5.83E-33 |
| Cd300lf | ENSMUSG00000047798 | -0.897684112 | 0.04454403 |
| Krt20 | ENSMUSG00000035775 | -0.896650338 | 0.02104448 |
| Sycp1 | ENSMUSG00000027855 | -0.889910571 | 0.02034151 |
| 5930430L01R | ENSMUSG00000106951 | -0.889727989 | 4.80E-06 |
| Gm44645 | ENSMUSG00000108601 | -0.886566855 | 0.00056019 |
| Ring1 | ENSMUSG00000024325 | -0.884910138 | 1.19E-39 |
| Mob3b | ENSMUSG00000073910 | -0.884347601 | 2.89E-07 |
| Clcnka | ENSMUSG00000033770 | -0.883172327 | 0.00276419 |
| Ybx2 | ENSMUSG00000018554 | -0.87688882 | 1.30E-09 |

| | | | |
|---|---|---|---|
| Zfp729a | ENSMUSG00000021510 | -0.876393116 | 1.65E-22 |
| 4933431C10I | ENSMUSG00000108923 | -0.870295895 | 0.03304356 |
| Gm16299 | ENSMUSG00000089679 | -0.868499709 | 0.00099093 |
| Abca6 | ENSMUSG00000044749 | -0.86648789 | 4.35E-05 |
| Serpine3 | ENSMUSG00000091155 | -0.864087508 | 3.98E-05 |
| Gm20605 | ENSMUSG00000029720 | -0.860066616 | 4.66E-08 |
| Gm25082 | ENSMUSG00000093734 | -0.859639887 | 0.0455524 |
| Myo1d | ENSMUSG00000035441 | -0.857001795 | 1.07E-11 |
| Rps4l | ENSMUSG00000063171 | -0.856017894 | 7.16E-10 |
| Pde6b | ENSMUSG00000029491 | -0.853383039 | 7.22E-06 |
| Glns-ps1 | ENSMUSG00000082100 | -0.842287735 | 0.0463155 |
| Gm38405 | ENSMUSG00000109002 | -0.84100002 | 0.00010555 |
| Prss41 | ENSMUSG00000024114 | -0.838464119 | 6.05E-07 |
| Col27a1 | ENSMUSG00000045672 | -0.838194001 | 1.11E-13 |
| Nr1h3 | ENSMUSG00000002108 | -0.837490798 | 5.15E-05 |
| Gm5297 | ENSMUSG00000105124 | -0.836430954 | 0.02079725 |
| Draxin | ENSMUSG00000029005 | -0.835112582 | 1.06E-06 |
| Pgap2 | ENSMUSG00000030990 | -0.831797989 | 1.05E-05 |
| Clstn1 | ENSMUSG00000039953 | -0.828336899 | 3.20E-55 |
| Plekhn1 | ENSMUSG00000078485 | -0.822372774 | 8.16E-07 |
| Col8a2 | ENSMUSG00000056174 | -0.819810212 | 0.01632307 |
| 4932442E05I | ENSMUSG00000104443 | -0.817607385 | 0.02625488 |
| Gm14966 | ENSMUSG00000079467 | -0.817062713 | 0.00022746 |
| Lrat | ENSMUSG00000028003 | -0.816811778 | 3.45E-11 |
| Col6a1 | ENSMUSG00000001119 | -0.816607391 | 3.13E-09 |
| Cfap58 | ENSMUSG00000046585 | -0.808163486 | 4.59E-05 |
| Igfn1 | ENSMUSG00000051985 | -0.808015118 | 1.77E-07 |
| Ly6g6d | ENSMUSG00000073413 | -0.805271352 | 0.02262851 |
| Zfp133-ps | ENSMUSG00000083674 | -0.803891556 | 0.00769821 |
| Haus5 | ENSMUSG00000078762 | -0.798341684 | 6.65E-06 |
| Dxo | ENSMUSG00000040482 | -0.797440171 | 1.90E-28 |
| Slc45a3 | ENSMUSG00000026435 | -0.795355598 | 0.00729453 |
| Gm14681 | ENSMUSG00000081603 | -0.794658985 | 0.00711143 |
| 2900052L18I | ENSMUSG00000043993 | -0.79420576 | 3.00E-09 |
| Gm13112 | ENSMUSG00000087437 | -0.793952116 | 0.03234637 |
| Gm26852 | ENSMUSG00000097231 | -0.791767939 | 3.49E-05 |
| 8430408G22 | ENSMUSG00000048489 | -0.790387802 | 0.01442969 |
| Cldn7 | ENSMUSG00000018569 | -0.787710106 | 1.57E-06 |
| Gpsm3 | ENSMUSG00000034786 | -0.785766182 | 0.01983805 |
| H2-DMa | ENSMUSG00000037649 | -0.785759263 | 6.22E-08 |
| Gm15777 | ENSMUSG00000085269 | -0.785107264 | 0.01552025 |
| Echdc2 | ENSMUSG00000028601 | -0.784915116 | 3.35E-07 |
| 4933408B17I | ENSMUSG00000049357 | -0.783674133 | 0.02882618 |
| C2 | ENSMUSG00000024371 | -0.781562496 | 0.00586339 |
| Zfp85os | ENSMUSG00000044081 | -0.780118009 | 0.00560753 |

| | | | |
|---|---|---|---|
| Wisp3 | ENSMUSG00000062074 | -0.77802675 | 0.02061993 |
| Gm11754 | ENSMUSG00000086306 | -0.776405034 | 0.04684686 |
| Skor2 | ENSMUSG00000091519 | -0.775468603 | 3.40E-05 |
| Rab37 | ENSMUSG00000020732 | -0.774202727 | 0.00073422 |
| Il3ra | ENSMUSG00000068758 | -0.773961298 | 2.68E-06 |
| Gm13421 | ENSMUSG00000085929 | -0.773929762 | 0.00850926 |
| Sec1 | ENSMUSG00000040364 | -0.771975785 | 0.01443947 |
| Slc2a4rg-ps | ENSMUSG00000085028 | -0.770591049 | 0.00033747 |
| Prr22 | ENSMUSG00000090273 | -0.766289619 | 0.03350989 |
| Gm6733 | ENSMUSG00000084329 | -0.763658665 | 0.04367354 |
| Fam210b | ENSMUSG00000027495 | -0.758281931 | 1.54E-28 |
| Tnfsf13 | ENSMUSG00000089669 | -0.753580388 | 5.38E-10 |
| F420014N23 | ENSMUSG00000097331 | -0.7535627 | 7.08E-05 |
| Gm10857 | ENSMUSG00000102887 | -0.74856919 | 0.00527621 |
| Pla2g4b | ENSMUSG00000098488 | -0.748396742 | 0.03211981 |
| Sdcbp2 | ENSMUSG00000027456 | -0.746911593 | 0.0114404 |
| Gpc2 | ENSMUSG00000029510 | -0.746820733 | 6.88E-05 |
| H2-Q6 | ENSMUSG00000073409 | -0.745461795 | 0.04519971 |
| Atp7b | ENSMUSG00000006567 | -0.744462344 | 3.36E-13 |
| Thap6 | ENSMUSG00000102644 | -0.744325896 | 1.83E-10 |
| Gm5601 | ENSMUSG00000100153 | -0.739065738 | 0.00181564 |
| Apcdd1 | ENSMUSG00000071847 | -0.738717045 | 1.89E-26 |
| St6galnac2 | ENSMUSG00000110170 | -0.734578793 | 0.0330748 |
| Gm37764 | ENSMUSG00000102990 | -0.733556489 | 0.01249253 |
| Glb1l | ENSMUSG00000026200 | -0.727517265 | 2.61E-06 |
| Gm10644 | ENSMUSG00000074219 | -0.726887889 | 0.00016154 |
| B230206L02I | ENSMUSG00000086003 | -0.725298362 | 0.00088079 |
| Gm36210 | ENSMUSG00000110221 | -0.722609054 | 9.64E-05 |
| Rtel1 | ENSMUSG00000038685 | -0.721849452 | 1.03E-13 |
| Ccdc163 | ENSMUSG00000028689 | -0.719833686 | 1.73E-10 |
| Cdt1 | ENSMUSG00000006585 | -0.715757537 | 6.15E-07 |
| Ankrd33b | ENSMUSG00000022237 | -0.714170318 | 2.39E-10 |
| Zmynd15 | ENSMUSG00000040829 | -0.713925997 | 0.00066966 |
| 5830462I19R | ENSMUSG00000091102 | -0.7122867 | 0.03300815 |
| Ankrd23 | ENSMUSG00000067653 | -0.710984053 | 0.00602623 |
| Pcsk4 | ENSMUSG00000020131 | -0.710851066 | 1.45E-06 |
| Cfap69 | ENSMUSG00000040473 | -0.710415374 | 2.99E-07 |
| Cacna1f | ENSMUSG00000031142 | -0.707063075 | 6.13E-15 |
| Gm16982 | ENSMUSG00000097170 | -0.705834344 | 0.00348482 |
| Malt1 | ENSMUSG00000032688 | -0.702638958 | 2.64E-06 |
| Prcp | ENSMUSG00000061119 | -0.701929877 | 2.40E-08 |
| Gm36445 | ENSMUSG00000110393 | -0.700494518 | 0.01003761 |
| Doc2g | ENSMUSG00000024871 | -0.698374832 | 0.00056676 |
| Trim68 | ENSMUSG00000073968 | -0.696488039 | 3.23E-05 |
| Gstp1 | ENSMUSG00000060803 | -0.694615541 | 1.18E-10 |

| | | | |
|---|---|---|---|
| 4932422M17 | ENSMUSG00000105881 | -0.692576916 | 0.00622204 |
| Snx22 | ENSMUSG00000039452 | -0.691133707 | 0.03271533 |
| Igsf21 | ENSMUSG00000040972 | -0.689880743 | 7.17E-14 |
| Jmjd7 | ENSMUSG00000098789 | -0.687495831 | 0.0059278 |
| Gm11696 | ENSMUSG00000056687 | -0.685389596 | 3.36E-12 |
| 4833428L15R | ENSMUSG00000097074 | -0.684159779 | 0.00790209 |
| Gm29442 | ENSMUSG00000101008 | -0.683967105 | 0.0013477 |
| Tle2 | ENSMUSG00000034771 | -0.683937971 | 3.34E-07 |
| Itgb3bp | ENSMUSG00000028549 | -0.682226846 | 2.21E-05 |
| Gm13029 | ENSMUSG00000086164 | -0.680540627 | 0.00347916 |
| Ccdc57 | ENSMUSG00000048445 | -0.677480769 | 7.20E-09 |
| Mterf1a | ENSMUSG00000040429 | -0.674946028 | 0.00085275 |
| Saal1 | ENSMUSG00000006763 | -0.674916852 | 2.90E-14 |
| Pisd-ps1 | ENSMUSG00000082286 | -0.67247838 | 1.24E-05 |
| Impg2 | ENSMUSG00000035270 | -0.670192445 | 1.23E-11 |
| 9330159M07 | ENSMUSG00000097177 | -0.668310733 | 0.03534763 |
| Rps3a1 | ENSMUSG00000028081 | -0.667251588 | 7.53E-19 |
| Cyp4f17 | ENSMUSG00000091586 | -0.666340395 | 3.48E-07 |
| Xaf1 | ENSMUSG00000040483 | -0.665746786 | 0.03964004 |
| Gm26670 | ENSMUSG00000097416 | -0.664683112 | 0.00016352 |
| Cdk3-ps | ENSMUSG00000092300 | -0.664501844 | 0.04058811 |
| Zfp692 | ENSMUSG00000037243 | -0.664173457 | 3.52E-08 |
| A330074K22R | ENSMUSG00000097960 | -0.663015046 | 0.00157658 |
| Wfikkn2 | ENSMUSG00000044177 | -0.659164592 | 3.01E-06 |
| Clasrp | ENSMUSG00000061028 | -0.657840539 | 1.10E-11 |
| 2310058D17R | ENSMUSG00000085379 | -0.656459177 | 0.00018881 |
| Gm15489 | ENSMUSG00000086942 | -0.654511155 | 0.01478313 |
| Efcab2 | ENSMUSG00000026495 | -0.654178347 | 2.92E-07 |
| Mir124a-1hg | ENSMUSG00000097545 | -0.65389315 | 1.66E-06 |
| Gmip | ENSMUSG00000036246 | -0.651063754 | 8.41E-07 |
| Srp54b | ENSMUSG00000079108 | -0.648779816 | 8.22E-21 |
| Sfxn2 | ENSMUSG00000025036 | -0.648470949 | 4.40E-07 |
| Sema4c | ENSMUSG00000026121 | -0.64733993 | 1.29E-13 |
| Gm8066 | ENSMUSG00000106237 | -0.645697029 | 0.03720798 |
| NA | ENSMUSG00000091089 | -0.645105251 | 0.00054615 |
| A930003O13 | ENSMUSG00000105076 | -0.644893268 | 4.31E-06 |
| Ccdc187 | ENSMUSG00000048038 | -0.644702003 | 0.00028515 |
| Ttc34 | ENSMUSG00000046637 | -0.643033565 | 2.54E-05 |
| Tmem253 | ENSMUSG00000072571 | -0.643024761 | 0.01264457 |
| Met | ENSMUSG00000009376 | -0.64042665 | 0.00024679 |
| Gm26666 | ENSMUSG00000097419 | -0.640316659 | 0.00076081 |
| Spata6 | ENSMUSG00000034401 | -0.640228846 | 8.58E-08 |
| Zan | ENSMUSG00000079173 | -0.63944338 | 1.57E-06 |
| Fam129c | ENSMUSG00000043243 | -0.637190706 | 0.00155455 |
| Dhx35 | ENSMUSG00000027655 | -0.635867998 | 3.42E-21 |

| | | | |
|---|---|---|---|
| Rabgef1 | ENSMUSG00000025340 | -0.633994858 | 1.23E-08 |
| Igsf9 | ENSMUSG00000037995 | -0.630575812 | 7.04E-13 |
| Pvr | ENSMUSG00000040511 | -0.628296715 | 3.14E-11 |
| Fam193b | ENSMUSG00000021495 | -0.625937385 | 1.12E-09 |
| Chtf18 | ENSMUSG00000019214 | -0.62573507 | 3.84E-06 |
| Asic3 | ENSMUSG00000038276 | -0.622350262 | 2.11E-09 |
| Khsrp | ENSMUSG00000007670 | -0.621681865 | 1.41E-20 |
| Gucy2e | ENSMUSG00000020890 | -0.618986938 | 7.23E-11 |
| Stom | ENSMUSG00000026880 | -0.618691769 | 1.40E-12 |
| Gm44280 | ENSMUSG00000107944 | -0.617759144 | 0.03814794 |
| Tnfsfm13 | ENSMUSG00000018752 | -0.616956668 | 6.83E-08 |
| NA | ENSMUSG00000101767 | -0.616731097 | 0.03787369 |
| Klhl33 | ENSMUSG00000090799 | -0.616480863 | 0.00018226 |
| Rhpn1 | ENSMUSG00000022580 | -0.615656037 | 7.79E-07 |
| Smug1 | ENSMUSG00000036061 | -0.614528512 | 6.01E-22 |
| Gm15587 | ENSMUSG00000086769 | -0.613816915 | 0.00037022 |
| Ccdc13 | ENSMUSG00000079235 | -0.613589227 | 0.00130357 |
| Gprin2 | ENSMUSG00000071531 | -0.61348938 | 0.04447061 |
| Scrn2 | ENSMUSG00000020877 | -0.613114159 | 6.82E-06 |
| Scn5a | ENSMUSG00000032511 | -0.611591624 | 1.01E-06 |
| Spef2 | ENSMUSG00000072663 | -0.611432064 | 0.00732434 |
| Mzb1 | ENSMUSG00000024353 | -0.611155729 | 0.03011169 |
| Upp2 | ENSMUSG00000026839 | -0.61028347 | 0.00642985 |
| Sema3c | ENSMUSG00000028780 | -0.610022031 | 3.89E-08 |
| Ccdc155 | ENSMUSG00000038292 | -0.609653353 | 0.00121813 |
| Kif21b | ENSMUSG00000041642 | -0.608850766 | 4.07E-05 |
| Morn1 | ENSMUSG00000029049 | -0.607917111 | 5.38E-09 |
| Gm38336 | ENSMUSG00000104002 | -0.606262442 | 0.0416651 |
| Papln | ENSMUSG00000021223 | -0.60429309 | 0.00733438 |
| AC160637.1 | ENSMUSG00000111394 | -0.603187484 | 0.00017549 |
| Tead3 | ENSMUSG00000002249 | -0.596589561 | 1.48E-07 |
| Dnase1l2 | ENSMUSG00000024136 | -0.595764123 | 4.41E-05 |
| 1810012K08l | ENSMUSG00000087595 | -0.594042849 | 0.03695817 |
| Chrna2 | ENSMUSG00000022041 | -0.593923632 | 3.15E-06 |
| Disp3 | ENSMUSG00000041544 | -0.593574327 | 1.59E-06 |
| Myocd | ENSMUSG00000020542 | -0.593384177 | 0.03425046 |
| Il18bp | ENSMUSG00000070427 | -0.593340796 | 0.00930121 |
| 1700018L02l | ENSMUSG00000100075 | -0.588642531 | 8.25E-05 |
| Tnfrsf13c | ENSMUSG00000068105 | -0.588341989 | 0.04567384 |
| L3hypdh | ENSMUSG00000019718 | -0.587035308 | 9.44E-09 |
| Nmbr | ENSMUSG00000019865 | -0.585393522 | 0.00698541 |
| Hspa1l | ENSMUSG00000007033 | -0.585194935 | 0.04636931 |
| Snhg17 | ENSMUSG00000085385 | -0.583986493 | 3.47E-07 |
| Hexdc | ENSMUSG00000039307 | -0.583501761 | 5.32E-12 |
| Crocc | ENSMUSG00000040860 | -0.583261321 | 4.52E-09 |

| | | | |
|---|---|---|---|
| Slc25a47 | ENSMUSG00000048856 | -0.582495269 | 0.00668911 |
| Gm13027 | ENSMUSG00000085968 | -0.581275254 | 0.00056457 |
| Gm48677 | ENSMUSG00000110824 | -0.581052582 | 0.0359797 |
| Firre | ENSMUSG00000085396 | -0.580158419 | 1.48E-07 |
| Pdzrn3 | ENSMUSG00000035357 | -0.579041603 | 4.20E-05 |
| Iqcd | ENSMUSG00000029601 | -0.577548994 | 0.00086635 |
| Leng8 | ENSMUSG00000035545 | -0.577536215 | 4.00E-05 |
| Sh3pxd2b | ENSMUSG00000040711 | -0.577449901 | 0.0014947 |
| Kif19a | ENSMUSG00000010021 | -0.576966385 | 0.00014537 |
| Ubxn11 | ENSMUSG00000012126 | -0.576643743 | 2.67E-06 |
| Gm46136 | ENSMUSG00000111163 | -0.576552832 | 0.03868289 |
| Tmem255b | ENSMUSG00000038457 | -0.576245863 | 0.01420635 |
| Gm34655 | ENSMUSG00000110863 | -0.576157986 | 0.04173986 |
| Bcar3 | ENSMUSG00000028121 | -0.574629516 | 1.53E-06 |
| Abhd14b | ENSMUSG00000042073 | -0.574422867 | 1.29E-07 |
| Cubn | ENSMUSG00000026726 | -0.572732726 | 0.02496131 |
| 4930447C04I | ENSMUSG00000021098 | -0.572327346 | 4.36E-10 |
| Gm43012 | ENSMUSG00000105802 | -0.5693794 | 0.03525764 |
| Tmem150a | ENSMUSG00000055912 | -0.568757504 | 1.16E-06 |
| AC162938.1 | ENSMUSG00000111334 | -0.567833835 | 0.02708002 |
| 1700029I15R | ENSMUSG00000044916 | -0.567163395 | 0.03264013 |
| Ing4 | ENSMUSG00000030330 | -0.566432011 | 0.00899325 |
| Mc1r | ENSMUSG00000074037 | -0.565818866 | 5.10E-06 |
| Dyrk2 | ENSMUSG00000028630 | -0.56501104 | 0.00038939 |
| Prkab1 | ENSMUSG00000029513 | -0.562089571 | 1.47E-08 |
| Ppef2 | ENSMUSG00000029410 | -0.560839477 | 2.77E-10 |
| Plch2 | ENSMUSG00000029055 | -0.560097983 | 1.07E-13 |
| Rgs20 | ENSMUSG00000002459 | -0.559949328 | 0.00015448 |
| Kbtbd4 | ENSMUSG00000005505 | -0.559483928 | 2.23E-08 |
| Abca4 | ENSMUSG00000028125 | -0.557879456 | 9.82E-08 |
| Cables1 | ENSMUSG00000040957 | -0.557155806 | 3.57E-09 |
| Dynlrb2 | ENSMUSG00000034467 | -0.556740489 | 0.00995319 |
| Ubap1l | ENSMUSG00000086228 | -0.556414406 | 4.23E-07 |
| Polm | ENSMUSG00000020474 | -0.556215371 | 1.64E-05 |
| Mamdc4 | ENSMUSG00000026941 | -0.554644906 | 0.00147336 |
| Kcns1 | ENSMUSG00000040164 | -0.554110361 | 0.00231551 |
| Gm45767 | ENSMUSG00000110702 | -0.55310438 | 0.00027134 |
| Psd2 | ENSMUSG00000024347 | -0.552805717 | 1.41E-05 |
| Stard9 | ENSMUSG00000033705 | -0.551564202 | 0.002862 |
| Lipg | ENSMUSG00000053846 | -0.549628534 | 7.61E-05 |
| Samhd1 | ENSMUSG00000027639 | -0.549407362 | 1.41E-11 |
| 4930581F22I | ENSMUSG00000070315 | -0.548271543 | 2.05E-07 |
| Crhr2 | ENSMUSG00000003476 | -0.547200669 | 0.00351124 |
| Wisp1 | ENSMUSG00000005124 | -0.546254279 | 1.42E-05 |
| Rad54b | ENSMUSG00000078773 | -0.545872133 | 0.01839393 |

| | | | |
|---|---|---|---|
| Rexo5 | ENSMUSG00000030924 | -0.545370327 | 0.0330748 |
| Nog | ENSMUSG00000048616 | -0.544981816 | 0.01778077 |
| Wnt2b | ENSMUSG00000027840 | -0.543524963 | 5.99E-05 |
| Radil | ENSMUSG00000029576 | -0.543373417 | 2.09E-07 |
| Fank1 | ENSMUSG00000053111 | -0.541413578 | 0.04773431 |
| Plcd3 | ENSMUSG00000020937 | -0.540997294 | 2.22E-09 |
| Hmgb2 | ENSMUSG00000054717 | -0.540097896 | 2.95E-06 |
| Rfx1 | ENSMUSG00000031706 | -0.539852113 | 1.01E-08 |
| Dnah9 | ENSMUSG00000056752 | -0.539617541 | 4.64E-05 |
| A730011C13 | ENSMUSG00000084846 | -0.539526388 | 0.00137733 |
| Slc25a19 | ENSMUSG00000020744 | -0.539525912 | 6.27E-08 |
| Fscn2 | ENSMUSG00000025380 | -0.539038929 | 0.00010467 |
| Cul9 | ENSMUSG00000040327 | -0.538543859 | 8.78E-18 |
| Gigyf1 | ENSMUSG00000029714 | -0.53751981 | 1.90E-06 |
| Pfkfb2 | ENSMUSG00000026409 | -0.536841023 | 1.00E-13 |
| Cpt1a | ENSMUSG00000024900 | -0.535706174 | 7.25E-07 |
| Stx3 | ENSMUSG00000041488 | -0.53489457 | 1.29E-16 |
| Amy1 | ENSMUSG00000074264 | -0.534804193 | 0.00225038 |
| Uckl1 | ENSMUSG00000089917 | -0.531808978 | 8.47E-07 |
| Slc9a5 | ENSMUSG00000014786 | -0.530942672 | 1.96E-09 |
| Tdrd9 | ENSMUSG00000054003 | -0.530602896 | 7.31E-05 |
| Mesp1 | ENSMUSG00000030544 | -0.530538883 | 0.03140554 |
| Csf2ra | ENSMUSG00000059326 | -0.530370378 | 1.90E-06 |
| Pitpnm1 | ENSMUSG00000024851 | -0.529699694 | 1.05E-10 |
| Mbd6 | ENSMUSG00000025409 | -0.529615935 | 4.54E-06 |
| Lct | ENSMUSG00000026354 | -0.528821457 | 0.04997142 |
| Acadvl | ENSMUSG00000018574 | -0.52812498 | 2.03E-07 |
| Amh | ENSMUSG00000035262 | -0.528087593 | 0.00880436 |
| Stk11ip | ENSMUSG00000026213 | -0.528056884 | 6.79E-14 |
| AA386476 | ENSMUSG00000074357 | -0.528053886 | 0.00203557 |
| Tgfbr3l | ENSMUSG00000089736 | -0.527229943 | 0.00786133 |
| Ung | ENSMUSG00000029591 | -0.525860729 | 0.00963416 |
| Aqp6 | ENSMUSG00000043144 | -0.525020259 | 0.0058819 |
| Smad2 | ENSMUSG00000024563 | -0.524103996 | 1.75E-13 |
| 3632451O06 | ENSMUSG00000036242 | -0.523835886 | 1.26E-07 |
| Cdan1 | ENSMUSG00000027284 | -0.523772183 | 1.95E-08 |
| Robo3 | ENSMUSG00000032128 | -0.522602735 | 0.01068208 |
| C1qtnf12 | ENSMUSG00000023571 | -0.521300123 | 3.70E-08 |
| Kif24 | ENSMUSG00000028438 | -0.521224491 | 0.01663924 |
| Ttc30b | ENSMUSG00000075273 | -0.5201502 | 4.65E-09 |
| Gm3764 | ENSMUSG00000097156 | -0.518438131 | 0.00560114 |
| Nmrk1 | ENSMUSG00000037847 | -0.517679006 | 0.00755729 |
| Kifc3 | ENSMUSG00000031788 | -0.515419747 | 4.71E-12 |
| Adprhl1 | ENSMUSG00000031448 | -0.514493888 | 0.00714809 |
| Ctu2 | ENSMUSG00000049482 | -0.513684127 | 1.15E-06 |

| | | | |
|---|---|---|---|
| 6030443J06F | ENSMUSG00000097207 | -0.512330632 | 0.04891833 |
| Aipl1 | ENSMUSG00000040554 | -0.512272478 | 6.22E-06 |
| Fnbp4 | ENSMUSG00000008200 | -0.5109324 | 8.87E-11 |
| Alpk2 | ENSMUSG00000032845 | -0.510545006 | 0.0401122 |
| Caprin2 | ENSMUSG00000030309 | -0.509665963 | 0.00019951 |
| Psd | ENSMUSG00000037126 | -0.50942962 | 4.71E-13 |
| Llgl2 | ENSMUSG00000020782 | -0.507909368 | 6.43E-07 |
| Tsnaxip1 | ENSMUSG00000031893 | -0.507886288 | 0.03556437 |
| Ptchd1 | ENSMUSG00000041552 | -0.50770888 | 0.04240947 |
| Zfp541 | ENSMUSG00000078796 | -0.507383432 | 0.01060475 |
| Zscan18 | ENSMUSG00000070822 | -0.506683591 | 0.00269039 |
| Zfpm1 | ENSMUSG00000049577 | -0.506677873 | 6.83E-08 |
| Rrp1b | ENSMUSG00000058392 | -0.505780983 | 9.31E-11 |
| Ntng2 | ENSMUSG00000035513 | -0.505342781 | 1.29E-10 |
| Gm10686 | ENSMUSG00000111685 | -0.504324106 | 0.00427895 |
| Acrbp | ENSMUSG00000072770 | -0.504311284 | 0.00122068 |
| Fpgs | ENSMUSG00000009566 | -0.50420434 | 5.32E-05 |
| Cntn3 | ENSMUSG00000030075 | -0.503492492 | 0.00946314 |
| Tmem106c | ENSMUSG00000052369 | -0.50290518 | 1.02E-07 |
| BC051226 | ENSMUSG00000092564 | -0.502775795 | 0.00355602 |
| Ccdc107 | ENSMUSG00000028461 | -0.502760934 | 5.60E-06 |
| Dennd6b | ENSMUSG00000015377 | -0.500912326 | 1.72E-05 |
| Fance | ENSMUSG00000007570 | -0.500305143 | 9.38E-07 |

**Supplementary data sheet 2**

#### Differentially expressed endothelial cell (EC) genes in the P10 Apcdd1-/- retina

##### Significantly upregulated genes

| Gene_name | Ensembl_ID | log2FoldChange | padj |
|---|---|---|---|
| Arhgap8 | ENSMUSG00000078954.6 | 0.586548095 | 3.52E-06 |
| Bche | ENSMUSG00000027792.8 | 0.727919092 | 8.57E-11 |
| Calml4 | ENSMUSG00000032246.10 | 0.529095506 | 9.36E-05 |
| Ccna2 | ENSMUSG00000027715.6 | 0.972399884 | 1.38E-15 |
| Cd1d2 | ENSMUSG00000041750.10 | 0.588685848 | 3.36E-04 |
| Cdc37l1 | ENSMUSG00000024780.6 | 0.554913032 | 3.78E-10 |
| Cep250 | ENSMUSG00000038241.13 | 0.544480873 | 2.36E-08 |
| Cfi | ENSMUSG00000058952.9 | 0.564369837 | 4.01E-04 |
| Cwc22 | ENSMUSG00000027014.11 | 0.527087346 | 6.99E-04 |
| Cyp4f16 | ENSMUSG00000048440.12 | 0.943522257 | 2.52E-16 |
| Dtd1 | ENSMUSG00000027430.9 | 0.502929076 | 1.13E-07 |
| Fancg | ENSMUSG00000028453.7 | 0.578822885 | 8.06E-06 |
| Frem2 | ENSMUSG00000037016.8 | 0.899741099 | 6.28E-19 |
| Fxyd6 | ENSMUSG00000066705.6 | 0.666125749 | 4.50E-10 |
| H2-D1 | ENSMUSG00000073411.8 | 2.388354934 | 1.33E-186 |
| H2-K1 | ENSMUSG00000061232.12 | 0.920807862 | 8.63E-12 |
| H2-Q4 | ENSMUSG00000035929.11 | 1.275125296 | 5.65E-28 |
| Hebp2 | ENSMUSG00000019853.5 | 0.554960147 | 1.97E-05 |
| Hist1h2bc | ENSMUSG00000018102.4 | 0.518366657 | 9.64E-06 |
| Hs3st3a1 | ENSMUSG00000047759.6 | 0.643886902 | 2.69E-05 |
| Hspe1 | ENSMUSG00000073676.4 | 0.587872204 | 1.39E-09 |
| Ifi30 | ENSMUSG00000031838.7 | -0.533926718 | 4.35E-06 |
| Il11ra1 | ENSMUSG00000073889.7 | 1.323589324 | 1.59E-40 |
| Irak1bp1 | ENSMUSG00000032251.9 | 0.591567773 | 3.92E-08 |
| Itga6 | ENSMUSG00000027111.12 | 0.595201244 | 2.48E-11 |
| Krt18 | ENSMUSG00000023043.6 | 1.062922846 | 1.25E-22 |
| Lrpprc | ENSMUSG00000024120.9 | 0.777528424 | 2.77E-38 |
| Moxd1 | ENSMUSG00000020000.7 | 0.530092546 | 2.90E-06 |
| Mt3 | ENSMUSG00000031760.8 | 1.033404476 | 3.23E-22 |
| Ndufa4l2 | ENSMUSG00000040280.9 | 0.596559115 | 1.88E-05 |
| Npb | ENSMUSG00000044034.8 | 0.754670852 | 7.40E-06 |
| Nudt19 | ENSMUSG00000034875.5 | 0.741253607 | 1.36E-14 |
| Pdk1 | ENSMUSG00000006494.8 | 0.525088303 | 9.20E-17 |
| Pomc | ENSMUSG00000020660.5 | 0.591905084 | 8.81E-05 |
| Rab27b | ENSMUSG00000024511.12 | 0.595012147 | 1.04E-13 |
| Rnaset2b | ENSMUSG00000094724.4 | 0.85925806 | 1.55E-07 |
| Slc29a1 | ENSMUSG00000023942.12 | 0.622051 | 4.47E-04 |
| Sspn | ENSMUSG00000030255.10 | 0.518189587 | 3.58E-06 |
| Syt15 | ENSMUSG00000041479.10 | 1.409765644 | 5.88E-47 |
| Tmem59l | ENSMUSG00000035964.7 | 0.538314991 | 5.75E-10 |

| | | | |
|---|---|---|---|
| Tuba4a | ENSMUSG00000026202.10 | 0.819390438 | 1.23E-14 |
| Vcp | ENSMUSG00000028452.7 | 0.876030174 | 3.18E-29 |
| Vil1 | ENSMUSG00000026175.9 | 0.901565397 | 2.35E-08 |
| Zfp738 | ENSMUSG00000048280.14 | 0.853811659 | 6.58E-30 |

**Differentially expressed endothelial cell (EC) genes in the P10 Apcdd1-/- retina**

**Significantly downregulated genes**

| Gene_name | Ensembl_ID | log2FoldChange | padj |
|---|---|---|---|
| Acot1 | ENSMUSG00000072949.5 | -0.599754484 | 1.69E-05 |
| Alox5ap | ENSMUSG00000060063.6 | -1.26184423 | 3.28E-17 |
| Apcdd1 | ENSMUSG00000071847.9 | -0.837383081 | 6.75E-31 |
| Aurka | ENSMUSG00000027496.12 | -1.411782715 | 2.31E-35 |
| Calcrl | ENSMUSG00000059588.10 | -0.567152965 | 3.14E-05 |
| Ccdc171 | ENSMUSG00000052407.13 | -1.118809169 | 1.83E-20 |
| Ccdc24 | ENSMUSG00000078588.7 | -0.658904859 | 9.34E-05 |
| Col20a1 | ENSMUSG00000016356.14 | -0.681575818 | 5.80E-05 |
| Cyp4f13 | ENSMUSG00000024055.11 | -1.551764604 | 8.76E-50 |
| Dctn3 | ENSMUSG00000028447.8 | -0.503775922 | 6.96E-05 |
| Dxo | ENSMUSG00000040482.12 | -0.576322828 | 8.40E-15 |
| Fam210b | ENSMUSG00000027495.4 | -0.913504807 | 1.15E-40 |
| Fndc1 | ENSMUSG00000071984.7 | -0.924101471 | 1.47E-19 |
| Gdpd3 | ENSMUSG00000030703.7 | -0.557088638 | 0.002187315 |
| Gem | ENSMUSG00000028214.10 | -0.917873857 | 1.21E-19 |
| Glb1l | ENSMUSG00000026200.10 | -0.52057035 | 3.21E-06 |
| Gpc2 | ENSMUSG00000029510.12 | -0.53040727 | 0.00318171 |
| Gstp1 | ENSMUSG00000060803.5 | -0.509636682 | 2.59E-08 |
| H2-DMa | ENSMUSG00000037649.9 | -0.688043181 | 6.73E-09 |
| Hexdc | ENSMUSG00000039307.13 | -0.614404354 | 1.58E-14 |
| Hist1h2al | ENSMUSG00000091383.1 | -0.86919983 | 1.15E-08 |
| Itgb3bp | ENSMUSG00000028549.14 | -0.734683823 | 1.89E-10 |
| Lgals4 | ENSMUSG00000053964.13 | -0.803315253 | 4.20E-08 |
| Malt1 | ENSMUSG00000032688.7 | -0.712375007 | 1.17E-26 |
| Mob3b | ENSMUSG00000073910.7 | -0.560039248 | 3.61E-07 |
| Myo1d | ENSMUSG00000035441.11 | -0.563983532 | 8.91E-11 |
| Nos3 | ENSMUSG00000028978.9 | -0.796499121 | 1.44E-06 |
| Olfml2a | ENSMUSG00000046618.7 | -0.558525393 | 0.002024602 |
| Pgap2 | ENSMUSG00000030990.14 | -0.941898609 | 2.78E-17 |
| Plekhn1 | ENSMUSG00000078485.2 | -0.64394736 | 5.54E-08 |
| Polr2k | ENSMUSG00000045996.9 | -1.610521968 | 1.71E-49 |
| Prcp | ENSMUSG00000061119.6 | -0.529616729 | 3.18E-06 |
| Ring1 | ENSMUSG00000024325.8 | -0.676803013 | 7.62E-14 |
| Rreb1 | ENSMUSG00000039087.13 | -0.718518234 | 1.93E-20 |
| Rsph9 | ENSMUSG00000023966.5 | -0.548567717 | 3.36E-04 |
| Samd15 | ENSMUSG00000090812.5 | -0.763320161 | 1.17E-13 |
| Samhd1 | ENSMUSG00000027639.13 | -0.536330446 | 3.35E-09 |
| Spef2 | ENSMUSG00000072663.8 | -0.505044659 | 9.89E-04 |
| Stk19 | ENSMUSG00000061207.8 | -0.816553797 | 3.61E-18 |
| Stx3 | ENSMUSG00000041488.12 | -0.621444848 | 5.03E-17 |

| | | | |
|---|---|---|---|
| Tpm3-rs7 | ENSMUSG00000058126.6 | -3.570676155 | 2.29E-236 |
| Trim68 | ENSMUSG00000073968.3 | -0.968771797 | 9.27E-17 |
| Ulk4 | ENSMUSG00000040936.11 | -0.822923588 | 2.74E-18 |
| Upp2 | ENSMUSG00000026839.13 | -0.892776198 | 8.24E-12 |
| Arhgef39 | ENSMUSG00000051517.11 | -0.59500779 | 8.73E-04 |
| Efhc1 | ENSMUSG00000041809.5 | -0.642002398 | 1.24E-04 |
| Sh2d2a | ENSMUSG00000028071.9 | -1.202344098 | 7.31E-17 |
| Trim12c | ENSMUSG00000057143.11 | -1.440058456 | 6.77E-26 |

**Differentially expressed endothelial cell (EC) genes in the P14 Apcdd1-/- retina**

**Significantly upregulated genes**

| Gene_name | Ensembl_ID | log2FoldChange | padj |
|---|---|---|---|
| Arhgap8 | ENSMUSG00000078954 | 7.382382014 | 7.41E-78 |
| Rps3a3 | ENSMUSG00000059751 | 7.352887321 | 6.01E-201 |
| Rps3a2 | ENSMUSG00000062611 | 4.100287793 | 7.03E-44 |
| Rnaset2b | ENSMUSG00000094724 | 4.066275029 | 8.84E-74 |
| Cd1d2 | ENSMUSG00000041750 | 3.690347587 | 3.35E-11 |
| Lbp | ENSMUSG00000016024 | 3.66432372 | 9.50E-86 |
| Cwc22 | ENSMUSG00000027014 | 3.609834916 | 4.13E-24 |
| Aspg | ENSMUSG00000037686 | 3.581579065 | 2.38E-08 |
| Wnt10a | ENSMUSG00000026167 | 2.360052333 | 3.88E-05 |
| Glycam1 | ENSMUSG00000022491 | 2.165587869 | 0.00376477 |
| H2-D1 | ENSMUSG00000073411 | 2.090209147 | 1.85E-98 |
| Hs3st3a1 | ENSMUSG00000047759 | 1.890463021 | 8.29E-18 |
| Tinagl1 | ENSMUSG00000028776 | 1.794251824 | 1.48E-10 |
| Gstm2 | ENSMUSG00000040562 | 1.789107874 | 0.00303107 |
| Nos3 | ENSMUSG00000028978 | 1.787594364 | 2.17E-06 |
| Trim5 | ENSMUSG00000060441 | 1.729971267 | 2.69E-06 |
| Fosb | ENSMUSG00000003545 | 1.70050788 | 0.02595726 |
| Cfi | ENSMUSG00000058952 | 1.696312259 | 0.0003064 |
| Il11ra1 | ENSMUSG00000073889 | 1.670649256 | 8.13E-52 |
| Vil1 | ENSMUSG00000026175 | 1.598407979 | 5.61E-12 |
| Ctla2b | ENSMUSG00000074874 | 1.593284887 | 0.00294843 |
| H2-Q4 | ENSMUSG00000035929 | 1.552829397 | 4.48E-42 |
| Npb | ENSMUSG00000044034 | 1.551665791 | 3.43E-07 |
| Serpinb6b | ENSMUSG00000042842 | 1.528406514 | 4.50E-07 |
| Adh1 | ENSMUSG00000074207 | 1.515519746 | 0.00396103 |
| Perp | ENSMUSG00000019851 | 1.515513357 | 1.85E-26 |
| Slc26a11 | ENSMUSG00000039908 | 1.498214654 | 1.34E-43 |
| AI467606 | ENSMUSG00000045165 | 1.496868568 | 3.21E-10 |
| Kcne1 | ENSMUSG00000039639 | 1.494998998 | 0.00335182 |
| Dio3 | ENSMUSG00000075707 | 1.48746969 | 2.61E-09 |
| Tfpi | ENSMUSG00000027082 | 1.484475813 | 9.95E-14 |
| Optc | ENSMUSG00000010311 | 1.451802056 | 0.00459598 |
| Cyp2c39 | ENSMUSG00000025003 | 1.439439876 | 0.01016389 |
| Abi3bp | ENSMUSG00000035258 | 1.4084498 | 2.72E-06 |
| Fbln1 | ENSMUSG00000006369 | 1.382214811 | 0.00788349 |
| Hsd17b2 | ENSMUSG00000031844 | 1.380397299 | 6.13E-10 |
| Syt15 | ENSMUSG00000041479 | 1.374086256 | 2.57E-30 |
| Pla2g4a | ENSMUSG00000056220 | 1.334802516 | 1.81E-07 |
| Orm2 | ENSMUSG00000061540 | 1.329458148 | 1.43E-06 |
| Fos | ENSMUSG00000021250 | 1.30794222 | 0.0354153 |

| | | | |
|---|---|---|---|
| S100a6 | ENSMUSG00000001025 | 1.288018029 | 7.85E-12 |
| Frem2 | ENSMUSG00000037016 | 1.279804121 | 1.59E-07 |
| Fbln2 | ENSMUSG00000064080 | 1.269181266 | 4.85E-09 |
| Ndc80 | ENSMUSG00000024056 | 1.257749786 | 0.03031214 |
| Bcl3 | ENSMUSG00000053175 | 1.249762073 | 0.00302196 |
| Kifc1 | ENSMUSG00000079553 | 1.247358144 | 4.48E-14 |
| Zic4 | ENSMUSG00000036972 | 1.235011777 | 4.36E-05 |
| Ifitm1 | ENSMUSG00000025491 | 1.229138913 | 0.00504677 |
| Vcan | ENSMUSG00000021614 | 1.222609123 | 0.00673527 |
| Serping1 | ENSMUSG00000023224 | 1.220599464 | 4.85E-09 |
| Gstm1 | ENSMUSG00000058135 | 1.218773352 | 5.65E-09 |
| Stac | ENSMUSG00000032502 | 1.208317116 | 3.35E-07 |
| Hspb2 | ENSMUSG00000038086 | 1.187442123 | 0.00334527 |
| Colec12 | ENSMUSG00000036103 | 1.176767216 | 7.62E-08 |
| Klk8 | ENSMUSG00000064023 | 1.172902887 | 0.00089172 |
| Lrpprc | ENSMUSG00000024120 | 1.16849876 | 3.48E-65 |
| Gkn3 | ENSMUSG00000030048 | 1.166167435 | 0.02898672 |
| Col6a3 | ENSMUSG00000048126 | 1.165729253 | 0.00155246 |
| Pbk | ENSMUSG00000022033 | 1.160164893 | 0.01447432 |
| Ltc4s | ENSMUSG00000020377 | 1.155019326 | 0.00703393 |
| Gpr137b | ENSMUSG00000021306 | 1.128420356 | 2.64E-24 |
| Serpine1 | ENSMUSG00000037411 | 1.124201947 | 1.73E-07 |
| Plau | ENSMUSG00000021822 | 1.112059483 | 0.00834209 |
| Islr | ENSMUSG00000037206 | 1.111026998 | 0.00801283 |
| Egfl8 | ENSMUSG00000015467 | 1.110587307 | 7.79E-09 |
| Cd300a | ENSMUSG00000034652 | 1.110200568 | 0.00403279 |
| Scd4 | ENSMUSG00000050195 | 1.106568611 | 0.03517101 |
| Slc16a12 | ENSMUSG00000009378 | 1.100564726 | 3.74E-05 |
| Cpsf4l | ENSMUSG00000018727 | 1.094196758 | 0.00210323 |
| Ccna2 | ENSMUSG00000027715 | 1.087289009 | 3.26E-12 |
| Penk | ENSMUSG00000045573 | 1.083956414 | 2.09E-32 |
| H2-K1 | ENSMUSG00000061232 | 1.081186286 | 1.32E-18 |
| Serpinf1 | ENSMUSG00000000753 | 1.067388633 | 1.65E-10 |
| Eif4ebp1 | ENSMUSG00000031490 | 1.066726141 | 3.37E-08 |
| Dtl | ENSMUSG00000037474 | 1.066053113 | 0.01548692 |
| Nudt19 | ENSMUSG00000034875 | 1.059135142 | 3.77E-43 |
| Hrct1 | ENSMUSG00000071001 | 1.057520905 | 0.00186559 |
| Pglyrp1 | ENSMUSG00000030413 | 1.049464701 | 7.00E-06 |
| Ogn | ENSMUSG00000021390 | 1.047494775 | 2.03E-05 |
| Mt3 | ENSMUSG00000031760 | 1.044478444 | 2.70E-17 |
| Wfdc1 | ENSMUSG00000023336 | 1.042895268 | 5.75E-09 |
| Tnxb | ENSMUSG00000033327 | 1.042242071 | 2.26E-05 |
| Pcolce | ENSMUSG00000029718 | 1.038902907 | 5.17E-08 |
| Cybrd1 | ENSMUSG00000027015 | 1.036211782 | 0.00377074 |
| Tmprss11e | ENSMUSG00000054537 | 1.035805779 | 0.00296115 |

| | | | |
|---|---|---|---|
| Fzd1 | ENSMUSG00000044674 | 1.033822772 | 4.64E-14 |
| Rpph1 | ENSMUSG00000092837 | 1.02539214 | 0.00011072 |
| Zfp185 | ENSMUSG00000031351 | 1.02155113 | 2.71E-05 |
| Lix1 | ENSMUSG00000047786 | 1.021479733 | 9.74E-09 |
| Col10a1 | ENSMUSG00000039462 | 1.019678078 | 0.00447051 |
| Htra3 | ENSMUSG00000029096 | 1.018546679 | 3.55E-08 |
| S100a4 | ENSMUSG00000001020 | 1.016141116 | 0.00157684 |
| Zfp738 | ENSMUSG00000048280 | 1.001088578 | 1.64E-19 |
| Hspg2 | ENSMUSG00000028763 | 0.991144861 | 0.02626044 |
| Apbb1ip | ENSMUSG00000026786 | 0.988410881 | 0.03571352 |
| Igf2bp1 | ENSMUSG00000013415 | 0.977306549 | 0.00838845 |
| Mgst1 | ENSMUSG00000008540 | 0.959356606 | 3.44E-09 |
| Creb3l4 | ENSMUSG00000027938 | 0.947938211 | 0.00740055 |
| Ifitm2 | ENSMUSG00000060591 | 0.946441783 | 3.76E-11 |
| Gypc | ENSMUSG00000090523 | 0.939084001 | 0.00338326 |
| Nid1 | ENSMUSG00000005397 | 0.931835878 | 0.00610575 |
| Hpgds | ENSMUSG00000029919 | 0.931074286 | 0.01135332 |
| Cdc37l1 | ENSMUSG00000024780 | 0.928987836 | 5.59E-35 |
| Pon3 | ENSMUSG00000029759 | 0.928811902 | 2.02E-05 |
| Slc16a10 | ENSMUSG00000019838 | 0.928419879 | 3.17E-05 |
| Cyp4f16 | ENSMUSG00000048440 | 0.9225911 | 1.62E-06 |
| Slc5a5 | ENSMUSG00000000792 | 0.922457361 | 3.32E-10 |
| Mdfi | ENSMUSG00000032717 | 0.921753899 | 0.000209 |
| Fstl1 | ENSMUSG00000022816 | 0.921403692 | 6.23E-12 |
| Rpl35a | ENSMUSG00000060636 | 0.918976055 | 2.28E-19 |
| Anxa5 | ENSMUSG00000027712 | 0.918951093 | 9.78E-19 |
| Rnase4 | ENSMUSG00000021876 | 0.909971145 | 0.0035933 |
| Iqcg | ENSMUSG00000035578 | 0.905265677 | 1.03E-19 |
| Angptl2 | ENSMUSG00000004105 | 0.904556735 | 0.00012213 |
| Cyp2j9 | ENSMUSG00000015224 | 0.904282761 | 0.00034649 |
| Gja1 | ENSMUSG00000050953 | 0.901632998 | 0.01521701 |
| Aldh1a7 | ENSMUSG00000024747 | 0.891878054 | 3.72E-13 |
| Nr4a3 | ENSMUSG00000028341 | 0.887441293 | 0.00341143 |
| Rac2 | ENSMUSG00000033220 | 0.884789389 | 0.0447272 |
| Calml4 | ENSMUSG00000032246 | 0.875545226 | 6.12E-09 |
| Itga6 | ENSMUSG00000027111 | 0.874797958 | 1.43E-09 |
| Ccdc80 | ENSMUSG00000022665 | 0.872369723 | 5.02E-05 |
| Spon2 | ENSMUSG00000037379 | 0.871073605 | 1.75E-10 |
| Smagp | ENSMUSG00000053559 | 0.867269392 | 0.01443947 |
| Fcer1g | ENSMUSG00000058715 | 0.865797897 | 0.03970787 |
| Emp2 | ENSMUSG00000022505 | 0.86273157 | 6.51E-11 |
| Tmem37 | ENSMUSG00000050777 | 0.862412199 | 3.40E-05 |
| Ctgf | ENSMUSG00000019997 | 0.861809412 | 6.81E-07 |
| Slc15a2 | ENSMUSG00000022899 | 0.860240247 | 6.17E-34 |
| Zic1 | ENSMUSG00000032368 | 0.852381046 | 0.0001068 |

| | | | |
|---|---|---|---|
| Mtap | ENSMUSG00000062937 | 0.85216434 | 3.44E-07 |
| Sspn | ENSMUSG00000030255 | 0.850147071 | 6.08E-07 |
| Dtd1 | ENSMUSG00000027430 | 0.846058865 | 6.59E-14 |
| Rarres2 | ENSMUSG00000009281 | 0.843843295 | 6.75E-05 |
| Bche | ENSMUSG00000027792 | 0.843789906 | 1.55E-09 |
| Enpp1 | ENSMUSG00000037370 | 0.842126522 | 0.00013925 |
| Txnip | ENSMUSG00000038393 | 0.841887213 | 0.0001187 |
| Hist1h2bc | ENSMUSG00000018102 | 0.841745908 | 9.94E-09 |
| Ltbp1 | ENSMUSG00000001870 | 0.837608829 | 0.00043299 |
| Plekhf1 | ENSMUSG00000074170 | 0.82859868 | 4.08E-05 |
| Psmc3ip | ENSMUSG00000019303 | 0.824511175 | 0.0002005 |
| Ndufa4l2 | ENSMUSG00000040280 | 0.822001567 | 3.38E-05 |
| Atp1a2 | ENSMUSG00000007097 | 0.820246224 | 1.13E-08 |
| Paqr6 | ENSMUSG00000041423 | 0.818377537 | 0.00277284 |
| Zfp429 | ENSMUSG00000078994 | 0.817660767 | 0.00786133 |
| Ccnd2 | ENSMUSG00000000184 | 0.817209782 | 9.45E-05 |
| Rhod | ENSMUSG00000041845 | 0.817197662 | 1.31E-07 |
| Fcgr3 | ENSMUSG00000059498 | 0.814258259 | 0.02231543 |
| Rassf3 | ENSMUSG00000025795 | 0.809144703 | 5.25E-09 |
| Col18a1 | ENSMUSG00000001435 | 0.805157116 | 0.0404106 |
| Fbxl8 | ENSMUSG00000033313 | 0.800534482 | 0.00883114 |
| Pdk1 | ENSMUSG00000006494 | 0.79908467 | 3.10E-24 |
| Nupr1 | ENSMUSG00000030717 | 0.798888859 | 0.00013923 |
| Fancg | ENSMUSG00000028453 | 0.798175135 | 2.84E-07 |
| Mki67 | ENSMUSG00000031004 | 0.798122757 | 0.01128756 |
| Igfbp7 | ENSMUSG00000036256 | 0.792993356 | 4.05E-05 |
| Cd44 | ENSMUSG00000005087 | 0.789963393 | 1.08E-05 |
| Antxr1 | ENSMUSG00000033420 | 0.780545812 | 4.67E-09 |
| Elf4 | ENSMUSG00000031103 | 0.775297859 | 0.04406086 |
| Ass1 | ENSMUSG00000076441 | 0.771089837 | 4.51E-19 |
| Tnfrsf10b | ENSMUSG00000022074 | 0.762717692 | 0.00133978 |
| Col4a5 | ENSMUSG00000031274 | 0.759786081 | 0.00018587 |
| Psmb9 | ENSMUSG00000096727 | 0.759530414 | 0.01497576 |
| Ifitm3 | ENSMUSG00000025492 | 0.758566168 | 3.84E-06 |
| Slc29a1 | ENSMUSG00000023942 | 0.756884661 | 0.01853492 |
| Tspo | ENSMUSG00000041736 | 0.750605322 | 1.70E-07 |
| Cenpf | ENSMUSG00000026605 | 0.74756641 | 0.01869037 |
| Clec18a | ENSMUSG00000033633 | 0.745123856 | 0.0005277 |
| Nkd2 | ENSMUSG00000021567 | 0.742514976 | 0.0003696 |
| Bmp4 | ENSMUSG00000021835 | 0.739085424 | 0.00065171 |
| Metap1d | ENSMUSG00000041921 | 0.736408035 | 1.61E-07 |
| Ccnb1 | ENSMUSG00000041431 | 0.73518044 | 0.03373068 |
| Tk1 | ENSMUSG00000025574 | 0.734829134 | 0.01490851 |
| Rny3 | ENSMUSG00000064945 | 0.733877377 | 0.04018942 |
| Gprc5c | ENSMUSG00000051043 | 0.732522334 | 0.00212863 |

| | | | |
|---|---|---|---|
| Rps2-ps10 | ENSMUSG00000091957 | 0.731131602 | 0.000471 |
| Krt18 | ENSMUSG00000023043 | 0.729153852 | 2.70E-11 |
| Cebpb | ENSMUSG00000056501 | 0.724255096 | 0.00386757 |
| Tmem59l | ENSMUSG00000035964 | 0.719335521 | 1.80E-12 |
| Slc25a45 | ENSMUSG00000024818 | 0.716710543 | 0.02842455 |
| Lpar2 | ENSMUSG00000031861 | 0.713346058 | 0.00099848 |
| Gpc4 | ENSMUSG00000031119 | 0.711313638 | 1.54E-06 |
| Cp | ENSMUSG00000003617 | 0.708888869 | 3.86E-08 |
| Lrrc55 | ENSMUSG00000075224 | 0.70729457 | 0.00476568 |
| Myl9 | ENSMUSG00000067818 | 0.705971152 | 8.49E-12 |
| Rn7sk | ENSMUSG00000065037 | 0.704412707 | 0.00389739 |
| Hexb | ENSMUSG00000021665 | 0.703611509 | 0.00013925 |
| Mecom | ENSMUSG00000027684 | 0.702202731 | 0.00547585 |
| Kitl | ENSMUSG00000019966 | 0.701416132 | 1.05E-15 |
| Frk | ENSMUSG00000019779 | 0.701322807 | 0.02386451 |
| Nme1 | ENSMUSG00000037601 | 0.700971446 | 4.71E-13 |
| Fam60a | ENSMUSG00000039985 | 0.700824501 | 3.67E-05 |
| Cyba | ENSMUSG00000006519 | 0.696929368 | 1.70E-05 |
| Kdelr3 | ENSMUSG00000010830 | 0.695671608 | 0.004485 |
| Bace2 | ENSMUSG00000040605 | 0.694630768 | 0.0001508 |
| Ctsh | ENSMUSG00000032359 | 0.694126793 | 1.35E-07 |
| Aldh1a1 | ENSMUSG00000053279 | 0.691998349 | 1.75E-06 |
| Gpx8 | ENSMUSG00000021760 | 0.691421425 | 1.06E-14 |
| Hhex | ENSMUSG00000024986 | 0.689495158 | 0.02739009 |
| Selenop | ENSMUSG00000064373 | 0.68869521 | 6.97E-06 |
| Agmo | ENSMUSG00000050103 | 0.68752633 | 0.04928382 |
| Vcp | ENSMUSG00000028452 | 0.687173204 | 4.64E-24 |
| Dse | ENSMUSG00000039497 | 0.684687083 | 0.0008346 |
| Efemp1 | ENSMUSG00000020467 | 0.682933524 | 1.98E-10 |
| S100a10 | ENSMUSG00000041959 | 0.68257472 | 5.82E-09 |
| Fam107a | ENSMUSG00000021750 | 0.682192083 | 0.03033048 |
| Cpxm1 | ENSMUSG00000027408 | 0.677600783 | 6.55E-08 |
| Abca9 | ENSMUSG00000041797 | 0.673314256 | 0.00305537 |
| Stbd1 | ENSMUSG00000047963 | 0.673154294 | 1.30E-05 |
| Nexn | ENSMUSG00000039103 | 0.67303593 | 0.04275988 |
| Cd38 | ENSMUSG00000029084 | 0.672723992 | 0.00582417 |
| Atp1b3 | ENSMUSG00000032412 | 0.670376039 | 3.75E-10 |
| Matn2 | ENSMUSG00000022324 | 0.668640936 | 0.00193311 |
| Ahnak | ENSMUSG00000069833 | 0.667311148 | 0.00172803 |
| S100a11 | ENSMUSG00000027907 | 0.666569361 | 0.00081494 |
| Tal1 | ENSMUSG00000028717 | 0.665334461 | 0.03271533 |
| Crip1 | ENSMUSG00000006360 | 0.663030455 | 0.01127085 |
| Adm | ENSMUSG00000030790 | 0.661363993 | 0.01284728 |
| Klhl6 | ENSMUSG00000043008 | 0.661117263 | 0.01709747 |
| Pxdn | ENSMUSG00000020674 | 0.660743742 | 3.41E-06 |

| | | | |
|---|---|---|---|
| Scd1 | ENSMUSG00000037071 | 0.659453835 | 7.65E-08 |
| Emp3 | ENSMUSG00000040212 | 0.655145496 | 0.00931323 |
| Rpl22l1 | ENSMUSG00000039221 | 0.650451604 | 4.31E-07 |
| Fgfr2 | ENSMUSG00000030849 | 0.644932484 | 0.00478497 |
| Rcn3 | ENSMUSG00000019539 | 0.644219313 | 7.98E-06 |
| Ramp1 | ENSMUSG00000034353 | 0.639035946 | 0.0002462 |
| Glb1 | ENSMUSG00000045594 | 0.637938811 | 1.19E-09 |
| Hist1h4i | ENSMUSG00000060639 | 0.634084522 | 0.00754005 |
| Il1r1 | ENSMUSG00000026072 | 0.6334096 | 4.40E-05 |
| Tec | ENSMUSG00000029217 | 0.633350154 | 0.01065947 |
| Hmcn1 | ENSMUSG00000066842 | 0.633256495 | 0.02942465 |
| Zcchc9 | ENSMUSG00000021621 | 0.632953226 | 2.35E-12 |
| Tcp11l1 | ENSMUSG00000027175 | 0.632854828 | 0.00316604 |
| Mfap2 | ENSMUSG00000060572 | 0.63265134 | 2.16E-05 |
| Colgalt2 | ENSMUSG00000032649 | 0.630489529 | 1.14E-10 |
| Ackr3 | ENSMUSG00000044337 | 0.630385513 | 4.88E-05 |
| Col9a1 | ENSMUSG00000026147 | 0.629593123 | 7.79E-07 |
| Fxyd6 | ENSMUSG00000066705 | 0.629202382 | 8.84E-05 |
| Hspe1 | ENSMUSG00000073676 | 0.628098425 | 3.07E-07 |
| Mmp2 | ENSMUSG00000031740 | 0.627884766 | 5.10E-06 |
| Ech1 | ENSMUSG00000053898 | 0.62665877 | 1.54E-14 |
| Smco4 | ENSMUSG00000058173 | 0.623694266 | 0.01154656 |
| Peg10 | ENSMUSG00000092035 | 0.623013619 | 0.00397014 |
| Zfp605 | ENSMUSG00000023284 | 0.622729561 | 7.03E-05 |
| Slc8b1 | ENSMUSG00000032754 | 0.620628192 | 3.31E-05 |
| Tnfaip8 | ENSMUSG00000062210 | 0.619110067 | 4.28E-10 |
| Itpr3 | ENSMUSG00000042644 | 0.61858566 | 0.0174185 |
| Ctsc | ENSMUSG00000030560 | 0.617365811 | 1.74E-08 |
| Sgsh | ENSMUSG00000005043 | 0.617302765 | 2.44E-07 |
| Tnfrsf12a | ENSMUSG00000023905 | 0.614999667 | 0.00097249 |
| Mlc1 | ENSMUSG00000035805 | 0.614044318 | 1.18E-14 |
| Hebp2 | ENSMUSG00000019853 | 0.613551677 | 1.41E-05 |
| Pign | ENSMUSG00000056536 | 0.613465901 | 2.29E-06 |
| Slc6a13 | ENSMUSG00000030108 | 0.612542905 | 0.00025929 |
| Rrm2 | ENSMUSG00000020649 | 0.610398915 | 0.0262763 |
| Tuba4a | ENSMUSG00000026202 | 0.610048055 | 0.04878346 |
| Tmbim1 | ENSMUSG00000006301 | 0.607409978 | 4.22E-09 |
| Ptgs1 | ENSMUSG00000047250 | 0.601733294 | 0.040691 |
| Ppic | ENSMUSG00000024538 | 0.601602874 | 6.70E-06 |
| S1pr3 | ENSMUSG00000067586 | 0.600515479 | 6.13E-05 |
| Kcnip4 | ENSMUSG00000029088 | 0.599714577 | 3.31E-16 |
| Igf2 | ENSMUSG00000048583 | 0.598124804 | 3.30E-17 |
| Arhgdib | ENSMUSG00000030220 | 0.596405679 | 0.00076883 |
| Traf3ip3 | ENSMUSG00000037318 | 0.594348455 | 0.04342472 |
| Csf1 | ENSMUSG00000014599 | 0.591530538 | 0.00459021 |

| | | | |
|---|---|---|---|
| Ctsk | ENSMUSG00000028111 | 0.589596718 | 0.04154892 |
| Arpc1b | ENSMUSG00000029622 | 0.589486804 | 1.01E-06 |
| Sdc1 | ENSMUSG00000020592 | 0.586488032 | 0.02211185 |
| Mrc2 | ENSMUSG00000020695 | 0.585414443 | 0.00307837 |
| Apoe | ENSMUSG00000002985 | 0.584954013 | 8.91E-11 |
| Hfe | ENSMUSG00000006611 | 0.584042062 | 3.75E-09 |
| Iqgap2 | ENSMUSG00000021676 | 0.582390192 | 0.00552437 |
| Olfml2a | ENSMUSG00000046618 | 0.581403347 | 0.04022057 |
| Fign | ENSMUSG00000075324 | 0.581331339 | 0.01173101 |
| Ecm1 | ENSMUSG00000028108 | 0.580854177 | 0.0005673 |
| Itih5 | ENSMUSG00000025780 | 0.580188802 | 0.00051732 |
| Irak1bp1 | ENSMUSG00000032251 | 0.578062185 | 0.00297862 |
| Tgif1 | ENSMUSG00000047407 | 0.577446318 | 0.00316987 |
| Pomc | ENSMUSG00000020660 | 0.577220493 | 0.00048614 |
| Zfp36 | ENSMUSG00000044786 | 0.570392991 | 0.00100244 |
| Cst3 | ENSMUSG00000027447 | 0.56866207 | 1.95E-08 |
| Moxd1 | ENSMUSG00000020000 | 0.56861417 | 2.45E-07 |
| Rwdd3 | ENSMUSG00000028133 | 0.568262512 | 4.43E-05 |
| Arhgap25 | ENSMUSG00000030047 | 0.568110115 | 0.04662058 |
| Rab27b | ENSMUSG00000024511 | 0.567343773 | 1.13E-12 |
| Igfbp2 | ENSMUSG00000039323 | 0.565420238 | 6.10E-06 |
| Zfp758 | ENSMUSG00000044501 | 0.564847382 | 0.00052589 |
| Cmtm8 | ENSMUSG00000041012 | 0.564191223 | 0.00043585 |
| Tnfrsf1b | ENSMUSG00000028599 | 0.562224688 | 0.03163298 |
| Fam114a1 | ENSMUSG00000029185 | 0.561421077 | 0.00083438 |
| Col9a2 | ENSMUSG00000028626 | 0.558834875 | 2.32E-05 |
| Nhsl1 | ENSMUSG00000039835 | 0.55861205 | 0.0002128 |
| Ifi30 | ENSMUSG00000031838 | 0.558556317 | 5.86E-05 |
| Gas5 | ENSMUSG00000053332 | 0.557821064 | 8.24E-06 |
| Bambi | ENSMUSG00000024232 | 0.556284457 | 5.04E-07 |
| Cgn | ENSMUSG00000068876 | 0.555910899 | 0.0001018 |
| Itm2a | ENSMUSG00000031239 | 0.555554936 | 2.49E-09 |
| Il10rb | ENSMUSG00000022969 | 0.554585863 | 0.00083842 |
| Lyn | ENSMUSG00000042228 | 0.551013622 | 0.00372338 |
| Gng11 | ENSMUSG00000032766 | 0.546729958 | 0.0007345 |
| Bhlhe40 | ENSMUSG00000030103 | 0.544757488 | 5.03E-07 |
| Cep250 | ENSMUSG00000038241 | 0.543653654 | 5.04E-09 |
| Icam1 | ENSMUSG00000037405 | 0.542302439 | 0.04059287 |
| Crim1 | ENSMUSG00000024074 | 0.540888707 | 0.00030928 |
| Sri | ENSMUSG00000003161 | 0.539828322 | 1.73E-13 |
| Herc6 | ENSMUSG00000029798 | 0.538353623 | 0.03254669 |
| Six2 | ENSMUSG00000024134 | 0.537989701 | 0.03093853 |
| Mrps6 | ENSMUSG00000039680 | 0.536482086 | 2.96E-06 |
| Arsk | ENSMUSG00000021592 | 0.535215243 | 0.00014244 |
| Sparc | ENSMUSG00000018593 | 0.534698888 | 2.70E-21 |

| | | | |
|---|---|---|---|
| Golm1 | ENSMUSG00000021556 | 0.533589325 | 0.00732682 |
| Cnbd2 | ENSMUSG00000038085 | 0.532103702 | 1.64E-07 |
| Thbs2 | ENSMUSG00000023885 | 0.530587028 | 0.01705699 |
| Cd320 | ENSMUSG00000002308 | 0.529756015 | 8.03E-07 |
| Gusb | ENSMUSG00000025534 | 0.528349244 | 0.00012925 |
| Top2a | ENSMUSG00000020914 | 0.52800728 | 0.03019222 |
| Zfp697 | ENSMUSG00000050064 | 0.524334735 | 0.00223375 |
| Tagln2 | ENSMUSG00000026547 | 0.524144417 | 0.00486834 |
| Fkbp9 | ENSMUSG00000029781 | 0.522430293 | 8.57E-07 |
| Zic3 | ENSMUSG00000067860 | 0.518548366 | 0.03190567 |
| Pmp22 | ENSMUSG00000018217 | 0.518310451 | 2.05E-08 |
| Cd63 | ENSMUSG00000025351 | 0.517079516 | 9.67E-10 |
| Cgnl1 | ENSMUSG00000032232 | 0.51676679 | 0.00055365 |
| Fcgrt | ENSMUSG00000003420 | 0.516472919 | 0.00124559 |
| Wnt5b | ENSMUSG00000030170 | 0.513667319 | 0.00013723 |
| Marveld2 | ENSMUSG00000021636 | 0.51315534 | 0.00948718 |
| Plp2 | ENSMUSG00000031146 | 0.512758728 | 0.02421786 |
| Thbd | ENSMUSG00000074743 | 0.512581936 | 0.02810901 |
| Anxa2 | ENSMUSG00000032231 | 0.511944245 | 5.15E-07 |
| Nrros | ENSMUSG00000052384 | 0.511454803 | 0.00881317 |
| Hist1h2be | ENSMUSG00000047246 | 0.509442917 | 0.02409815 |
| Prelp | ENSMUSG00000041577 | 0.506744501 | 0.00071721 |
| Slc22a8 | ENSMUSG00000063796 | 0.504174693 | 0.00518332 |
| Efna1 | ENSMUSG00000027954 | 0.503281376 | 0.00013067 |
| Bub1b | ENSMUSG00000040084 | 0.50266146 | 0.00547585 |
| Mfsd2a | ENSMUSG00000028655 | 0.501647802 | 8.20E-05 |
| Suclg2 | ENSMUSG00000061838 | 0.501563008 | 0.00092556 |
| Ptpn14 | ENSMUSG00000026604 | 0.501187067 | 0.01918108 |
| Lipa | ENSMUSG00000024781 | 0.500909301 | 1.13E-08 |
| Hspb8 | ENSMUSG00000041548 | 0.500554422 | 0.00388364 |
| Stab1 | ENSMUSG00000042286 | 0.500200658 | 0.00898819 |

#### Differentially expressed endothelial cell (EC) genes in the P14 Apcdd1-/- retina

##### Significantly downregulated genes

| Gene_name | Ensembl_ID | log2FoldChange | padj |
|---|---|---|---|
| Tpm3-rs7 | ENSMUSG00000058126 | -5.885759703 | 8.49E-93 |
| Rpsa-ps10 | ENSMUSG00000047676 | -3.895177598 | 0.00548932 |
| Gdpd3 | ENSMUSG00000030703 | -3.704505962 | 2.39E-45 |
| Sh2d2a | ENSMUSG00000028071 | -3.680911951 | 0.00077388 |
| Pax5 | ENSMUSG00000014030 | -2.642748817 | 2.62E-11 |
| Alox5ap | ENSMUSG00000060063 | -2.168444026 | 1.84E-49 |
| Tmod4 | ENSMUSG00000005628 | -1.876385392 | 9.44E-09 |
| Bcan | ENSMUSG00000004892 | -1.846246177 | 1.64E-24 |
| Clic3 | ENSMUSG00000015093 | -1.801625099 | 6.34E-14 |
| Padi6 | ENSMUSG00000040935 | -1.526364755 | 0.00131541 |
| Ulk4 | ENSMUSG00000040936 | -1.512493436 | 6.90E-30 |
| Lgals4 | ENSMUSG00000053964 | -1.505215908 | 5.94E-20 |
| Efhc1 | ENSMUSG00000041809 | -1.499155219 | 4.47E-08 |
| Cyp4f13 | ENSMUSG00000024055 | -1.491463641 | 2.30E-26 |
| Gpr137b-ps | ENSMUSG00000097715 | -1.455830473 | 6.00E-24 |
| Hist1h2al | ENSMUSG00000091383 | -1.450278602 | 1.14E-47 |
| Polr2k | ENSMUSG00000045996 | -1.431334143 | 1.92E-35 |
| Rsph9 | ENSMUSG00000023966 | -1.417479732 | 6.24E-27 |
| Col20a1 | ENSMUSG00000016356 | -1.3512778 | 4.47E-30 |
| Fndc1 | ENSMUSG00000071984 | -1.320665706 | 9.39E-14 |
| Ccdc171 | ENSMUSG00000052407 | -1.253616039 | 1.24E-15 |
| Endou | ENSMUSG00000022468 | -1.209691799 | 0.00120927 |
| Trib1 | ENSMUSG00000032501 | -1.207504839 | 2.64E-17 |
| Stk19 | ENSMUSG00000061207 | -1.205132554 | 4.60E-38 |
| Htr3a | ENSMUSG00000032269 | -1.190080128 | 3.13E-14 |
| Xlr3a | ENSMUSG00000057836 | -1.189701148 | 0.00298479 |
| Ccdc24 | ENSMUSG00000078588 | -1.166297595 | 4.11E-36 |
| Aurka | ENSMUSG00000027496 | -1.145364461 | 4.55E-16 |
| Rreb1 | ENSMUSG00000039087 | -1.128640345 | 1.15E-12 |
| Trim12c | ENSMUSG00000057143 | -1.122692805 | 0.0001936 |
| Acr | ENSMUSG00000022622 | -1.118083297 | 0.02817838 |
| Gem | ENSMUSG00000028214 | -1.112609495 | 3.03E-39 |
| Rec8 | ENSMUSG00000002324 | -1.109584903 | 0.00623423 |
| Calcrl | ENSMUSG00000059588 | -1.06255826 | 2.70E-17 |
| U2af1l4 | ENSMUSG00000109378 | -1.052960924 | 1.47E-06 |
| Lama3 | ENSMUSG00000024421 | -1.044360535 | 1.33E-10 |
| Col11a2 | ENSMUSG00000024330 | -1.039438789 | 7.89E-20 |
| Zfp97 | ENSMUSG00000095990 | -1.02956866 | 1.16E-05 |
| Rnf207 | ENSMUSG00000058498 | -1.019467089 | 1.26E-16 |
| H2-Q10 | ENSMUSG00000067235 | -1.001866951 | 0.03051375 |

| | | | |
|---|---|---|---|
| Arhgef39 | ENSMUSG00000051517 | -1.000101395 | 0.00036361 |
| Wdr66 | ENSMUSG00000029442 | -0.995215137 | 1.40E-06 |
| Fam110a | ENSMUSG00000027459 | -0.987190883 | 3.62E-42 |
| Catsper4 | ENSMUSG00000048003 | -0.986859874 | 0.00053914 |
| Plvap | ENSMUSG00000034845 | -0.980562775 | 9.56E-11 |
| Samd15 | ENSMUSG00000090812 | -0.979549551 | 2.39E-07 |
| Sspo | ENSMUSG00000029797 | -0.961063415 | 0.00388867 |
| Acot1 | ENSMUSG00000072949 | -0.953252782 | 5.42E-12 |
| Samd11 | ENSMUSG00000096351 | -0.952704051 | 1.55E-19 |
| Ero1lb | ENSMUSG00000057069 | -0.944305596 | 2.77E-23 |
| Ptpmt1 | ENSMUSG00000063235 | -0.909807886 | 9.70E-13 |
| Nudt8 | ENSMUSG00000024869 | -0.903205204 | 5.64E-05 |
| Dctn3 | ENSMUSG00000028447 | -0.900143271 | 5.83E-33 |
| Ring1 | ENSMUSG00000024325 | -0.884910138 | 1.19E-39 |
| Mob3b | ENSMUSG00000073910 | -0.884347601 | 2.89E-07 |
| Ybx2 | ENSMUSG00000018554 | -0.87688882 | 1.30E-09 |
| Myo1d | ENSMUSG00000035441 | -0.857001795 | 1.07E-11 |
| Pde6b | ENSMUSG00000029491 | -0.853383039 | 7.22E-06 |
| Col27a1 | ENSMUSG00000045672 | -0.838194001 | 1.11E-13 |
| Nr1h3 | ENSMUSG00000002108 | -0.837490798 | 5.15E-05 |
| Pgap2 | ENSMUSG00000030990 | -0.831797989 | 1.05E-05 |
| Clstn1 | ENSMUSG00000039953 | -0.828336899 | 3.20E-55 |
| Plekhn1 | ENSMUSG00000078485 | -0.822372774 | 8.16E-07 |
| Col6a1 | ENSMUSG00000001119 | -0.816607391 | 3.13E-09 |
| Ly6g6d | ENSMUSG00000073413 | -0.805271352 | 0.02262851 |
| Dxo | ENSMUSG00000040482 | -0.797440171 | 1.90E-28 |
| Cldn7 | ENSMUSG00000018569 | -0.787710106 | 1.57E-06 |
| Gpsm3 | ENSMUSG00000034786 | -0.785766182 | 0.01983805 |
| H2-DMa | ENSMUSG00000037649 | -0.785759263 | 6.22E-08 |
| Rab37 | ENSMUSG00000020732 | -0.774202727 | 0.00073422 |
| Il3ra | ENSMUSG00000068758 | -0.773961298 | 2.68E-06 |
| Fam210b | ENSMUSG00000027495 | -0.758281931 | 1.54E-28 |
| Sdcbp2 | ENSMUSG00000027456 | -0.746911593 | 0.0114404 |
| Gpc2 | ENSMUSG00000029510 | -0.746820733 | 6.88E-05 |
| Apcdd1 | ENSMUSG00000071847 | -0.738717045 | 1.89E-26 |
| St6galnac2 | ENSMUSG00000110170 | -0.734578793 | 0.0330748 |
| Glb1l | ENSMUSG00000026200 | -0.727517265 | 2.61E-06 |
| Rtel1 | ENSMUSG00000038685 | -0.721849452 | 1.03E-13 |
| Ccdc163 | ENSMUSG00000028689 | -0.719833686 | 1.73E-10 |
| Ankrd33b | ENSMUSG00000022237 | -0.714170318 | 2.39E-10 |
| Zmynd15 | ENSMUSG00000040829 | -0.713925997 | 0.00066966 |
| Cacna1f | ENSMUSG00000031142 | -0.707063075 | 6.13E-15 |
| Malt1 | ENSMUSG00000032688 | -0.702638958 | 2.64E-06 |
| Prcp | ENSMUSG00000061119 | -0.701929877 | 2.40E-08 |
| Doc2g | ENSMUSG00000024871 | -0.698374832 | 0.00056676 |

| | | | |
|---|---|---|---|
| Trim68 | ENSMUSG00000073968 | -0.696488039 | 3.23E-05 |
| Gstp1 | ENSMUSG00000060803 | -0.694615541 | 1.18E-10 |
| Snx22 | ENSMUSG00000039452 | -0.691133707 | 0.03271533 |
| Igsf21 | ENSMUSG00000040972 | -0.689880743 | 7.17E-14 |
| Jmjd7 | ENSMUSG00000098789 | -0.687495831 | 0.0059278 |
| Tle2 | ENSMUSG00000034771 | -0.683937971 | 3.34E-07 |
| Itgb3bp | ENSMUSG00000028549 | -0.682226846 | 2.21E-05 |
| Saal1 | ENSMUSG00000006763 | -0.674916852 | 2.90E-14 |
| Impg2 | ENSMUSG00000035270 | -0.670192445 | 1.23E-11 |
| Rps3a1 | ENSMUSG00000028081 | -0.667251588 | 7.53E-19 |
| Xaf1 | ENSMUSG00000040483 | -0.665746786 | 0.03964004 |
| Zfp692 | ENSMUSG00000037243 | -0.664173457 | 3.52E-08 |
| Clasrp | ENSMUSG00000061028 | -0.657840539 | 1.10E-11 |
| Efcab2 | ENSMUSG00000026495 | -0.654178347 | 2.92E-07 |
| Gmip | ENSMUSG00000036246 | -0.651063754 | 8.41E-07 |
| Sfxn2 | ENSMUSG00000025036 | -0.648470949 | 4.40E-07 |
| Sema4c | ENSMUSG00000026121 | -0.64733993 | 1.29E-13 |
| Spata6 | ENSMUSG00000034401 | -0.640228846 | 8.58E-08 |
| Zan | ENSMUSG00000079173 | -0.63944338 | 1.57E-06 |
| Dhx35 | ENSMUSG00000027655 | -0.635867998 | 3.42E-21 |
| Rabgef1 | ENSMUSG00000025340 | -0.633994858 | 1.23E-08 |
| Pvr | ENSMUSG00000040511 | -0.628296715 | 3.14E-11 |
| Fam193b | ENSMUSG00000021495 | -0.625937385 | 1.12E-09 |
| Chtf18 | ENSMUSG00000019214 | -0.62573507 | 3.84E-06 |
| Asic3 | ENSMUSG00000038276 | -0.622350262 | 2.11E-09 |
| Khsrp | ENSMUSG00000007670 | -0.621681865 | 1.41E-20 |
| Gucy2e | ENSMUSG00000020890 | -0.618986938 | 7.23E-11 |
| Stom | ENSMUSG00000026880 | -0.618691769 | 1.40E-12 |
| Klhl33 | ENSMUSG00000090799 | -0.616480863 | 0.00018226 |
| Rhpn1 | ENSMUSG00000022580 | -0.615656037 | 7.79E-07 |
| Smug1 | ENSMUSG00000036061 | -0.614528512 | 6.01E-22 |
| Scrn2 | ENSMUSG00000020877 | -0.613114159 | 6.82E-06 |
| Spef2 | ENSMUSG00000072663 | -0.611432064 | 0.00732434 |
| Upp2 | ENSMUSG00000026839 | -0.61028347 | 0.00642985 |
| Sema3c | ENSMUSG00000028780 | -0.610022031 | 3.89E-08 |
| Kif21b | ENSMUSG00000041642 | -0.608850766 | 4.07E-05 |
| Morn1 | ENSMUSG00000029049 | -0.607917111 | 5.38E-09 |
| Tead3 | ENSMUSG00000002249 | -0.596589561 | 1.48E-07 |
| Dnase1l2 | ENSMUSG00000024136 | -0.595764123 | 4.41E-05 |
| L3hypdh | ENSMUSG00000019718 | -0.587035308 | 9.44E-09 |
| Hexdc | ENSMUSG00000039307 | -0.583501761 | 5.32E-12 |
| Crocc | ENSMUSG00000040860 | -0.583261321 | 4.52E-09 |
| Pdzrn3 | ENSMUSG00000035357 | -0.579041603 | 4.20E-05 |
| Iqcd | ENSMUSG00000029601 | -0.577548994 | 0.00086635 |
| Leng8 | ENSMUSG00000035545 | -0.577536215 | 4.00E-05 |

| | | | |
|---|---|---|---|
| Sh3pxd2b | ENSMUSG00000040711 | -0.577449901 | 0.0014947 |
| Kif19a | ENSMUSG00000010021 | -0.576966385 | 0.00014537 |
| Ubxn11 | ENSMUSG00000012126 | -0.576643743 | 2.67E-06 |
| Bcar3 | ENSMUSG00000028121 | -0.574629516 | 1.53E-06 |
| Abhd14b | ENSMUSG00000042073 | -0.574422867 | 1.29E-07 |
| Tmem150a | ENSMUSG00000055912 | -0.568757504 | 1.16E-06 |
| Ing4 | ENSMUSG00000030330 | -0.566432011 | 0.00899325 |
| Dyrk2 | ENSMUSG00000028630 | -0.56501104 | 0.00038939 |
| Prkab1 | ENSMUSG00000029513 | -0.562089571 | 1.47E-08 |
| Ppef2 | ENSMUSG00000029410 | -0.560839477 | 2.77E-10 |
| Plch2 | ENSMUSG00000029055 | -0.560097983 | 1.07E-13 |
| Rgs20 | ENSMUSG00000002459 | -0.559949328 | 0.00015448 |
| Kbtbd4 | ENSMUSG00000005505 | -0.559483928 | 2.23E-08 |
| Abca4 | ENSMUSG00000028125 | -0.557879456 | 9.82E-08 |
| Cables1 | ENSMUSG00000040957 | -0.557155806 | 3.57E-09 |
| Dynlrb2 | ENSMUSG00000034467 | -0.556740489 | 0.00995319 |
| Polm | ENSMUSG00000020474 | -0.556215371 | 1.64E-05 |
| Mamdc4 | ENSMUSG00000026941 | -0.554644906 | 0.00147336 |
| Stard9 | ENSMUSG00000033705 | -0.551564202 | 0.002862 |
| Samhd1 | ENSMUSG00000027639 | -0.549407362 | 1.41E-11 |
| Wisp1 | ENSMUSG00000005124 | -0.546254279 | 1.42E-05 |
| Rad54b | ENSMUSG00000078773 | -0.545872133 | 0.01839393 |
| Radil | ENSMUSG00000029576 | -0.543373417 | 2.09E-07 |
| Plcd3 | ENSMUSG00000020937 | -0.540997294 | 2.22E-09 |
| Hmgb2 | ENSMUSG00000054717 | -0.540097896 | 2.95E-06 |
| Rfx1 | ENSMUSG00000031706 | -0.539852113 | 1.01E-08 |
| Dnah9 | ENSMUSG00000056752 | -0.539617541 | 4.64E-05 |
| Slc25a19 | ENSMUSG00000020744 | -0.539525912 | 6.27E-08 |
| Fscn2 | ENSMUSG00000025380 | -0.539038929 | 0.00010467 |
| Cul9 | ENSMUSG00000040327 | -0.538543859 | 8.78E-18 |
| Gigyf1 | ENSMUSG00000029714 | -0.53751981 | 1.90E-06 |
| Pfkfb2 | ENSMUSG00000026409 | -0.536841023 | 1.00E-13 |
| Cpt1a | ENSMUSG00000024900 | -0.535706174 | 7.25E-07 |
| Stx3 | ENSMUSG00000041488 | -0.53489457 | 1.29E-16 |
| Amy1 | ENSMUSG00000074264 | -0.534804193 | 0.00225038 |
| Uckl1 | ENSMUSG00000089917 | -0.531808978 | 8.47E-07 |
| Slc9a5 | ENSMUSG00000014786 | -0.530942672 | 1.96E-09 |
| Mesp1 | ENSMUSG00000030544 | -0.530538883 | 0.03140554 |
| Csf2ra | ENSMUSG00000059326 | -0.530370378 | 1.90E-06 |
| Pitpnm1 | ENSMUSG00000024851 | -0.529699694 | 1.05E-10 |
| Mbd6 | ENSMUSG00000025409 | -0.529615935 | 4.54E-06 |
| Acadvl | ENSMUSG00000018574 | -0.52812498 | 2.03E-07 |
| Stk11ip | ENSMUSG00000026213 | -0.528056884 | 6.79E-14 |
| Smad2 | ENSMUSG00000024563 | -0.524103996 | 1.75E-13 |
| Cdan1 | ENSMUSG00000027284 | -0.523772183 | 1.95E-08 |

Supplementary data sheet 3

### Differentially expressed pericyte (PC) genes in the P10 *Apcdd1* -/- retina

#### Significantly upregulated genes

| Gene_name | ID | log2FoldChange | Padj |
|---|---|---|---|
| Cdc37l1 | ENSMUSG00000024780.6 | 0.554913032 | 3.78E-10 |
| Cep250 | ENSMUSG00000038241.13 | 0.544480873 | 2.36E-08 |
| Cfap126 | ENSMUSG00000026649.11 | 1.155168057 | 1.27E-14 |
| Cwc22 | ENSMUSG00000027014.11 | 0.527087346 | 6.99E-04 |
| Dtd1 | ENSMUSG00000027430.9 | 0.502929076 | 1.13E-07 |
| Fxyd6 | ENSMUSG00000066705.6 | 0.666125749 | 4.50E-10 |
| H2-D1 | ENSMUSG00000073411.8 | 2.388354934 | 1.33E-186 |
| H2-K1 | ENSMUSG00000061232.12 | 0.920807862 | 8.63E-12 |
| H2-Q4 | ENSMUSG00000035929.11 | 1.275125296 | 5.65E-28 |
| Hebp2 | ENSMUSG00000019853.5 | 0.554960147 | 1.97E-05 |
| Hist1h2bc | ENSMUSG00000018102.4 | 0.518366657 | 9.64E-06 |
| Hspe1 | ENSMUSG00000073676.4 | 0.587872204 | 1.39E-09 |
| Ifi30 | ENSMUSG00000031838.7 | -0.533926718 | 4.35E-06 |
| Il11ra1 | ENSMUSG00000073889.7 | 1.323589324 | 1.59E-40 |
| Irak1bp1 | ENSMUSG00000032251.9 | 0.591567773 | 3.92E-08 |
| Itga6 | ENSMUSG00000027111.12 | 0.595201244 | 2.48E-11 |
| Lama2 | ENSMUSG00000019899.12 | 0.523332443 | 5.57E-05 |
| Lrpprc | ENSMUSG00000024120.9 | 0.777528424 | 2.77E-38 |
| Moxd1 | ENSMUSG00000020000.7 | 0.530092546 | 2.90E-06 |
| Mt3 | ENSMUSG00000031760.8 | 1.033404476 | 3.23E-22 |
| Ndufa4l2 | ENSMUSG00000040280.9 | 0.596559115 | 1.88E-05 |
| Nos3 | ENSMUSG00000028978.9 | -0.796499121 | 1.44E-06 |
| Nudt19 | ENSMUSG00000034875.5 | 0.741253607 | 1.36E-14 |
| Olfml2a | ENSMUSG00000046618.7 | -0.558525393 | 0.002024602 |
| Pdk1 | ENSMUSG00000006494.8 | 0.525088303 | 9.20E-17 |
| Rab27b | ENSMUSG00000024511.12 | 0.595012147 | 1.04E-13 |
| Rnaset2b | ENSMUSG00000094724.4 | 0.85925806 | 1.55E-07 |
| Slc29a1 | ENSMUSG00000023942.12 | 0.622051 | 4.47E-04 |
| Sspn | ENSMUSG00000030255.10 | 0.518189587 | 3.58E-06 |
| Syt15 | ENSMUSG00000041479.10 | 1.409765644 | 5.88E-47 |
| Tuba4a | ENSMUSG00000026202.10 | 0.819390438 | 1.23E-14 |
| Vcp | ENSMUSG00000028452.7 | 0.876030174 | 3.18E-29 |
| Zfp738 | ENSMUSG00000048280.14 | 0.853811659 | 6.58E-30 |

**Differentially expressed pericyte (PC) genes in the P10 *Apcdd1* -/- retina**

**Significantly downregulated genes**

| Gene_name | ID | log2FoldChange | padj |
|---|---|---|---|
| Acaca | ENSMUSG00000020532.15 | -0.514300121 | 1.77E-05 |
| Acot1 | ENSMUSG00000072949.5 | -0.599754484 | 1.69E-05 |
| Ankrd23 | ENSMUSG00000067653.8 | -0.575135622 | 8.73E-04 |
| Apcdd1 | ENSMUSG00000071847.9 | -0.837383081 | 6.75E-31 |
| Aurka | ENSMUSG00000027496.12 | -1.411782715 | 2.31E-35 |
| Ccdc171 | ENSMUSG00000052407.13 | -1.118809169 | 1.83E-20 |
| Dctn3 | ENSMUSG00000028447.8 | -0.503775922 | 6.96E-05 |
| Dxo | ENSMUSG00000040482.12 | -0.576322828 | 8.40E-15 |
| Efhc1 | ENSMUSG00000041809.5 | -0.642002398 | 1.24E-04 |
| Fam210b | ENSMUSG00000027495.4 | -0.913504807 | 1.15E-40 |
| Gdpd3 | ENSMUSG00000030703.7 | -0.557088638 | 0.002187315 |
| Gem | ENSMUSG00000028214.10 | -0.917873857 | 1.21E-19 |
| Gstp1 | ENSMUSG00000060803.5 | -0.509636682 | 2.59E-08 |
| Haus5 | ENSMUSG00000078762.7 | -1.043704145 | 1.84E-12 |
| Hexdc | ENSMUSG00000039307.13 | -0.614404354 | 1.58E-14 |
| Hist1h2al | ENSMUSG00000091383.1 | -0.86919983 | 1.15E-08 |
| Itgb3bp | ENSMUSG00000028549.14 | -0.734683823 | 1.89E-10 |
| Malt1 | ENSMUSG00000032688.7 | -0.712375007 | 1.17E-26 |
| Myo1d | ENSMUSG00000035441.11 | -0.563983532 | 8.91E-11 |
| Pgap2 | ENSMUSG00000030990.14 | -0.941898609 | 2.78E-17 |
| Prcp | ENSMUSG00000061119.6 | -0.529616729 | 3.18E-06 |
| Ring1 | ENSMUSG00000024325.8 | -0.676803013 | 7.62E-14 |
| Rreb1 | ENSMUSG00000039087.13 | -0.718518234 | 1.93E-20 |
| Samhd1 | ENSMUSG00000027639.13 | -0.536330446 | 3.35E-09 |
| Stk19 | ENSMUSG00000061207.8 | -0.816553797 | 3.61E-18 |
| Stx3 | ENSMUSG00000041488.12 | -0.621444848 | 5.03E-17 |
| Tpm3-rs7 | ENSMUSG00000058126.6 | -3.570676155 | 2.29E-236 |
| Trim12c | ENSMUSG00000057143.11 | -1.440058456 | 6.77E-26 |

**Differentially expressed pericyte (PC) genes in the P14 *Apcdd1* -/- retina**

**Significantly upregulated genes**

| Gene_name | ID | log2FoldChange | Padj |
|---|---|---|---|
| Abca9 | ENSMUSG00000041797 | 0.673314256 | 0.00305537 |
| Ackr3 | ENSMUSG00000044337 | 0.630385513 | 4.88E-05 |
| Adm | ENSMUSG00000030790 | 0.661363993 | 0.01284728 |
| Agmo | ENSMUSG00000050103 | 0.68752633 | 0.04928382 |
| Ahnak | ENSMUSG00000069833 | 0.667311148 | 0.00172803 |
| Angptl2 | ENSMUSG00000004105 | 0.904556735 | 0.00012213 |
| Antxr1 | ENSMUSG00000033420 | 0.780545812 | 4.67E-09 |
| Anxa2 | ENSMUSG00000032231 | 0.511944245 | 5.15E-07 |
| Anxa5 | ENSMUSG00000027712 | 0.918951093 | 9.78E-19 |
| Apbb1ip | ENSMUSG00000026786 | 0.988410881 | 0.03571352 |
| Apoe | ENSMUSG00000002985 | 0.584954013 | 8.91E-11 |
| Arhgdib | ENSMUSG00000030220 | 0.596405679 | 0.00076883 |
| Arpc1b | ENSMUSG00000029622 | 0.589486804 | 1.01E-06 |
| Arsk | ENSMUSG00000021592 | 0.535215243 | 0.00014244 |
| Atp1a2 | ENSMUSG00000007097 | 0.820246224 | 1.13E-08 |
| Atp1b3 | ENSMUSG00000032412 | 0.670376039 | 3.75E-10 |
| Bace2 | ENSMUSG00000040605 | 0.694630768 | 0.0001508 |
| Baiap2l1 | ENSMUSG00000038859 | 0.62797116 | 0.04558007 |
| Bambi | ENSMUSG00000024232 | 0.556284457 | 5.04E-07 |
| BC022687 | ENSMUSG00000037594 | 0.573564711 | 2.75E-08 |
| Bhlhe40 | ENSMUSG00000030103 | 0.544757488 | 5.03E-07 |
| Bmp4 | ENSMUSG00000021835 | 0.739085424 | 0.00065171 |
| Ccdc80 | ENSMUSG00000022665 | 0.872369723 | 5.02E-05 |
| Ccnd2 | ENSMUSG00000000184 | 0.817209782 | 9.45E-05 |
| Cd300a | ENSMUSG00000034652 | 1.110200568 | 0.00403279 |
| Cd320 | ENSMUSG00000002308 | 0.529756015 | 8.03E-07 |
| Cd44 | ENSMUSG00000005087 | 0.789963393 | 1.08E-05 |
| Cd63 | ENSMUSG00000025351 | 0.517079516 | 9.67E-10 |
| Cdc37l1 | ENSMUSG00000024780 | 0.928987836 | 5.59E-35 |
| Cebpa | ENSMUSG00000034957 | 0.713447857 | 0.01568274 |
| Cebpb | ENSMUSG00000056501 | 0.724255096 | 0.00386757 |
| Cenpf | ENSMUSG00000026605 | 0.74756641 | 0.01869037 |
| Cep250 | ENSMUSG00000038241 | 0.543653654 | 5.04E-09 |
| Cfap126 | ENSMUSG00000026649 | 0.796036414 | 2.95E-05 |
| Cgnl1 | ENSMUSG00000032232 | 0.51676679 | 0.00055365 |
| Cmtm7 | ENSMUSG00000032436 | 0.960927924 | 3.62E-05 |
| Col10a1 | ENSMUSG00000039462 | 1.019678078 | 0.00447051 |
| Col18a1 | ENSMUSG00000001435 | 0.805157116 | 0.0404106 |
| Col4a5 | ENSMUSG00000031274 | 0.759786081 | 0.00018587 |
| Col4a6 | ENSMUSG00000031273 | 1.283940675 | 0.00198635 |

| | | | |
|---|---|---|---|
| Col6a3 | ENSMUSG00000048126 | 1.165729253 | 0.00155246 |
| Col9a1 | ENSMUSG00000026147 | 0.629593123 | 7.79E-07 |
| Colec12 | ENSMUSG00000036103 | 1.176767216 | 7.62E-08 |
| Colgalt2 | ENSMUSG00000032649 | 0.630489529 | 1.14E-10 |
| Cp | ENSMUSG00000003617 | 0.708888869 | 3.86E-08 |
| Crim1 | ENSMUSG00000024074 | 0.540888707 | 0.00030928 |
| Crip1 | ENSMUSG00000006360 | 0.663030455 | 0.01127085 |
| Csf1 | ENSMUSG00000014599 | 0.591530538 | 0.00459021 |
| Cst3 | ENSMUSG00000027447 | 0.56866207 | 1.95E-08 |
| Ctgf | ENSMUSG00000019997 | 0.861809412 | 6.81E-07 |
| Ctsc | ENSMUSG00000030560 | 0.617365811 | 1.74E-08 |
| Ctsh | ENSMUSG00000032359 | 0.694126793 | 1.35E-07 |
| Ctsk | ENSMUSG00000028111 | 0.589596718 | 0.04154892 |
| Cwc22 | ENSMUSG00000027014 | 3.609834916 | 4.13E-24 |
| Cyba | ENSMUSG00000006519 | 0.696929368 | 1.70E-05 |
| Cyp2j9 | ENSMUSG00000015224 | 0.904282761 | 0.00034649 |
| Dse | ENSMUSG00000039497 | 0.684687083 | 0.0008346 |
| Dtd1 | ENSMUSG00000027430 | 0.846058865 | 6.59E-14 |
| Dtl | ENSMUSG00000037474 | 1.066053113 | 0.01548692 |
| Dydc2 | ENSMUSG00000021791 | 2.410831382 | 2.43E-06 |
| Ech1 | ENSMUSG00000053898 | 0.62665877 | 1.54E-14 |
| Ecm1 | ENSMUSG00000028108 | 0.580854177 | 0.0005673 |
| Efemp1 | ENSMUSG00000020467 | 0.682933524 | 1.98E-10 |
| Efna1 | ENSMUSG00000027954 | 0.503281376 | 0.00013067 |
| Egfl8 | ENSMUSG00000015467 | 1.110587307 | 7.79E-09 |
| Eif4ebp1 | ENSMUSG00000031490 | 1.066726141 | 3.37E-08 |
| Elf4 | ENSMUSG00000031103 | 0.775297859 | 0.04406086 |
| Emp2 | ENSMUSG00000022505 | 0.86273157 | 6.51E-11 |
| Emp3 | ENSMUSG00000040212 | 0.655145496 | 0.00931323 |
| En2 | ENSMUSG00000039095 | 0.624080247 | 0.02401918 |
| Enpp1 | ENSMUSG00000037370 | 0.842126522 | 0.00013925 |
| Fam107a | ENSMUSG00000021750 | 0.682192083 | 0.03033048 |
| Fam114a1 | ENSMUSG00000029185 | 0.561421077 | 0.00083438 |
| Fbln1 | ENSMUSG00000006369 | 1.382214811 | 0.00788349 |
| Fbln2 | ENSMUSG00000064080 | 1.269181266 | 4.85E-09 |
| Fcgrt | ENSMUSG00000003420 | 0.516472919 | 0.00124559 |
| Fkbp9 | ENSMUSG00000029781 | 0.522430293 | 8.57E-07 |
| Fos | ENSMUSG00000021250 | 1.30794222 | 0.0354153 |
| Fosb | ENSMUSG00000003545 | 1.70050788 | 0.02595726 |
| Frk | ENSMUSG00000019779 | 0.701322807 | 0.02386451 |
| Fstl1 | ENSMUSG00000022816 | 0.921403692 | 6.23E-12 |
| Fxyd6 | ENSMUSG00000066705 | 0.629202382 | 8.84E-05 |
| Gas5 | ENSMUSG00000053332 | 0.557821064 | 8.24E-06 |
| Gfap | ENSMUSG00000020932 | 1.276516329 | 0.01551296 |
| Gja1 | ENSMUSG00000050953 | 0.901632998 | 0.01521701 |

| | | | |
|---|---|---|---|
| Gja5 | ENSMUSG00000057123 | 1.189320041 | 0.0002979 |
| Gng11 | ENSMUSG00000032766 | 0.546729958 | 0.0007345 |
| Golm1 | ENSMUSG00000021556 | 0.533589325 | 0.00732682 |
| Gpr137b | ENSMUSG00000021306 | 1.128420356 | 2.64E-24 |
| Gprc5c | ENSMUSG00000051043 | 0.732522334 | 0.00212863 |
| Gpx8 | ENSMUSG00000021760 | 0.691421425 | 1.06E-14 |
| Gstm1 | ENSMUSG00000058135 | 1.218773352 | 5.65E-09 |
| Gstm2 | ENSMUSG00000040562 | 1.789107874 | 0.00303107 |
| Gusb | ENSMUSG00000025534 | 0.528349244 | 0.00012925 |
| Gypc | ENSMUSG00000090523 | 0.939084001 | 0.00338326 |
| H2-D1 | ENSMUSG00000073411 | 2.090209147 | 1.85E-98 |
| H2-K1 | ENSMUSG00000061232 | 1.081186286 | 1.32E-18 |
| H2-Q4 | ENSMUSG00000035929 | 1.552829397 | 4.48E-42 |
| H60b | ENSMUSG00000075297 | 1.564967446 | 0.00124559 |
| Hebp2 | ENSMUSG00000019853 | 0.613551677 | 1.41E-05 |
| Herc6 | ENSMUSG00000029798 | 0.538353623 | 0.03254669 |
| Hexb | ENSMUSG00000021665 | 0.703611509 | 0.00013925 |
| Hfe | ENSMUSG00000006611 | 0.584042062 | 3.75E-09 |
| Hhex | ENSMUSG00000024986 | 0.689495158 | 0.02739009 |
| Hist1h2bc | ENSMUSG00000018102 | 0.841745908 | 9.94E-09 |
| Hpgds | ENSMUSG00000029919 | 0.931074286 | 0.01135332 |
| Hrct1 | ENSMUSG00000071001 | 1.057520905 | 0.00186559 |
| Hspb2 | ENSMUSG00000038086 | 1.187442123 | 0.00334527 |
| Hspb8 | ENSMUSG00000041548 | 0.500554422 | 0.00388364 |
| Hspe1 | ENSMUSG00000073676 | 0.628098425 | 3.07E-07 |
| Hspg2 | ENSMUSG00000028763 | 0.991144861 | 0.02626044 |
| Ifi30 | ENSMUSG00000031838 | 0.558556317 | 5.86E-05 |
| Ifitm1 | ENSMUSG00000025491 | 1.229138913 | 0.00504677 |
| Ifitm2 | ENSMUSG00000060591 | 0.946441783 | 3.76E-11 |
| Ifitm3 | ENSMUSG00000025492 | 0.758566168 | 3.84E-06 |
| Igf2 | ENSMUSG00000048583 | 0.598124804 | 3.30E-17 |
| Igf2bp1 | ENSMUSG00000013415 | 0.977306549 | 0.00838845 |
| Igfbp2 | ENSMUSG00000039323 | 0.565420238 | 6.10E-06 |
| Igfbp7 | ENSMUSG00000036256 | 0.792993356 | 4.05E-05 |
| Il10rb | ENSMUSG00000022969 | 0.554585863 | 0.00083842 |
| Il11ra1 | ENSMUSG00000073889 | 1.670649256 | 8.13E-52 |
| Il1r1 | ENSMUSG00000026072 | 0.6334096 | 4.40E-05 |
| Iqgap2 | ENSMUSG00000021676 | 0.582390192 | 0.00552437 |
| Irak1bp1 | ENSMUSG00000032251 | 0.578062185 | 0.00297862 |
| Islr | ENSMUSG00000037206 | 1.111026998 | 0.00801283 |
| Itga6 | ENSMUSG00000027111 | 0.874797958 | 1.43E-09 |
| Itih5 | ENSMUSG00000025780 | 0.580188802 | 0.00051732 |
| Itm2a | ENSMUSG00000031239 | 0.555554936 | 2.49E-09 |
| Kcns3 | ENSMUSG00000043673 | 0.886680271 | 0.00478429 |
| Kdelr3 | ENSMUSG00000010830 | 0.695671608 | 0.004485 |

| | | | |
|---|---|---|---|
| Kitl | ENSMUSG00000019966 | 0.701416132 | 1.05E-15 |
| Klhl6 | ENSMUSG00000043008 | 0.661117263 | 0.01709747 |
| Lama2 | ENSMUSG00000019899 | 1.024020241 | 1.41E-05 |
| Lgals3 | ENSMUSG00000050335 | 0.832827307 | 2.20E-08 |
| Lipa | ENSMUSG00000024781 | 0.500909301 | 1.13E-08 |
| Loxl1 | ENSMUSG00000032334 | 0.755923928 | 0.00031677 |
| Lrpprc | ENSMUSG00000024120 | 1.16849876 | 3.48E-65 |
| Ltbp1 | ENSMUSG00000001870 | 0.837608829 | 0.00043299 |
| Lyn | ENSMUSG00000042228 | 0.551013622 | 0.00372338 |
| Lysmd1 | ENSMUSG00000053769 | 0.600452799 | 3.40E-07 |
| Marveld2 | ENSMUSG00000021636 | 0.51315534 | 0.00948718 |
| Masp2 | ENSMUSG00000028979 | 1.606786781 | 0.005733 |
| Matn2 | ENSMUSG00000022324 | 0.668640936 | 0.00193311 |
| Metap1d | ENSMUSG00000041921 | 0.736408035 | 1.61E-07 |
| Mfap2 | ENSMUSG00000060572 | 0.63265134 | 2.16E-05 |
| Mki67 | ENSMUSG00000031004 | 0.798122757 | 0.01128756 |
| Mlc1 | ENSMUSG00000035805 | 0.614044318 | 1.18E-14 |
| Mmp2 | ENSMUSG00000031740 | 0.627884766 | 5.10E-06 |
| Moxd1 | ENSMUSG00000020000 | 0.56861417 | 2.45E-07 |
| Mrc1 | ENSMUSG00000026712 | 2.420444055 | 2.52E-09 |
| Mrc2 | ENSMUSG00000020695 | 0.585414443 | 0.00307837 |
| Mrps6 | ENSMUSG00000039680 | 0.536482086 | 2.96E-06 |
| Msx1 | ENSMUSG00000048450 | 1.151656255 | 7.62E-05 |
| mt-Te | ENSMUSG00000064369 | 0.572205155 | 0.00262433 |
| Mt3 | ENSMUSG00000031760 | 1.044478444 | 2.70E-17 |
| Mtap | ENSMUSG00000062937 | 0.85216434 | 3.44E-07 |
| Myl9 | ENSMUSG00000067818 | 0.705971152 | 8.49E-12 |
| Ndc80 | ENSMUSG00000024056 | 1.257749786 | 0.03031214 |
| Ndufa4l2 | ENSMUSG00000040280 | 0.822001567 | 3.38E-05 |
| Nexn | ENSMUSG00000039103 | 0.67303593 | 0.04275988 |
| Nhsl1 | ENSMUSG00000039835 | 0.55861205 | 0.0002128 |
| Nid1 | ENSMUSG00000005397 | 0.931835878 | 0.00610575 |
| Nkd2 | ENSMUSG00000021567 | 0.742514976 | 0.0003696 |
| Nme1 | ENSMUSG00000037601 | 0.700971446 | 4.71E-13 |
| Nos3 | ENSMUSG00000028978 | 1.787594364 | 2.17E-06 |
| Nudt19 | ENSMUSG00000034875 | 1.059135142 | 3.77E-43 |
| Nupr1 | ENSMUSG00000030717 | 0.798888859 | 0.00013923 |
| Ogn | ENSMUSG00000021390 | 1.047494775 | 2.03E-05 |
| Olfml2a | ENSMUSG00000046618 | 0.581403347 | 0.04022057 |
| P2ry12 | ENSMUSG00000036353 | 0.79892026 | 0.000531 |
| Pbk | ENSMUSG00000022033 | 1.160164893 | 0.01447432 |
| Pcolce | ENSMUSG00000029718 | 1.038902907 | 5.17E-08 |
| Pdk1 | ENSMUSG00000006494 | 0.79908467 | 3.10E-24 |
| Peg10 | ENSMUSG00000092035 | 0.623013619 | 0.00397014 |
| Perp | ENSMUSG00000019851 | 1.515513357 | 1.85E-26 |

| | | | |
|---|---|---|---|
| Phex | ENSMUSG00000057457 | 1.288830124 | 0.03350989 |
| Pign | ENSMUSG00000056536 | 0.613465901 | 2.29E-06 |
| Pla2g4a | ENSMUSG00000056220 | 1.334802516 | 1.81E-07 |
| Plau | ENSMUSG00000021822 | 1.112059483 | 0.00834209 |
| Plekhf1 | ENSMUSG00000074170 | 0.82859868 | 4.08E-05 |
| Pmp22 | ENSMUSG00000018217 | 0.518310451 | 2.05E-08 |
| Pon3 | ENSMUSG00000029759 | 0.928811902 | 2.02E-05 |
| Ppic | ENSMUSG00000024538 | 0.601602874 | 6.70E-06 |
| Ppp1r3b | ENSMUSG00000046794 | 0.583899861 | 0.0067462 |
| Prelp | ENSMUSG00000041577 | 0.506744501 | 0.00071721 |
| Psmb9 | ENSMUSG00000096727 | 0.759530414 | 0.01497576 |
| Psmc3ip | ENSMUSG00000019303 | 0.824511175 | 0.0002005 |
| Ptgfr | ENSMUSG00000028036 | 0.80074999 | 0.04027961 |
| Ptgs1 | ENSMUSG00000047250 | 0.601733294 | 0.040691 |
| Ptpn14 | ENSMUSG00000026604 | 0.501187067 | 0.01918108 |
| Pxdn | ENSMUSG00000020674 | 0.660743742 | 3.41E-06 |
| Rab27b | ENSMUSG00000024511 | 0.567343773 | 1.13E-12 |
| Ramp1 | ENSMUSG00000034353 | 0.639035946 | 0.0002462 |
| Rarres2 | ENSMUSG00000009281 | 0.843843295 | 6.75E-05 |
| Rassf3 | ENSMUSG00000025795 | 0.809144703 | 5.25E-09 |
| Rcn3 | ENSMUSG00000019539 | 0.644219313 | 7.98E-06 |
| Rhod | ENSMUSG00000041845 | 0.817197662 | 1.31E-07 |
| Rnase4 | ENSMUSG00000021876 | 0.909971145 | 0.0035933 |
| Rnaset2b | ENSMUSG00000094724 | 4.066275029 | 8.84E-74 |
| Rny3 | ENSMUSG00000064945 | 0.733877377 | 0.04018942 |
| Rpl22l1 | ENSMUSG00000039221 | 0.650451604 | 4.31E-07 |
| Rpl35a | ENSMUSG00000060636 | 0.918976055 | 2.28E-19 |
| Rps3a2 | ENSMUSG00000062611 | 4.100287793 | 7.03E-44 |
| Rps3a3 | ENSMUSG00000059751 | 7.352887321 | 6.01E-201 |
| Rrm2 | ENSMUSG00000020649 | 0.610398915 | 0.0262763 |
| Rwdd3 | ENSMUSG00000028133 | 0.568262512 | 4.43E-05 |
| S100a10 | ENSMUSG00000041959 | 0.68257472 | 5.82E-09 |
| S100a11 | ENSMUSG00000027907 | 0.666569361 | 0.00081494 |
| S100a4 | ENSMUSG00000001020 | 1.016141116 | 0.00157684 |
| S100a6 | ENSMUSG00000001025 | 1.288018029 | 7.85E-12 |
| S1pr3 | ENSMUSG00000067586 | 0.600515479 | 6.13E-05 |
| Sdc1 | ENSMUSG00000020592 | 0.586488032 | 0.02211185 |
| Serpine1 | ENSMUSG00000037411 | 1.124201947 | 1.73E-07 |
| Serpinf1 | ENSMUSG00000000753 | 1.067388633 | 1.65E-10 |
| Serping1 | ENSMUSG00000023224 | 1.220599464 | 4.85E-09 |
| Six2 | ENSMUSG00000024134 | 0.537989701 | 0.03093853 |
| Slc15a2 | ENSMUSG00000022899 | 0.860240247 | 6.17E-34 |
| Slc16a12 | ENSMUSG00000009378 | 1.100564726 | 3.74E-05 |
| Slc22a8 | ENSMUSG00000063796 | 0.504174693 | 0.00518332 |
| Slc25a45 | ENSMUSG00000024818 | 0.716710543 | 0.02842455 |

| | | | |
|---|---|---|---|
| Slc26a11 | ENSMUSG00000039908 | 1.498214654 | 1.34E-43 |
| Slc29a1 | ENSMUSG00000023942 | 0.756884661 | 0.01853492 |
| Slc38a4 | ENSMUSG00000022464 | 1.290902481 | 4.75E-05 |
| Slc5a5 | ENSMUSG00000000792 | 0.922457361 | 3.32E-10 |
| Slc8b1 | ENSMUSG00000032754 | 0.620628192 | 3.31E-05 |
| Slit3 | ENSMUSG00000056427 | 1.226470085 | 0.00217061 |
| Smagp | ENSMUSG00000053559 | 0.867269392 | 0.01443947 |
| Smco4 | ENSMUSG00000058173 | 0.623694266 | 0.01154656 |
| Sparc | ENSMUSG00000018593 | 0.534698888 | 2.70E-21 |
| Spon2 | ENSMUSG00000037379 | 0.871073605 | 1.75E-10 |
| Spp1 | ENSMUSG00000029304 | 2.415034388 | 0.00525567 |
| Sri | ENSMUSG00000003161 | 0.539828322 | 1.73E-13 |
| Sspn | ENSMUSG00000030255 | 0.850147071 | 6.08E-07 |
| Suclg2 | ENSMUSG00000061838 | 0.501563008 | 0.00092556 |
| Syt15 | ENSMUSG00000041479 | 1.374086256 | 2.57E-30 |
| Tagln2 | ENSMUSG00000026547 | 0.524144417 | 0.00486834 |
| Tec | ENSMUSG00000029217 | 0.633350154 | 0.01065947 |
| Tfpi | ENSMUSG00000027082 | 1.484475813 | 9.95E-14 |
| Tgif1 | ENSMUSG00000047407 | 0.577446318 | 0.00316987 |
| Thbd | ENSMUSG00000074743 | 0.512581936 | 0.02810901 |
| Thbs2 | ENSMUSG00000023885 | 0.530587028 | 0.01705699 |
| Tinagl1 | ENSMUSG00000028776 | 1.794251824 | 1.48E-10 |
| Tk1 | ENSMUSG00000025574 | 0.734829134 | 0.01490851 |
| Tmbim1 | ENSMUSG00000006301 | 0.607409978 | 4.22E-09 |
| Tmem37 | ENSMUSG00000050777 | 0.862412199 | 3.40E-05 |
| Tnfaip8 | ENSMUSG00000062210 | 0.619110067 | 4.28E-10 |
| Tnfrsf12a | ENSMUSG00000023905 | 0.614999667 | 0.00097249 |
| Tnfrsf1b | ENSMUSG00000028599 | 0.562224688 | 0.03163298 |
| Top2a | ENSMUSG00000020914 | 0.52800728 | 0.03019222 |
| Tspo | ENSMUSG00000041736 | 0.750605322 | 1.70E-07 |
| Tuba4a | ENSMUSG00000026202 | 0.610048055 | 0.04878346 |
| Txnip | ENSMUSG00000038393 | 0.841887213 | 0.0001187 |
| Vcp | ENSMUSG00000028452 | 0.687173204 | 4.64E-24 |
| Wnt5b | ENSMUSG00000030170 | 0.513667319 | 0.00013723 |
| Zcchc9 | ENSMUSG00000021621 | 0.632953226 | 2.35E-12 |
| Zfp36 | ENSMUSG00000044786 | 0.570392991 | 0.00100244 |
| Zfp429 | ENSMUSG00000078994 | 0.817660767 | 0.00786133 |
| Zfp738 | ENSMUSG00000048280 | 1.001088578 | 1.64E-19 |
| Zfp758 | ENSMUSG00000044501 | 0.564847382 | 0.00052589 |
| Zic1 | ENSMUSG00000032368 | 0.852381046 | 0.0001068 |
| Zic4 | ENSMUSG00000036972 | 1.235011777 | 4.36E-05 |

**Differentially expressed pericyte (PC) genes in the P14 *Apcdd1* -/- retina**

**Significantly downregulated genes**

| Gene_name | ID | log2FoldChange | Padj |
|---|---|---|---|
| AA386476 | ENSMUSG00000074357 | -0.528053886 | 0.00203557 |
| Abca4 | ENSMUSG00000028125 | -0.557879456 | 9.82E-08 |
| Abhd14b | ENSMUSG00000042073 | -0.574422867 | 1.29E-07 |
| Acadvl | ENSMUSG00000018574 | -0.52812498 | 2.03E-07 |
| Acot1 | ENSMUSG00000072949 | -0.953252782 | 5.42E-12 |
| Acrbp | ENSMUSG00000072770 | -0.504311284 | 0.00122068 |
| Aipl1 | ENSMUSG00000040554 | -0.512272478 | 6.22E-06 |
| Ankrd23 | ENSMUSG00000067653 | -0.710984053 | 0.00602623 |
| Apcdd1 | ENSMUSG00000071847 | -0.738717045 | 1.89E-26 |
| Asic3 | ENSMUSG00000038276 | -0.622350262 | 2.11E-09 |
| Aurka | ENSMUSG00000027496 | -1.145364461 | 4.55E-16 |
| Bcar3 | ENSMUSG00000028121 | -0.574629516 | 1.53E-06 |
| Cables1 | ENSMUSG00000040957 | -0.557155806 | 3.57E-09 |
| Caprin2 | ENSMUSG00000030309 | -0.509665963 | 0.00019951 |
| Ccdc107 | ENSMUSG00000028461 | -0.502760934 | 5.60E-06 |
| Ccdc163 | ENSMUSG00000028689 | -0.719833686 | 1.73E-10 |
| Ccdc171 | ENSMUSG00000052407 | -1.253616039 | 1.24E-15 |
| Chtf18 | ENSMUSG00000019214 | -0.62573507 | 3.84E-06 |
| Clasrp | ENSMUSG00000061028 | -0.657840539 | 1.10E-11 |
| Clstn1 | ENSMUSG00000039953 | -0.828336899 | 3.20E-55 |
| Col27a1 | ENSMUSG00000045672 | -0.838194001 | 1.11E-13 |
| Col6a1 | ENSMUSG00000001119 | -0.816607391 | 3.13E-09 |
| Cpt1a | ENSMUSG00000024900 | -0.535706174 | 7.25E-07 |
| Crhr2 | ENSMUSG00000003476 | -0.547200669 | 0.00351124 |
| Ctu2 | ENSMUSG00000049482 | -0.513684127 | 1.15E-06 |
| Dctn3 | ENSMUSG00000028447 | -0.900143271 | 5.83E-33 |
| Dennd1c | ENSMUSG00000002668 | -1.741063004 | 1.99E-05 |
| Dennd6b | ENSMUSG00000015377 | -0.500912326 | 1.72E-05 |
| Dhx35 | ENSMUSG00000027655 | -0.635867998 | 3.42E-21 |
| Dnah9 | ENSMUSG00000056752 | -0.539617541 | 4.64E-05 |
| Dxo | ENSMUSG00000040482 | -0.797440171 | 1.90E-28 |
| Dyrk2 | ENSMUSG00000028630 | -0.56501104 | 0.00038939 |
| Efhc1 | ENSMUSG00000041809 | -1.499155219 | 4.47E-08 |
| Fam110a | ENSMUSG00000027459 | -0.987190883 | 3.62E-42 |
| Fam193b | ENSMUSG00000021495 | -0.625937385 | 1.12E-09 |
| Fam210b | ENSMUSG00000027495 | -0.758281931 | 1.54E-28 |
| Fnbp4 | ENSMUSG00000008200 | -0.5109324 | 8.87E-11 |
| Fpgs | ENSMUSG00000009566 | -0.50420434 | 5.32E-05 |
| Gdpd3 | ENSMUSG00000030703 | -3.704505962 | 2.39E-45 |
| Gem | ENSMUSG00000028214 | -1.112609495 | 3.03E-39 |

| | | | |
|---|---|---|---|
| Gigyf1 | ENSMUSG00000029714 | -0.53751981 | 1.90E-06 |
| Gpr137b-ps | ENSMUSG00000097715 | -1.455830473 | 6.00E-24 |
| Gpsm3 | ENSMUSG00000034786 | -0.785766182 | 0.01983805 |
| Gstp1 | ENSMUSG00000060803 | -0.694615541 | 1.18E-10 |
| Gucy2e | ENSMUSG00000020890 | -0.618986938 | 7.23E-11 |
| Haus5 | ENSMUSG00000078762 | -0.798341684 | 6.65E-06 |
| Hexdc | ENSMUSG00000039307 | -0.583501761 | 5.32E-12 |
| Hist1h2al | ENSMUSG00000091383 | -1.450278602 | 1.14E-47 |
| Hmgb2 | ENSMUSG00000054717 | -0.540097896 | 2.95E-06 |
| Hspa1l | ENSMUSG00000007033 | -0.585194935 | 0.04636931 |
| Igsf21 | ENSMUSG00000040972 | -0.689880743 | 7.17E-14 |
| Igsf9 | ENSMUSG00000037995 | -0.630575812 | 7.04E-13 |
| Il3ra | ENSMUSG00000068758 | -0.773961298 | 2.68E-06 |
| Impg2 | ENSMUSG00000035270 | -0.670192445 | 1.23E-11 |
| Ing4 | ENSMUSG00000030330 | -0.566432011 | 0.00899325 |
| Itgb3bp | ENSMUSG00000028549 | -0.682226846 | 2.21E-05 |
| Jmjd7 | ENSMUSG00000098789 | -0.687495831 | 0.0059278 |
| Kbtbd4 | ENSMUSG00000005505 | -0.559483928 | 2.23E-08 |
| Khsrp | ENSMUSG00000007670 | -0.621681865 | 1.41E-20 |
| Kif21b | ENSMUSG00000041642 | -0.608850766 | 4.07E-05 |
| Kifc3 | ENSMUSG00000031788 | -0.515419747 | 4.71E-12 |
| L3hypdh | ENSMUSG00000019718 | -0.587035308 | 9.44E-09 |
| Leng8 | ENSMUSG00000035545 | -0.577536215 | 4.00E-05 |
| Ly6g6d | ENSMUSG00000073413 | -0.805271352 | 0.02262851 |
| Malt1 | ENSMUSG00000032688 | -0.702638958 | 2.64E-06 |
| Mbd6 | ENSMUSG00000025409 | -0.529615935 | 4.54E-06 |
| Myo1d | ENSMUSG00000035441 | -0.857001795 | 1.07E-11 |
| Myocd | ENSMUSG00000020542 | -0.593384177 | 0.03425046 |
| Nr1h3 | ENSMUSG00000002108 | -0.837490798 | 5.15E-05 |
| Nudt8 | ENSMUSG00000024869 | -0.903205204 | 5.64E-05 |
| Pde6b | ENSMUSG00000029491 | -0.853383039 | 7.22E-06 |
| Pdzrn3 | ENSMUSG00000035357 | -0.579041603 | 4.20E-05 |
| Pfkfb2 | ENSMUSG00000026409 | -0.536841023 | 1.00E-13 |
| Pgap2 | ENSMUSG00000030990 | -0.831797989 | 1.05E-05 |
| Pitpnm1 | ENSMUSG00000024851 | -0.529699694 | 1.05E-10 |
| Plcd3 | ENSMUSG00000020937 | -0.540997294 | 2.22E-09 |
| Plch2 | ENSMUSG00000029055 | -0.560097983 | 1.07E-13 |
| Plvap | ENSMUSG00000034845 | -0.980562775 | 9.56E-11 |
| Polm | ENSMUSG00000020474 | -0.556215371 | 1.64E-05 |
| Ppef2 | ENSMUSG00000029410 | -0.560839477 | 2.77E-10 |
| Prcp | ENSMUSG00000061119 | -0.701929877 | 2.40E-08 |
| Prkab1 | ENSMUSG00000029513 | -0.562089571 | 1.47E-08 |
| Prss41 | ENSMUSG00000024114 | -0.838464119 | 6.05E-07 |
| Psd | ENSMUSG00000037126 | -0.50942962 | 4.71E-13 |
| Ptpmt1 | ENSMUSG00000063235 | -0.909807886 | 9.70E-13 |

| | | | |
|---|---|---|---|
| Pvr | ENSMUSG00000040511 | -0.628296715 | 3.14E-11 |
| Rabgef1 | ENSMUSG00000025340 | -0.633994858 | 1.23E-08 |
| Radil | ENSMUSG00000029576 | -0.543373417 | 2.09E-07 |
| Rec8 | ENSMUSG00000002324 | -1.109584903 | 0.00623423 |
| Rfx1 | ENSMUSG00000031706 | -0.539852113 | 1.01E-08 |
| Ring1 | ENSMUSG00000024325 | -0.884910138 | 1.19E-39 |
| Rnf207 | ENSMUSG00000058498 | -1.019467089 | 1.26E-16 |
| Rps3a1 | ENSMUSG00000028081 | -0.667251588 | 7.53E-19 |
| Rpsa-ps10 | ENSMUSG00000047676 | -3.895177598 | 0.00548932 |
| Rreb1 | ENSMUSG00000039087 | -1.128640345 | 1.15E-12 |
| Rrp1b | ENSMUSG00000058392 | -0.505780983 | 9.31E-11 |
| Rtel1 | ENSMUSG00000038685 | -0.721849452 | 1.03E-13 |
| Saal1 | ENSMUSG00000006763 | -0.674916852 | 2.90E-14 |
| Samd11 | ENSMUSG00000096351 | -0.952704051 | 1.55E-19 |
| Samhd1 | ENSMUSG00000027639 | -0.549407362 | 1.41E-11 |
| Selenbp2 | ENSMUSG00000068877 | -2.529096681 | 1.13E-11 |
| Sema4c | ENSMUSG00000026121 | -0.64733993 | 1.29E-13 |
| Sh3pxd2b | ENSMUSG00000040711 | -0.577449901 | 0.0014947 |
| Smad2 | ENSMUSG00000024563 | -0.524103996 | 1.75E-13 |
| Smug1 | ENSMUSG00000036061 | -0.614528512 | 6.01E-22 |
| Snx22 | ENSMUSG00000039452 | -0.691133707 | 0.03271533 |
| Spata6 | ENSMUSG00000034401 | -0.640228846 | 8.58E-08 |
| Srp54b | ENSMUSG00000079108 | -0.648779816 | 8.22E-21 |
| St6galnac2 | ENSMUSG00000110170 | -0.734578793 | 0.0330748 |
| Stard9 | ENSMUSG00000033705 | -0.551564202 | 0.002862 |
| Stk11ip | ENSMUSG00000026213 | -0.528056884 | 6.79E-14 |
| Stk19 | ENSMUSG00000061207 | -1.205132554 | 4.60E-38 |
| Stom | ENSMUSG00000026880 | -0.618691769 | 1.40E-12 |
| Stx3 | ENSMUSG00000041488 | -0.53489457 | 1.29E-16 |
| Tdrd9 | ENSMUSG00000054003 | -0.530602896 | 7.31E-05 |
| Tead3 | ENSMUSG00000002249 | -0.596589561 | 1.48E-07 |
| Tle2 | ENSMUSG00000034771 | -0.683937971 | 3.34E-07 |
| Tmem106c | ENSMUSG00000052369 | -0.50290518 | 1.02E-07 |
| Tmem150a | ENSMUSG00000055912 | -0.568757504 | 1.16E-06 |
| Tmem255b | ENSMUSG00000038457 | -0.576245863 | 0.01420635 |
| Tpm3-rs7 | ENSMUSG00000058126 | -5.885759703 | 8.49E-93 |
| Trib1 | ENSMUSG00000032501 | -1.207504839 | 2.64E-17 |
| Trim12c | ENSMUSG00000057143 | -1.122692805 | 0.0001936 |
| U2af1l4 | ENSMUSG00000109378 | -1.052960924 | 1.47E-06 |
| Uckl1 | ENSMUSG00000089917 | -0.531808978 | 8.47E-07 |
| Wisp1 | ENSMUSG00000005124 | -0.546254279 | 1.42E-05 |
| Zfp97 | ENSMUSG00000095990 | -1.02956866 | 1.16E-05 |

**Supplementary data sheet 4**

#### Differentially expressed ECM genes (P10 *Apcdd1* -/- vs WT)

##### Significantly upregulated genes

| Gene_name | Gene | Chromosome | log2FoldChange | padj |
|---|---|---|---|---|
| Fras1 | ENSMUSG00000034687.5 | chr5 | 0.500390861 | 7.06E-09 |
| Fbn1 | ENSMUSG00000027204.10 | chr2 | 0.502610325 | 2.84E-06 |
| Sspn | ENSMUSG00000030255.10 | chr6 | 0.518189587 | 3.58E-06 |
| Lama2 | ENSMUSG00000019899.12 | chr10 | 0.523332443 | 5.57E-05 |
| Itga2 | ENSMUSG00000015533.8 | chr13 | 0.574918927 | 3.19E-04 |
| Itga6 | ENSMUSG00000027111.12 | chr2 | 0.595201244 | 2.48E-11 |
| Ccl28 | ENSMUSG00000074715.2 | chr13 | 0.695051602 | 9.21E-06 |
| Tectb | ENSMUSG00000024979.10 | chr19 | 0.869376405 | 4.22E-08 |
| Frem2 | ENSMUSG00000037016.8 | chr3 | 0.899741099 | 6.28E-19 |

#### Differentially expressed ECM genes (P10 *Apcdd1* -/- vs WT)

#### Significantly downregulated genes

| Gene_name | Gene | Chromosome | log2FoldChange | padj |
|---|---|---|---|---|
| Cthrc1 | ENSMUSG00000054196.6 | chr15 | -0.505566406 | 0.001859744 |
| Adamts10 | ENSMUSG00000024299.13 | chr17 | -0.527822837 | 2.46E-10 |
| Gpc2 | ENSMUSG00000029510.12 | chr5 | -0.53040727 | 0.00318171 |
| Podn | ENSMUSG00000028600.12 | chr4 | -0.537021417 | 1.47E-05 |
| Postn | ENSMUSG00000027750.13 | chr3 | -0.543828886 | 2.39E-04 |
| Olfml2a | ENSMUSG00000046618.7 | chr2 | -0.558525393 | 0.002024602 |
| Ctss | ENSMUSG00000038642.7 | chr3 | -0.582298919 | 3.88E-09 |
| Smoc2 | ENSMUSG00000023886.9 | chr17 | -0.610068581 | 1.44E-06 |
| Adamts19 | ENSMUSG00000053441.4 | chr18 | -0.638591756 | 2.06E-04 |
| Ltbp4 | ENSMUSG00000040488.13 | chr7 | -0.650522532 | 3.94E-12 |
| Col17a1 | ENSMUSG00000025064.11 | chr19 | -0.662756196 | 8.40E-05 |
| Col20a1 | ENSMUSG00000016356.14 | chr2 | -0.681575818 | 5.80E-05 |
| Col24a1 | ENSMUSG00000028197.4 | chr3 | -0.705233479 | 1.50E-05 |
| Hist2h2bb | ENSMUSG00000105827.1 | chr3 | -0.723171343 | 5.25E-06 |
| Adamts7 | ENSMUSG00000032363.12 | chr9 | -0.897427501 | 6.58E-15 |
| Hist2h2bb | ENSMUSG00000050936.5 | chr3 | -1.290682591 | 5.69E-17 |

**Differentially expressed ECM genes (P14 *Apcdd1* -/- vs WT)**

#### Significantly upregulated genes

| Gene.name | ID | log2FoldChange | padj |
|---|---|---|---|
| Nov | ENSMUSG00000037362 | 3.312603856 | 2.75E-10 |
| Oscar | ENSMUSG00000054594 | 2.693880227 | 1.05E-05 |
| Spp1 | ENSMUSG00000029304 | 2.415034388 | 0.00525567 |
| Cdh1 | ENSMUSG00000000303 | 1.846869256 | 0.0002214 |
| Tinagl1 | ENSMUSG00000028776 | 1.794251824 | 1.48E-10 |
| Mfap4 | ENSMUSG00000042436 | 1.624074261 | 0.00057869 |
| Ltbp2 | ENSMUSG00000002020 | 1.588659515 | 0.0001187 |
| Optc | ENSMUSG00000010311 | 1.451802056 | 0.00459598 |
| Abi3bp | ENSMUSG00000035258 | 1.4084498 | 2.72E-06 |
| Fbln1 | ENSMUSG00000006369 | 1.382214811 | 0.00788349 |
| Fmod | ENSMUSG00000041559 | 1.306987968 | 0.00155253 |
| Col4a6 | ENSMUSG00000031273 | 1.283940675 | 0.00198635 |
| Frem2 | ENSMUSG00000037016 | 1.279804121 | 1.59E-07 |
| Fbln2 | ENSMUSG00000064080 | 1.269181266 | 4.85E-09 |
| Slit3 | ENSMUSG00000056427 | 1.226470085 | 0.00217061 |
| Vcan | ENSMUSG00000021614 | 1.222609123 | 0.00673527 |
| Col6a3 | ENSMUSG00000048126 | 1.165729253 | 0.00155246 |
| Cpz | ENSMUSG00000036596 | 1.161286685 | 8.27E-05 |
| Serpine1 | ENSMUSG00000037411 | 1.124201947 | 1.73E-07 |
| Ccl28 | ENSMUSG00000074715 | 1.086391655 | 1.08E-06 |
| Serpinf1 | ENSMUSG00000000753 | 1.067388633 | 1.65E-10 |
| Ogn | ENSMUSG00000021390 | 1.047494775 | 2.03E-05 |
| Tnxb | ENSMUSG00000033327 | 1.042242071 | 2.26E-05 |
| Pcolce | ENSMUSG00000029718 | 1.038902907 | 5.17E-08 |
| Lama2 | ENSMUSG00000019899 | 1.024020241 | 1.41E-05 |
| Col10a1 | ENSMUSG00000039462 | 1.019678078 | 0.00447051 |
| Hspg2 | ENSMUSG00000028763 | 0.991144861 | 0.02626044 |
| Nid1 | ENSMUSG00000005397 | 0.931835878 | 0.00610575 |
| Itga6 | ENSMUSG00000027111 | 0.874797958 | 1.43E-09 |
| Ccdc80 | ENSMUSG00000022665 | 0.872369723 | 5.02E-05 |
| Spon2 | ENSMUSG00000037379 | 0.871073605 | 1.75E-10 |
| Emp2 | ENSMUSG00000022505 | 0.86273157 | 6.51E-11 |
| Ctgf | ENSMUSG00000019997 | 0.861809412 | 6.81E-07 |
| Sfrp1 | ENSMUSG00000031548 | 0.854869892 | 0.00551294 |
| Sspn | ENSMUSG00000030255 | 0.850147071 | 6.08E-07 |
| Rarres2 | ENSMUSG00000009281 | 0.843843295 | 6.75E-05 |
| Ltbp1 | ENSMUSG00000001870 | 0.837608829 | 0.00043299 |
| Lgals3 | ENSMUSG00000050335 | 0.832827307 | 2.20E-08 |
| Rhod | ENSMUSG00000041845 | 0.817197662 | 1.31E-07 |
| Col18a1 | ENSMUSG00000001435 | 0.805157116 | 0.0404106 |

| | | | |
|---|---|---|---|
| Igfbp7 | ENSMUSG00000036256 | 0.792993356 | 4.05E-05 |
| Cd44 | ENSMUSG00000005087 | 0.789963393 | 1.08E-05 |
| Col4a5 | ENSMUSG00000031274 | 0.759786081 | 0.00018587 |
| Loxl1 | ENSMUSG00000032334 | 0.755923928 | 0.00031677 |
| Bmp4 | ENSMUSG00000021835 | 0.739085424 | 0.00065171 |
| Gpc4 | ENSMUSG00000031119 | 0.711313638 | 1.54E-06 |
| Tectb | ENSMUSG00000024979 | 0.693755946 | 0.03500763 |
| Efemp1 | ENSMUSG00000020467 | 0.682933524 | 1.98E-10 |
| S100a10 | ENSMUSG00000041959 | 0.68257472 | 5.82E-09 |
| Podn | ENSMUSG00000028600 | 0.682271271 | 0.00156611 |
| Pcolce2 | ENSMUSG00000015354 | 0.681615662 | 1.56E-07 |
| Matn2 | ENSMUSG00000022324 | 0.668640936 | 0.00193311 |
| Pxdn | ENSMUSG00000020674 | 0.660743742 | 3.41E-06 |
| Fgfr2 | ENSMUSG00000030849 | 0.644932484 | 0.00478497 |
| Fbln7 | ENSMUSG00000027386 | 0.644634516 | 0.00013735 |
| Hmcn1 | ENSMUSG00000066842 | 0.633256495 | 0.02942465 |
| Mfap2 | ENSMUSG00000060572 | 0.63265134 | 2.16E-05 |
| Col9a1 | ENSMUSG00000026147 | 0.629593123 | 7.79E-07 |
| Mmp2 | ENSMUSG00000031740 | 0.627884766 | 5.10E-06 |
| NA | ENSMUSG00000102306 | 0.594892768 | 0.02560094 |
| Csf1 | ENSMUSG00000014599 | 0.591530538 | 0.00459021 |
| Ctsk | ENSMUSG00000028111 | 0.589596718 | 0.04154892 |
| Apoe | ENSMUSG00000002985 | 0.584954013 | 8.91E-11 |
| Olfml2a | ENSMUSG00000046618 | 0.581403347 | 0.04022057 |
| Ecm1 | ENSMUSG00000028108 | 0.580854177 | 0.0005673 |
| Cst3 | ENSMUSG00000027447 | 0.56866207 | 1.95E-08 |
| Col9a2 | ENSMUSG00000028626 | 0.558834875 | 2.32E-05 |
| Sparc | ENSMUSG00000018593 | 0.534698888 | 2.70E-21 |
| Thbs2 | ENSMUSG00000023885 | 0.530587028 | 0.01705699 |
| Cd63 | ENSMUSG00000025351 | 0.517079516 | 9.67E-10 |
| Anxa2 | ENSMUSG00000032231 | 0.511944245 | 5.15E-07 |
| Prelp | ENSMUSG00000041577 | 0.506744501 | 0.00071721 |

#### Differentially expressed ECM genes (P14 *Apcdd1* -/- vs WT)

#### Significantly downregulated genes

| Gene.name | ID | log2FoldChange | padj |
|---|---|---|---|
| Bcan | ENSMUSG00000004892 | -1.846246177 | 1.64E-24 |
| Lypd5 | ENSMUSG00000030484 | -1.405624335 | 0.00759046 |
| Col20a1 | ENSMUSG00000016356 | -1.3512778 | 4.47E-30 |
| Itga11 | ENSMUSG00000032243 | -1.247250916 | 0.01880939 |
| Tecta | ENSMUSG00000037705 | -1.093762806 | 4.16E-05 |
| Lama3 | ENSMUSG00000024421 | -1.044360535 | 1.33E-10 |
| Col11a2 | ENSMUSG00000024330 | -1.039438789 | 7.89E-20 |
| Col27a1 | ENSMUSG00000045672 | -0.838194001 | 1.11E-13 |
| Col8a2 | ENSMUSG00000056174 | -0.819810212 | 0.01632307 |
| Col6a1 | ENSMUSG00000001119 | -0.816607391 | 3.13E-09 |
| Wisp3 | ENSMUSG00000062074 | -0.77802675 | 0.02061993 |
| Gpc2 | ENSMUSG00000029510 | -0.746820733 | 6.88E-05 |
| Impg2 | ENSMUSG00000035270 | -0.670192445 | 1.23E-11 |
| Papln | ENSMUSG00000021223 | -0.60429309 | 0.00733438 |
| Sh3pxd2b | ENSMUSG00000040711 | -0.577449901 | 0.0014947 |
| Wisp1 | ENSMUSG00000005124 | -0.546254279 | 1.42E-05 |

Supplementary data sheet 5

### Differentially expressed astrocyte (AC) maturity genes in the P10 Apcdd1-/- retina

#### Significantly upregulated genes (mature astrocyte marker)

| Gene_name | ID | log2FoldChange | Padj |
|---|---|---|---|
| *Plcd4* | ENSMUSG00000026173.12 | 1.743047481 | 1.15E-76 |
| *Aqp4* | ENSMUSG00000024411.9 | 0.632105433 | 2.05E-11 |

**Differentially expressed astrocyte (AC) maturity genes in the P14 Apcdd1-/- retin[illegible]**

**Significantly upregulated genes (mature astrocyte marker)**

| Gene_name | ID | log2FoldChange | Padj |
|---|---|---|---|
| *Gfap* | ENSMUSG00000020932 | 1.276516329 | 0.01551296 |
| *Gja1* | ENSMUSG00000050953 | 0.901632998 | 0.01521701 |
| *Plcd4* | ENSMUSG00000026173 | 0.786165265 | 1.29E-11 |
| *Mlc1* | ENSMUSG00000035805 | 0.614044318 | 1.18E-14 |
| *Apoe* | ENSMUSG00000002985 | 0.584954013 | 8.91E-11 |
| *Aqp4* | ENSMUSG00000024411 | 0.542527436 | 5.67E-05 |

**Significantly downregulated genes (immature astrocyte marker)**

| Gene_name | ID | log2FoldChange | Padj |
|---|---|---|---|
| *Sh3pxd2b* | ENSMUSG00000040711 | -0.577449901 | 0.0014947 |
